# Analysis of a “Split-and-Stuttering” Module of an Assembly Line Polyketide Synthase

**DOI:** 10.1101/2021.01.15.426894

**Authors:** Katarina M. Guzman, Kai P. Yuet, Stephen R. Lynch, Corey W. Liu, Chaitan Khosla

**Author notes:** **Corresponding Author Chaitan Khosla** - Departments of Chemical Engineering, Chemistry, and Stanford Chem-H, Stanford University, Stanford, California 94305, United States. **Author Contributions**. S.R.L. and C.W.L. contributed equally.

## Abstract

Notwithstanding the “one-module-one-elongation-cycle” paradigm of assembly line polyketide synthases (PKSs), some PKSs harbor modules that iteratively elongate their substrates through a defined number of cycles. While some insights into module iteration, also referred to as “stuttering”, have been derived through *in vivo* and *in vitro* analysis of a few PKS modules, a general understanding of the mechanistic principles underlying module iteration remains elusive. This report serves as the first interrogation of a stuttering module from a *trans*-AT subfamily PKS that is also naturally split across two polypeptides. Previous work has shown that Module 5 of the NOCAP (**<u>noc</u>**ardiosis **<u>a</u>**ssociated **<u>p</u>**olyketide) synthase iterates precisely three times in the biosynthesis of its polyketide product, resulting in an all *trans*-configured triene moiety in the polyketide product. Here we describe the intrinsic catalytic properties of this NOCAP synthase module. Through complementary experiments *in vitro* and in *E. coli*, the “split-and-stuttering” module was shown to catalyze up to five elongation cycles, although its dehydratase domain ceased to function after three cycles. Unexpectedly, the central olefinic group of this truncated product had a *cis* configuration. Our findings set the stage for further in-depth analysis of a structurally and functionally unusual PKS module with contextual biosynthetic plasticity.

**TOC/Abstract Graphic:** 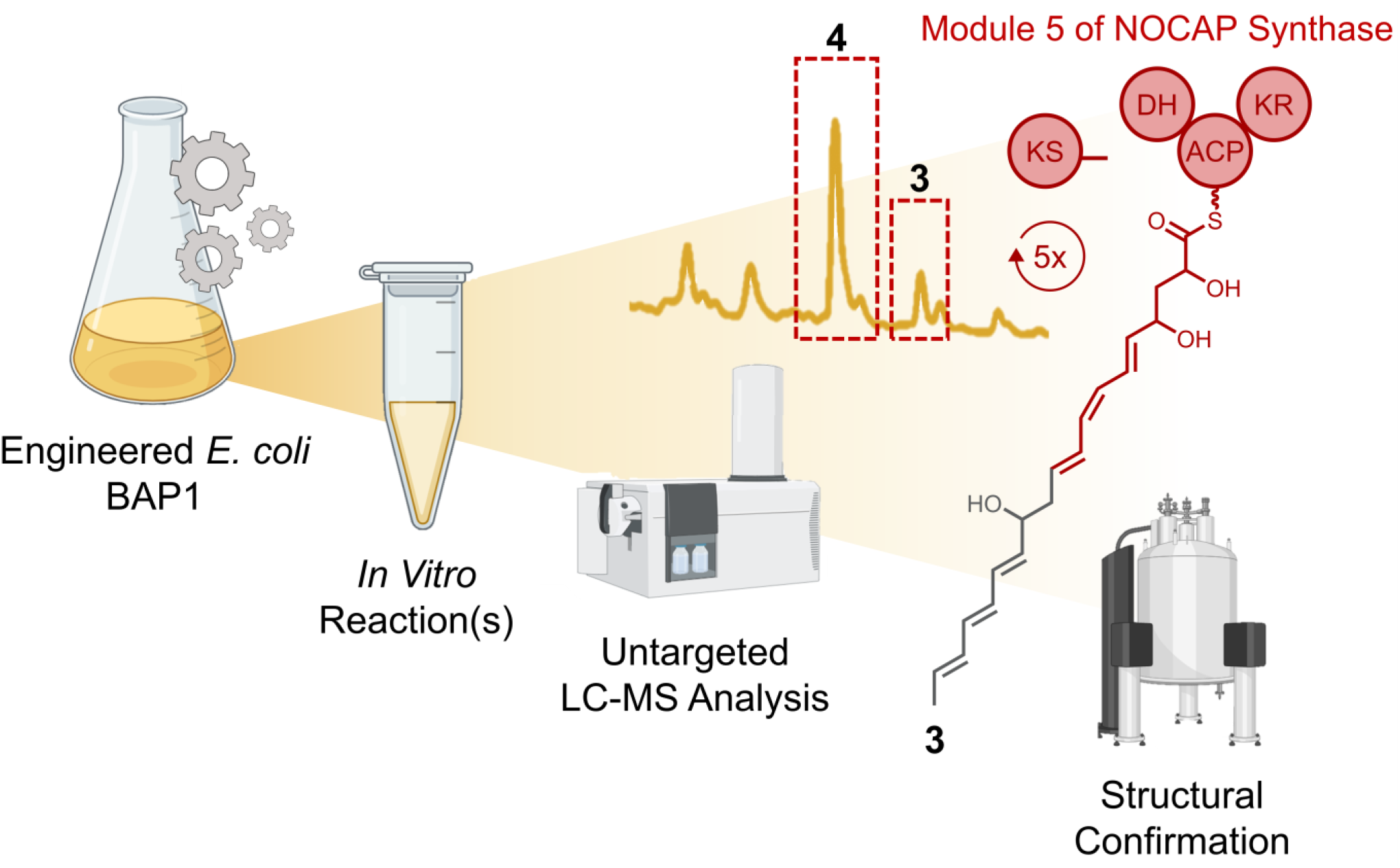

## Main

Polyketide synthases (PKSs) are multifunctional enzymes responsible for the biosynthesis of structurally diverse polyketide products, many of which are clinically used as medicines (1). Therefore, understanding their catalytic diversity and mechanisms has been of high interest to researchers in the hope of engineering novel pharmacologically useful molecules. One class of PKSs, called assembly line PKSs, consists of multiple catalytic modules each of which is responsible for a well-defined set of chemical transformations on the growing polyketide chain. Minimally a PKS module harbors a ketosynthase (KS), acyltransferase (AT), and acyl carrier protein (ACP) domain (2). Some PKS modules (hereafter designated *cis*-AT PKSs) include the AT as an integral domain of a multifunctional polypeptide, whereas other modules (designated *trans*-AT PKSs) engage stand-alone AT proteins (3).

Canonically, each module of an assembly line PKS catalyzes a single elongation and modification cycle, where after the growing polyketide chain is either translocated onto the KS domain of the next module or it is off-loaded from the assembly line (4, 5). Such a collinear architecture has facilitated prediction of structural features of the resulting polyketide products (6). However, not all PKSs operate according to a collinear one-module-one-elongation-cycle model (3). For example, the NOCAP synthase (7, 8) harbors a *trans*-AT module (Module 5) that catalyzes three successive elongation and modification cycles on the growing polyketide chain yielding products **1** and **2** (**Figure 1**) (9,10). After each cycle, the elongated and modified polyketide product is passed from the ACP domain of Module 5 back to its own KS domain (11). This type of programmed module iteration, also known as “stuttering” (12, 13), has been identified in a few other assembly line PKSs (14) such as the stigmatellin (15), lankacidin (16), borrelidin (17), neoaureothin and aureothin (18) PKS pathways.

**Figure 1.**
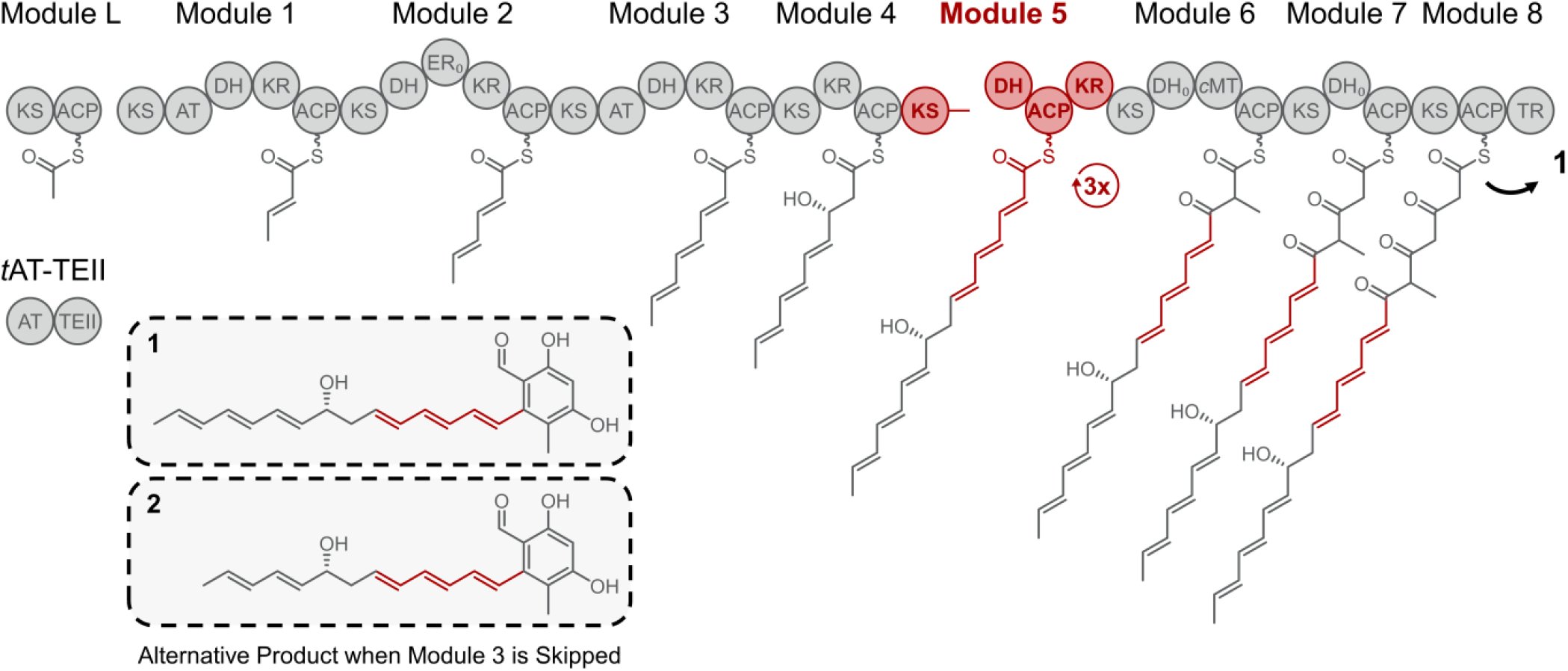
Biosynthetic pathway of **1** *via* Modules L-8-TR + *t*AT-TEII of NOCAP synthase. **2** is an alternative product generated when Module 3 is skipped. Key: KS, ketosynthase; AT, acyltransferase; DH, dehydratase; KR, ketoreductase; ER, enoylreductase; *c*MT, *C*-methyltransferase; ACP, acyl carrier protein; TR, thioester reductase; and TEII, thioesterase II. _0_; subscript implies inactive domain.

While the governing principles that control programmed iteration are not well understood, some insights have been gained through *in vitro* and *in vivo* analysis of *cis*-AT PKSs. Module 1 of the aureothin synthase (*cis*-AT PKS), which catalyzes two elongation and modification cycles in the context of the complete assembly line PKS, was expressed in *Streptomyces lividans* (19). The resulting strain produced a spectrum of compounds that had undergone up to 4 rounds of elongation, although products resulting from 3 and 4 rounds of iteration were orders of magnitude less abundant. Additionally, an *in vitro* analysis of Module 5 of the borrelidin PKS revealed that the KS domain of this stuttering module accepted non-native substrates corresponding to the chain lengths of its natural intermediates (20). Collectively, these and other findings have led to a mechanistic model in which the KS specificity at least partly dictates the number of iterations observed. However, the downstream module also has been suggested to exhibit a gatekeeping role (21). Given the lack of prior studies on stuttering modules from *trans*-AT PKSs, we interrogated Module 5 of the NOCAP synthase. Of particular interest to us was its additionally unusual “split-and-stuttering” feature, with its KS and ACP domains distributed across separate proteins (**Figure 1**).

In order to investigate the potential gatekeeping role of downstream modules in the NOCAP synthase we sought to develop a truncated assembly line excluding Modules 6-8-TR. Fortunately, a system for functionally expressing the entire NOCAP synthase in *E. coli* BAP1 (22) was recently developed which was adapted for the present study (10). Three plasmids with unique antibiotic resistance markers were introduced into *E. coli* BAP1 (**Figure 2, Figure S1**). Plasmid pCK-KPY178 encodes the malonyl-CoA synthetase MatB from *Streptomyces coelicolor* (23), the *Rhizobium leguminosarum* malonate carrier protein MatC (24), the loading module (Module L) and *trans*-AT-TEII (*t*AT-TEII) of the NOCAP synthase. Plasmid pCK-KPY222 encodes Modules 1-3 with engineered docking domains to facilitate protein-protein interactions (25). Finally, pKMG14 or alternatively pKMG15 encodes Module 4 and the DH-ACP-KR or the DH-ACP domains of Module 5, respectively. In the absence of a specific product release mechanism in these truncated assembly lines, we relied on spontaneous hydrolysis of polyketide products from ACP_5_.

**Figure 2.**
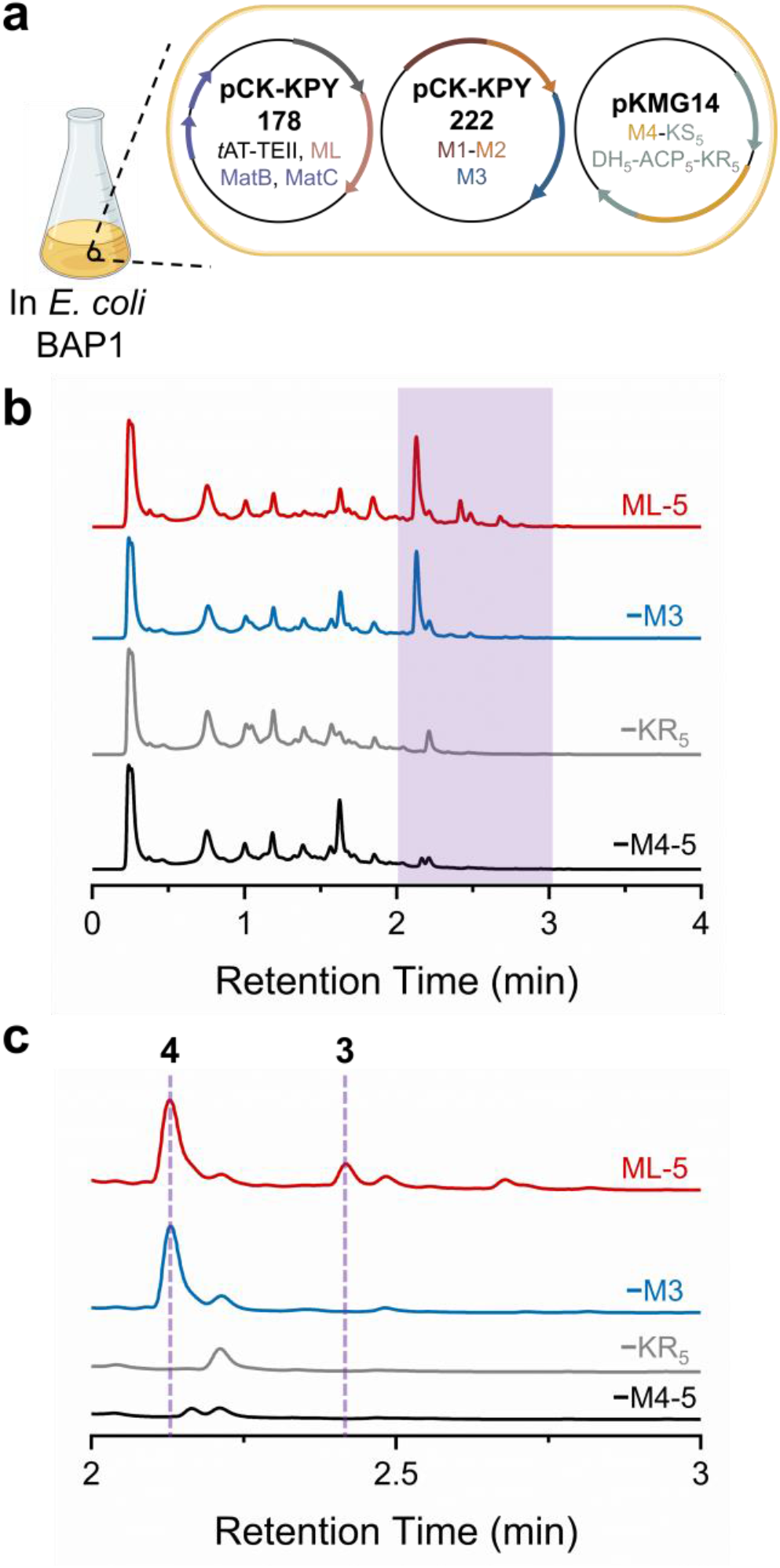
Compounds **3** and **4** identified in (a) an *E. coli* BAP1 strain housing NOCAP synthase Modules L-5 [pCK-KPY178/pCK-KPY222/pKMG14]. (b) HPLC UV chromatogram (at 280 ± 10 nm) comparing crude extracts from *E. coli* BAP1 supernatants. *E. coli* BAP1 strains: [pCK-KPY178/pCK-KPY222] (black, omit Module 4-KS_5_ and DH_5_-ACP_5_-KR_5_); [pCK-KPY178/pCK-KPY222/pKMG15] (grey, omit KR_5_); [pCK-KPY178/pCK-KPY102/pKMG14] (blue, omit Module 3); and [pCK-KPY178/pCK-KPY222/pKMG14] (red, Modules L through 5). (c) Enlarged region of HPLC UV trace for clarity. Data acquired on an Agilent 6545 Q-TOF LC-MS system.

One to three liters of *E. coli* BAP1 [pCK-KPY178/pCK-KPY222/pKMG14] supernatant was subjected to solid phase extraction (SPE) followed by high performance liquid chromatography (HPLC) (**Figure S2**). Encouragingly, two compounds **3** and **4** with distinct UV signatures were identified (**Figure 2, Figures S3-S4**). Control strains harboring pKMG15 in lieu of pKMG14 or lacking either plasmid failed to produce either **3** or **4** (**Figure S5**). High-resolution mass spectrometry (HRMS) analysis revealed that **3** had a molecular formula of C_20_H_28_O_5_ (*m/z*: [M−H]^-^ calcd. 347.1864, obsd. 347.1845, 5.5 ppm), while **4** had a molecular formula of C_18_H_26_O_5_ (*m/z*: [M−H]^-^calcd. 321.1707, obsd. 321.1704, 0.9 ppm) (**Figure S6**). Similar to the relationship between **1** and **2**, these predicted formulas differed by a mass shift consistent with an unsaturated two-carbon (C_2_H_2_) moiety. To verify the anticipated biosynthetic relationship between **3** and **4**, an *E. coli* BAP1 strain lacking Module 3 [pCK-KPY178/pCK-KPY102/pKMG14] was engineered (**SI Methods**). The resulting strain yielded **4** but not **3** (**Figure S5**), confirming a requirement of Module 3 for the biosynthesis of the latter compound. However, because these HRMS values were inconsistent with our expectation that Module 5 catalyzed three elongation and modification cycles, we turned to NMR spectroscopy to elucidate the structures of **3** and **4**.

The structures of **3** and **4** were uncovered by extensive 1-D and 2-D NMR analysis, including ^1^H 1D, ^13^C 1D, COSY, HSQC, HMBC, and ROESY experiments (**Figure 3, Figures S7-S31, Table 1**). The structures deduced from NMR analysis suggested that Module 5 catalyzed five elongation cycles in the biosynthesis of these compounds, but that the final two cycles were not accompanied by dehydration of the β-hydroxyacyl intermediate (**Figure 4, Figure S32**). Furthermore, the presence of a cross-peak in the ROESY spectrum implied the existence of a *cis*-olefinic bond between C_8_-C_9_ in contrast to the all-*trans* configuration of **1** and **2** (**Figures S28 and S31**). To our knowledge this is the first reported instance of “imperfect stuttering”, where the programmed sequence of chain modification reactions is not faithfully catalyzed in every iteration. The unexpected chain length and configuration of **3** (and **4**) raised the possibility that endogenous *E. coli* enzymes played a role in its biosynthesis. For instance, FabA from the fatty acid synthase of *E. coli* iteratively catalyzes the dehydration of ACP-bound β-hydroxyacyl intermediates generated during fatty acid biosynthesis. FabA can also reversibly isomerize *trans*-2-decenoyl-ACP to *cis*-3-decenoyl-ACP (26). Therefore, to rule out the possibility of artifactual involvement of host enzymes, we sought to functionally reconstitute the truncated hexamodular derivative of the NOCAP synthase *in vitro* using purified proteins.

**Figure 3.**
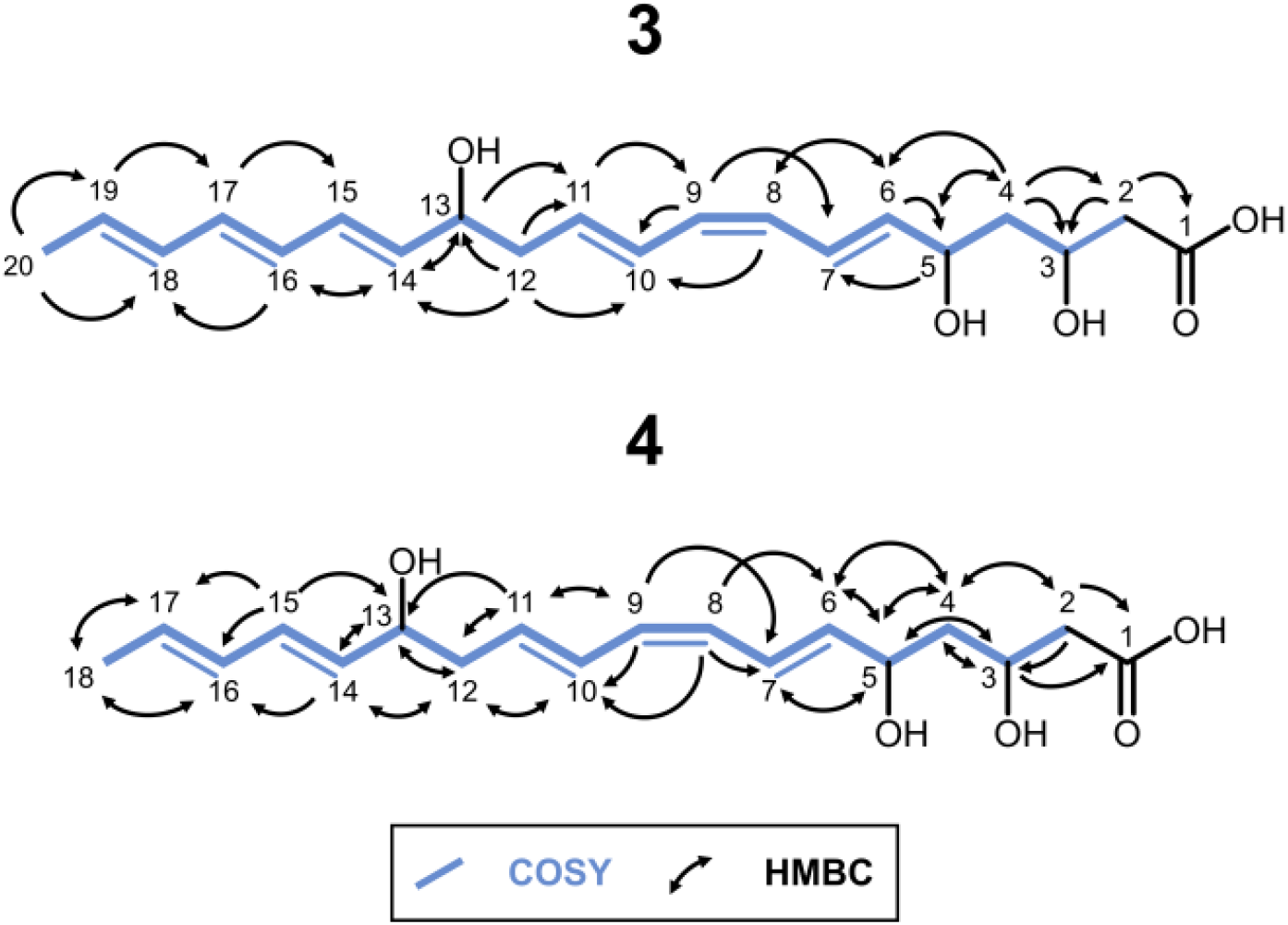
Structures of **3** and **4** assembled from 2D NMR data.

**Figure 4.**
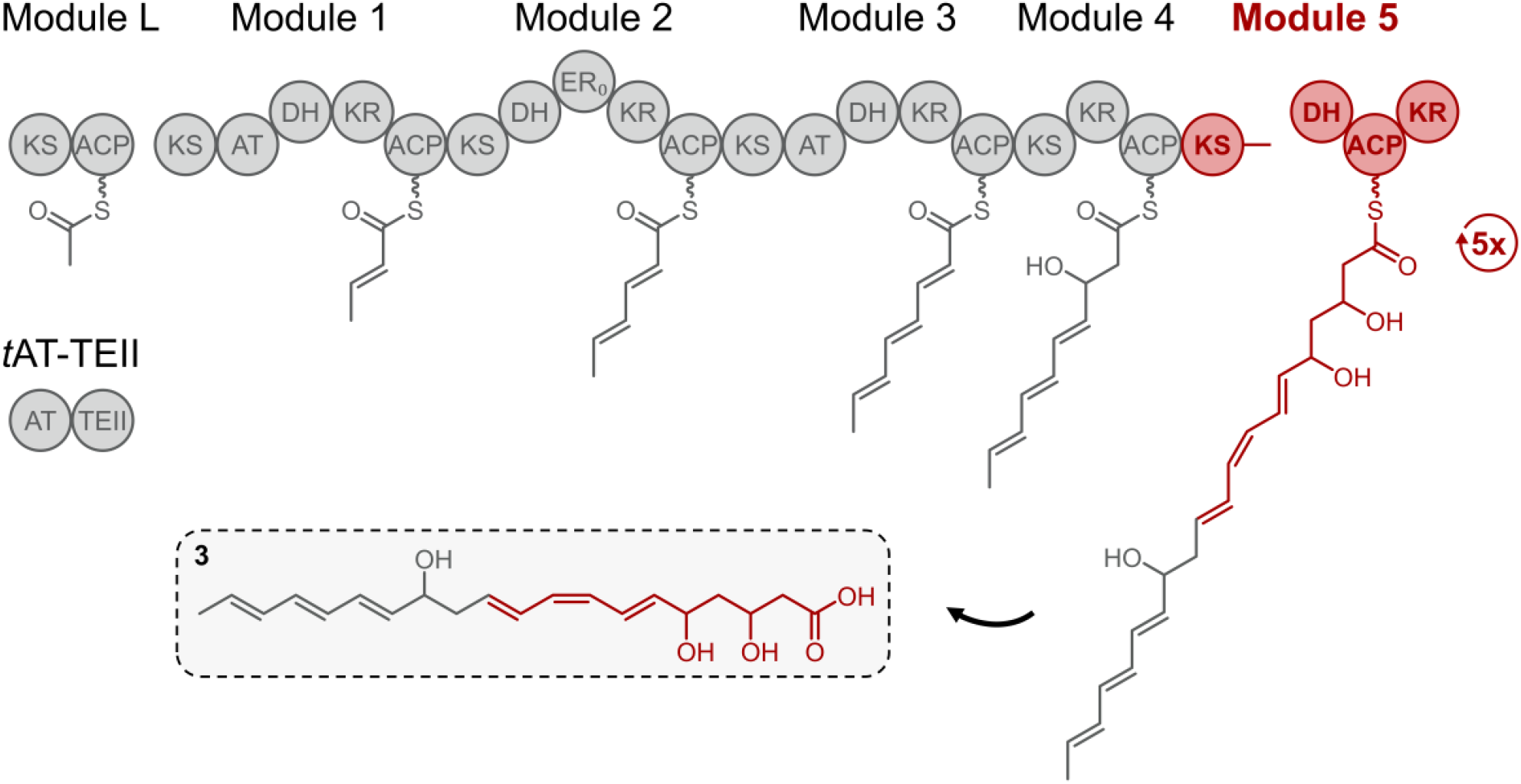
Proposed biosynthesis of **3** from NOCAP synthase Modules L-5 + *t*AT-TEII.

The following proteins were expressed and purified *via* immobilized metal ion chromatography followed by anion exchange chromatography: DH_5_-ACP_5_-KR_5_, DH_5_-ACP_5_ and *t*AT-TEII from the NOCAP synthase, and *S. coelicolor* MatB (**Figure S33, SI Methods**). Additionally, Module L, Modules 1-2, Module 3 and Module 4-KS_5_ were purified on a StrepII-Tag affinity column. Modules L, 1-2, 3, 4-KS_5_ and DH_5_-ACP_5_-KR_5_ were incubated in the presence of *t*AT-TEII, MatB, NADPH, *S*-adenosyl methionine (SAM), malonate, ATP, and CoASH. Parallel reactions were performed without malonate or with [2-^13^C]-, [1,3-^13^C_2_]-, or [^13^C_3_]-malonate to ensure the products detected were of polyketide origin. Product formation relied on spontaneous hydrolysis from ACP_5_. The predominant product of the truncated hexamodular PKS *in vitro* was identical to **3**, as judged by HRMS (**Figure 5**). The observation of +10, +10, and +20 mass shifts in reactions containing [2-^13^C]-, [1,3-^13^C_2_]-, and [^13^C_3_]-malonate, respectively, supported our hypothesis that *in vitro*-derived **3** a) originates from the truncated assembly line and b) had undergone five rounds of elongation at Module 5 (**Figure S34**). **4** was not synthesized in detectable amounts *in vitro*. These results largely support our proposed biosynthetic pathway. However, despite considerable effort, these *in vitro* reactions did not yield sufficient material for ROESY verification of **3**’s geometry.

**Figure 5.**
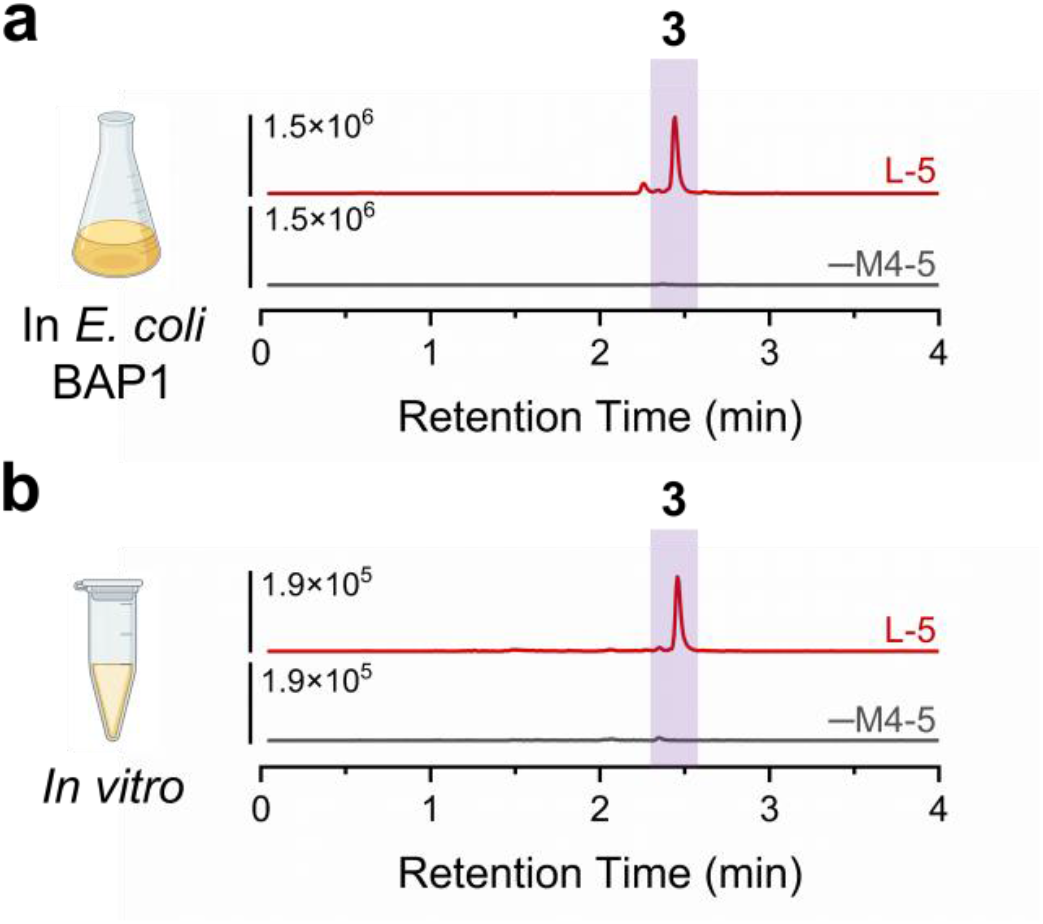
(a) EICs of **3** synthesized in *E. coli* BAP1 and (b) *in vitro* (upper red traces). Negative controls (lower grey traces) omitted Module 4-KS_5_ and DH_5_-ACP_5_-KR_5_.

In summary, this is the first description of a *trans*-AT “split-and-stuttering” module that preferentially iterates for additional cycles beyond those observed in the context of the full PKS assembly line. We therefore propose that Module 6 of the NOCAP synthase regulates the overall polyketide chain length by preferentially recognizing the ACP_5_-bound intermediate after it has undergone three elongation and modification cycles on Module 5. The inability of DH_5_ to catalyze dehydration after three elongation cycles revealed an unprecedented feature of this PKS domain. Future efforts will aim to elucidate the underlying mechanisms that govern module 5’s particularly unusual iterative behavior. Overall, our findings demonstrate the remarkable contextual biosynthetic plasticity of Module 5 of the NOCAP synthase while highlighting the opportunity for future polyketide engineering through a deeper understanding of iterative PKS modules.

## Experimental Section

For detailed descriptions, see the Supporting Information.

### Cloning

Genes were PCR amplified from *Nocardia* genomic DNA or codon optimized for *E. coli* and cloned into pET and Duet vectors using restriction enzyme-based methods, Gibson Assembly, or commercial assembly master mixes. Plasmids were cloned in *E. coli* DH5α, Stellar, or *E. coli* TOP10.

### *E. coli* cultures

*E. coli* BAP1 cultures housing the appropriate plasmids and antibiotics were grown at 30°C until an OD_600_ of 0.2 at which point they were induced with isopropyl-β-D-galactopyranoside (IPTG) and moved to 16°C for an additional 72 hrs.

### Protein Expression and Purification

Proteins were expressed in *E. coli* BAP1 and purified by either Ni-NTA- or Strep-Tactin-based affinity chromatography followed by anion exchange chromatography.

### *In Vitro* Assays and LC-MS

Purified proteins (4-8 µM) were incubated with appropriate cofactors for 24 hrs at 25°C. Reactions (50 µL) were quenched with 50 µL methanol, spun down, and the top-layer (∼50 µL) was analyzed by LC-MS (Agilent 6545 Q-TOF LC-MS system).

### NMR

Spectra were acquired on a 600 MHz Varian Inova spectrometer (Stanford University) and a 900 MHz Bruker NMR spectrometer at the Central California 900 MHz NMR Facility (University of California, Berkeley). Samples were prepared in 250-750 µL of CD_3_OD.

## Associated Content

The Supporting Information is available free of charge at # Detailed methods, figures, and tables (PDF)

## Author Information

### Authors

Complete contact information is available at: #

## Notes

The authors declare no competing financial interest.

## Acknowledgements

We thank members of the Khosla Lab for thoughtful discussions. We also thank Theresa McLaughlin (Stanford University Mass Spectrometry facility) and Jeffrey G. Pelton (University of California, Berkeley) for technical assistance performing LC-MS and NMR experiments respectively. This work was supported by National Institutes of Health (NIH) Grant 5R01 GM087934 (to C.K.) (with extension −26S1 (to C.K and K.M.G.), NIH Grant F32 GM123637 (to K.P.Y.) and a Stanford Graduate Fellowship (SGF) Award (to K.M.G). This work utilized the Stanford Cancer Institute Proteomics/Mass Spectrometry Shared Resource, which is supported by NIH Grant P30 CA124435, and the 900 MHz Bruker NMR spectrometer (funded by NIH Grant P41 GM068933) at the Central California 900 MHz NMR Facility.

