## Supplemental Information for "Analysis of a “Split-and-Stuttering” Module of an Assembly Line Polyketide Synthase"

†Department of Chemical Engineering, ‡Department of Chemistry, §Department of Structural Biology, ^Stanford ChEM-H, Stanford University, Stanford, CA 94305, ⊥Contributed equally

**Table of Contents**

**Materials and Methods**

Expression vectors introduced into *E. coli* BAP1……………………………………………………...1

Expression vectors for *In Vitro* Assays……………………………………………………………...2

Plasmid Sequences……………………………………………………………………………...3

Protein Sequences………………………………………………………………………………...4-11

Biosynthesis of **3** and **4** in *E. coli* BAP1……………………………………………………...12-13

Structural Elucidation of **3** and **4**………………………………………………………………..14-15

*In Vitro* Analysis of Modules L-5, *t*AT-TEII……………………………………………………...16-18

**Supplemental Figures and Table**

**Figure S1**. Plasmids used for biosynthesis of **3** and **4** in *E. coli* BAP1……………………………...19

**Figure S12**. $^{13}$C NMR spectrum (x-axis: δC, 200-0 ppm) of **4** in $CD_3OD$……………………….…..……...31
**Figure S13**. $^{1}$H-$^{1}$H COSY NMR spectrum (positive signals in red; x-axis: δH, 8-1 ppm; y-axis: δH, 8-1 ppm) of **3** in $CD_3OD$……………….…………………………………………………………........…..32
**Figure S14**. Enlarged region of $^{1}$H-$^{1}$H COSY NMR spectrum (positive signals in red; x-axis: δH, 6.8-5.4 ppm; y-axis: δH, 6.8-5.4 ppm) of **3** in $CD_3OD$ (for assignment clarity)……………………….…..33
**Figure S15**. $^{1}$H-$^{1}$H COSY NMR spectrum (positive signals in red; x-axis: δH, 8-1 ppm; y-axis: δH, 8-1 ppm) of **4** in $CD_3OD$………………………………………………………………………...…………34
**Figure S16**. Enlarged region of $^{1}$H-$^{1}$H COSY NMR spectrum (positive signals in red; x-axis: δH, 6.8-5.4 ppm; y-axis: δH, 6.8-5.4 ppm) of **4** in $CD_3OD$ (for assignment clarity)…………………………....35
**Figure S17**. $^{1}$H-$^{13}$C HSQC spectrum (positive signals in red; negative signals in blue; x-axis: δH, 7.4-0 ppm; y-axis: δC, 140-12 ppm) of **3** in $CD_3OD$…………………...……..……………………………...…36
**Figure S18**. Enlarged regions of $^{1}$H-$^{13}$C HSQC spectrum (positive signals in red; negative signals in blue) of **3** in $CD_3OD$……………………………………..…………………………………….……...37
**Figure S19**. $^{1}$H-$^{13}$C HSQC-SE* spectrum (positive signals in red; x-axis: δH, 7.4-0 ppm; y-axis: δC, 140.0-12.0 ppm) of **4** in $CD_3OD$……………………………………………………….……...…38
**Figure S20**. Enlarged regions of $^{1}$H-$^{13}$C HSQC-SE* spectrum (positive signals in red) of **4** in $CD_3OD$... ……………………………………………………………………………………………………...…39
**Figure S21.** $^{1}$H-$^{13}$C HMBC spectrum (positive signals in red; x-axis: δH, 7.2-0 ppm; y-axis: δC, 187-14 ppm) of **3** in $CD_3OD$……………………...…………………………………………………...…40
**Figure S22.** Enlarged regions of $^{1}$H-$^{13}$C HMBC spectrum (positive signals in red) of **3** in $CD_3OD$……..41
**Figure S23.** $^{1}$H-$^{13}$C HMBC spectrum (positive signals in red; x-axis: δH, 7.2-0 ppm; y-axis: δC, 187-14 ppm) of **4** in $CD_3OD$……………………………...………………………........……………...42
**Figure S24.** Enlarged regions of $^{1}$H-$^{13}$C HMBC spectrum (positive signals in red) of **4** in $CD_3OD$…........43
**Figure S25.** Enlarged regions of $^{1}$H-$^{13}$C HMBC spectrum (positive signals in red) of **4** in $CD_3OD$…......44
**Figure S26**. $^{1}$H-$^{1}$H ROESY spectrum (positive signals in red; negative signals in blue; x-axis: δH, 7.2-0.7 ppm; y-axis: δH, 7.2-0.7 ppm) of **3** in $CD_3OD$…………………………………………………...…45
**Figure S27**. Enlarged region of $^{1}$H-$^{1}$H ROESY spectrum (positive signals in red; negative signals in blue; x-axis: δH, 4.5-1.4 ppm; y-axis: δH, 4.5-1.4 ppm) of **3** in $CD_3OD$………………………..……...46
**Figure S28**. Enlarged region of $^{1}$H-$^{1}$H ROESY spectrum (positive signals in red; negative signals in blue; x-axis: δH, 6.9-5.4 ppm; y-axis: δH, 6.9-5.4 ppm) of **3** in $CD_3OD$……………………………………47
**Figure S29**. $^{1}$H-$^{1}$H ROESY spectrum (positive signals in red; negative signals in blue; x-axis: δH, 7.2 -0.7 ppm; y-axis: δH, 7.2-0.7 ppm) of **4** in $CD_3OD$……………………………..…………………...…48
**Figure S30**. Enlarged region of $^{1}$H-$^{1}$H ROESY spectrum (positive signals in red; negative signals in blue; x-axis: δH, 4.5-1.4 ppm; y-axis: δH, 4.5-1.4 ppm) of **4** in $CD_3OD$………………………………...…49
**Figure S31**. Enlarged region of $^{1}$H-$^{1}$H ROESY spectrum (positive signals in red; negative signals in blue; x-axis: δH, 6.9-5.4 ppm; y-axis: δH, 6.9-5.4 ppm) of **4** in $CD_3OD$……………………………………50

**Figure S32**. Biosynthesis of **4** from NOCAP synthase modules L-5 + *t*AT-TEII…….......................…...…51
**Figure S33.** SDS-PAGE of purified proteins used for *in vitro* reactions………………………………...….52
**Figure S34.** Mass spectra (ESI-) of *in vitro*-derived **3** using isotopically labeled malonate…..…...….....53

### I. Materials and Methods

**Expression vectors introduced into *E. coli***

| Plasmid | Vector Backbone | Protein(s) Expressed |
|---|---|---|
| pCK-KPY102 | pCOLADuet-1 (KanR) | *Np* Modules 1-2-DEBS(4)-Strep-tag II* |
| pCK-KPY178 | pCDFDuet-1 (StrR) | **1:** 6×His-*Np* *t*AT-TEII<br>**2:** MBP-*Na* Module L<br>**3:** *S. coelicolor* MatB<br>**4:** *R. leguminosarum* MatC* |
| pCK-KPY222 | pRSFDuet-1 (KanR) | **1:** *Np* Modules 1-2-DEBS(4)*<br>**2:** DEBS(5)-*Na* Module 3-DEBS(2)* |
| pKMG14 | pETDuet-1 (CbR) | **1:** DEBS(3)-*Na* Module 4-$KS_5$*<br>**2:** *Na* 6×His-$DH_5$-$ACP_5$-$KR_5$* |
| pKMG15 | pETDuet-1 (CbR) | **1:** DEBS(3)-*Na* Module 4-$KS_5$*<br>**2:** *Na* 6×His-$DH_5$-$ACP_5$* |

**Key:** *Na, N. araoensis; Np, N. pneumoniae*; DEBS(2), DEBS Module 2 C-terminal docking domain; DEBS(3), DEBS Module 3 N-terminal docking domain; DEBS(4), DEBS Module 4 C-terminal docking domain; DEBS(5), DEBS Module 5 N-terminal docking domain; *, Codon optimized for *E. coli*.

#### Expression vectors for *In Vitro* Assays

| Plasmid | Vector Backbone | Protein(s) Expressed |
|---|---|---|
| pJK52 | pET-28 (KanR) | *Np* *t*AT-TEII-6×His |
| pCK-KPY059 | pET-28 (KanR) | MBP-*Na* Module L-Twin-Strep-tag |
| pCK-KPY102 | pCOLADuet-1 (KanR) | *Np* Modules 1-2-DEBS(4)*-Strep-tag II |
| pCK-KPY293 | pRSFDuet-1 (KanR) | DEBS(5)-*Na* Module 3-DEBS(2)*-Strep-tag II |
| pCK-KPY294 | pETDuet-1 (CbR) | DEBS(3)-*Na* Module 4-$KS_5$*-Strep-tag II |
| pJK65 | pET-21 (CbR) | *Np* $DH_5$-$ACP_5$-$KR_5$-6×His |
| pJK66 | pET-21 (CbR) | *Np* $DH_5$-$ACP_5$-6×His |

**Key:** *Na, N. araoensis; Np, N. pneumoniae*; DEBS(2), DEBS Module 2 C-terminal docking domain; DEBS(3), DEBS Module 3 N-terminal docking domain; DEBS(4), DEBS Module 4 C-terminal docking domain; DEBS(5), DEBS Module 5 N-terminal docking domain; *, Codon optimized for *E. coli*.

The plasmid encoding *S. coelicolor* MatB (pET-28a-$His_6$-MatB.SCo) was a gift from Prof. Michelle Chang (University of California, Berkeley).

#### Plasmid Sequences

| | |
|---|---|
| pCK-KPY102: | https://benchling.com/s/seq-HjngepPFfzmldbpTa458<br>https://openwetware.org/wiki/Khosla:pCK-KPY102 |
| pCK-KPY178: | https://benchling.com/s/seq-m0tUNCCwNsulBo0Evyhj<br>https://openwetware.org/wiki/Khosla:pCK-KPY178 |
| pCK-KPY222: | https://benchling.com/s/seq-QvU91Kd2OXyGqnPOGXhr<br>https://openwetware.org/wiki/Khosla:pCK-KPY222 |
| pKMG14: | https://benchling.com/s/seq-3DSmDngTmIqm7SAlobul<br>https://openwetware.org/wiki/Khosla:pKMG14 |
| pKMG15: | https://benchling.com/s/seq-ycRJ1u3BDGA90haEewsm<br>https://openwetware.org/wiki/Khosla:pKMG15 |
| pJK52: | https://benchling.com/s/seq-Ca009vAMVNw3E1cIoeMA<br>https://openwetware.org/wiki/Khosla:pJK52 |
| pCK-KPY059: | https://benchling.com/s/seq-Dgc3YSpUPe43xLlWo08q<br>https://openwetware.org/wiki/Khosla:pCK-KPY059 |
| pCK-KPY293: | https://benchling.com/s/seq-oOVsdJ5vT3PaF9I7Vi70<br>https://openwetware.org/wiki/Khosla:pCK-KPY293 |
| pCK-KPY294: | https://benchling.com/s/seq-M9ugruYeDvRpw5Q4vrOh<br>https://openwetware.org/wiki/Khosla:pCK-KPY294 |
| pJK65: | https://benchling.com/s/seq-GofcEQRIN8GfTPX9j9vK<br>https://openwetware.org/wiki/Khosla:pJK65 |
| pJK66: | https://benchling.com/s/seq-BHxMVn4Y5t4tlnh69mV5<br>https://openwetware.org/wiki/Khosla:pJK66 |

### Protein Sequences

**pCK-KPY102** (*Np* Modules 1-2-DEBS(4)-Strep-tag II)

MADDGYDLRVLLTRALRRIQELETGPRQEPIAVIGMGCRFPGGAESPQQYWELLSSGRGAIVEVPENRWP
VADSAPRRARRRAGLLAGPIDAMDTEFLGIAPREAASMDPQQRLVLEVAWEAMEDAALAPNGPAAARTGV
FLGVSWQEYQRTITPDWVNSVDAHTLTGTMSSIVAGRVSYVLGLRGPAVAIDTACSSSLVAVHQACRSLR
AGECEVALAGGVNLLQSELTSEALARMGALSPDGHCRPFDARANGYVRGEGAGVVVLKPLSKALADGDPI
RALIRGSSVNHDGRSMGLTAPNPTAQRELLRDALADAGCAADQVGYVETHGTGTPLGDPIEIEALSQVLG
APRADGSVCLLGSVKSQIGHLEAAAGIAGLIKAVLQLEHERIPRQHDFATMNPKIALDGTALAVPTDDVA
WPRTDRPRLAGVSSFGFAGTNAHVLLEQAPEPSPTVDAEQPELLALSARTASALDALSRRYVEQLDTTQS
PLADIAATAALGRSHFAHRRAVVARTVAEAARKLAEPPGRSTDLDSGKPRVAMLFTGQGAQYAGMGAALD
RGYPAFRAALDRCDDILGPLPGGLRLRSVLFEANDEVLSRTEYAQPALFAIEWACAELWATFGVRPDIVL
GHSLGELVAATVAGVLDLETGLRLAAERGRLMQSVARPGAMAAVFGKPEEFAALLDELAAEVAVAAVNGP
GQFVVSGTAEAVRRILEDAARRSSRSVKLPGSTAFHSPLLDPMLDEFETRVAELVPAPRAAVIPLVSNVT
GTRIEAALDAAYWRAHARGTVRFADGVRSVADFGVDAYLEVGPHPVLLEFGRQADPDALWLPSMRRTSTD
DEVLLTGLGEMYCAGADIDWTAVIPPRARRRNGLPTYPFERERITVPIAGASSNTGYLAEHRFQDRAVVP
AAYLALRALRAAGGANRELVDFVAARPVPLEELADPIVSGADGHADVRFHVGNGARTAASGSVRAATSKR
PRDSNDLAEARRASSTEWEVADFYRRCADAGLSFGPRFRWITRLHVTGSSVLAELTRPAELAGEPDAALA
CLLDAAVQTGLALTARHSHAAQLPVGIGRLRVSGGETAAKAWALAERTGRTVRITVFDTAGAVVCEFDDV
WFAESGAAGQSQDRDLLSGLVRVPVWEPAAASATGRGPDATLVIGVGQQADRLAAALRAAGTDAVAVERT
TDPAAWPDDNRAVVYLADTEADDSADTARTETEFVLGVVQAMARAATAGPLYLVTMGATGALPGERVRPA
AATLWGLGAVVRTELPELRCRLIDLPVGARCDDRRLIDELASSAADFVVALRGRQRYVTRTESFDDAGEA
PRLERDAAYLITGGRGALGLTAARTLAECGAGTVVLVGRTDPDAHAQREMARIAALGTRVETATVDVADR
NALSGLLARFGGELPRLAGVVHCAGVLDDAMLIEQTPAHVDRVFAGKVAGAWHLHELTAEHSLELFVLFS
SVSATVGTAGQGNYAAANAYLDALARLRLAEGLPAVSIAWGPWAGTGMAGGLSDAARRAWERRGIAELAA
EDGARALRSLLAAEGVVAVFPAAADASTRQEAVADADEQDMEAPRAAADLLELVRTHAMTALGRSAAVSD
RTPLRELGLDSMMAVELRNALSADLGVRLPATLLFDHPTVAAVSEEIARLSAREGGAPRRTAQEQAEQRP
NTRISEQLTHSMYPTEPPVTPTAPNVSQFTNPESERAADDYADAIAIIGMGCRYPGGVRGPADLWHLLTT
ETDAVTEVPAHRWDVNAYFDPDPDAEGKTYTRWGGFVDGVEEFDAGFFDISVGEARMLDPQQRLLLETSW
EALEHAGYPASRLEGSRTGVFIGLMSNDYAERMARADLAPNAWFGTGNLSSVASGRLSYFFGANGPSLTV
DTACSSSLVTVHTAVRSLRSGECDLALAGGATVVLTPSLDIYFARARGLAADGRCKSFDARADGVVWSDG
AGVLVLKRLADARRDGDRVLGLVRGSAVNSDGRSQGLSAPNGPAQERVIRAALADAAVEAAQIDYVETHG
TGTALGDPIEVNALASVLGDGRTAAAPLWIGSVKSNLGHTQAAAGIANIIKVIESMRHERIPASLHFRSG
NPHIDWDSVAVRVVAEERPWRRSRRPRRAGVSAFGISGTNAHVVIEEAPPVAEAGPVDDSGPHLLVLSAK
STDALRELAARYRDFLAAHPETGLAEVCSSAALGRTHFGERVAITVDTAAQLRAALDNVASGITASGSVA
DLAESGDAGDLARRYVSGAEIDWAKVYSDRPRRHVDLPTYPFQRKRHWLELAARPDATPSSRALRVTPEM
FAGHRLRGVPTVPAAWLCTQVSGLAAQLPGTWVITGLRLVTPLEVTAEATVRVTLDLGSDGYDVRIVHAD
GTSIADAKLAVTEPNSFDGTAEDAATTDADARVWPAYYEKLARLGLHLGADYRCATEFRRISDTEAEAVF
RLPEHDAATLDPILLDACLHTVVLTGASADALFLPSAIDRAQLFGRPTVGDTTVRCRVRVTERGGRVVVG
DATLRTASGRLLARVSGVCLLAANTPVAEPEPHAPEAESSRIAQLLHHVDWTPAADDLPVRGSSAGRWLI
FSDGPVGDAIAAGLAARGAQTVVVRPEDIDDPASVIGMADTSSAHGVVFAWTAVASRTRIDAALRLARAL
DGSGRPPAALWLLTVNAQRVVAADRPDPDQALLWGFGRTLATEHPEFGCRLLDLPEEPTDGALIAEGMLG
AREAGEFAIRDGHWLAARLAAGPAPAEDYRLVAHRSGQFDRLRLESGARRAPGTGEVEIAVAATGLNAHD
IWVASGLHPDAAAALGAECAGRVVAVGAGVAEFAVGDRVVALANGSFARYVTVDHRLVAALPDQLSFAEG
ATVPFAFTTAWHALFDVGELRAGDRVLVHAPADGIGMAAVALAAHAGAQVFTTAALRRWADNAGLPVTQL
DAWRDHELAERVRVATAGRGLDVVVHALGGDLVVHGAEILCPSGRFVDVIDPVQTRVSARHDIDHREFDP
AAVDPQRLGAILRTVMALFEQGALHALPHAVHGIDDAATAFRAMSRREHPGRVVLLGRDAAPVAARRGDR
LTDGTYLVTGGFGGLGAAAARWLAARGAGHLVLVGRRAPDAAGSQLLESLRASGIRVAARQADVSDRAAV
AALLRSIEESGLPPLRGVVHCAGVLDDGVVTEQSWSRFETVLGPKYHAARHLDELTRDQPLDLFVTYSSV
AGVLGTPGQANYAAANAALDALAWRRRAEGLPALSIAWGPFADVGMAADRADITARLTRRGLPPLYANDA

DTALDHLLRLDSATPGVFDFVPQRWYANSSGPVTAGTTIPTATATATPIAGEVSTTPTTVPQSKAPTDSA
PWETIVQRVAMRVLDRASELDPDLPLHDTGVDSLTGMELRAALAAEIGMPLPAGLVFDYPTVRDIARFLQ
RKSFAASPAVDIGDRLDELEKALEALSAEDGHDDVGQRLESLLRRWNSRRADAPSTSAISEDASDDELFS
MLDQRFGGGEDLPNSSSVDKLGSWSHPQFEK

**pCK-KPY178 (1: 6×His-*Np t*AT-TEII)**

MGSSHHHHHHSQDPNSSSMSTEPSECTIVFPGQGAQRTGMGADFCAEFPLARDTFAEASAAVGEDLLRIC
VERDPRLHRTEYTQPCLLTMEIAAYRVLVAEYGVRPVAFGGHSLGEYAALVAAGVFDLADGVRLVRTRGA
LMQRAVPEGEGAMAALILPDIADYGVAELVAAAGAEVANDNSTDQLVISGAADAVSAARAVLAERLPDLR
FVPLRVSAPFHSRWMRGIEAEFAEHLAACAPRAHADRAVAVTSNYTGEFHRPETLAEHLVRQISAPVRWT
DNMRALLRSGTHRYEIGPSAPLSKFFATLGAPVTRIATVGDLRALPIQAGSRDREAKGPAAPLLPATLTQ
TTTQPARATPPSDDTGVLTIHRRTAGTPSLRLFCWPFAGGKAAAYTPWRRHLPDWVELCVIELPARQRHL
AQTPIRRFTDLVDASLAQVLPLTDLPFAFFGHSLGALTAYEVARGLPAGIEPRALFLGAVAAPHLPRPGR
LSGLPDHEFIAAVGHYGGIPPEVRDTPEVMALFLPALRSDFEIFDDYRFAPAAAPSCPAHLFGGRDDRQV
ATTQLEAWADVLPGLRSTELMPGGHFFLVEHREALLGSLTDKLTAVHPDVVPA

**pCK-KPY178 (2: MBP-*Na* Module L)**

MADLNWISAGLMRGSKIEEGKLVIWINGDKGYNGLAEVGKKFEKDTGIKVTVEHPDKLEEKFPQVAATGD
GPDIIFWAHDRFGGYAQSGLLAEITPDKAFQDKLYPFTWDAVRYNGKLIAYPIAVEALSLIYNKDLLPNP
PKTWEEIPALDKELKAKGKSALMFNLQEPYFTWPLIAADGGYAFKYENGKYDIKDVGVDNAGAKAGLTFL
VDLIKNKHMNADTDYSIAEAAFNKGETAMTINGPWAWSNIDTSKVNYGVTVLPTFKGQPSKPFVGVLSAG
INAASPNKELAKEFLENYLLTDEGLEAVNKDKPLGAVALKSYEEELAKDPRIAATMENAQKGEIMPNIPQ
MSAFWYAVRTAVINAASGRQTVDEALKDAQTRITKVPRPPARTSPERKRGFRKTAQVKSPAHVPTPQQDR
QIHRTACCASARSVDRGQVSSVKKTKRHARSRPDRVGAEPIAVIGMSCRFAGGADSPEALWRLLIEETDA
IADEPPPGRWSGDHRGRDRAAAGKTVSVAGGYVADVAGFDAEFFGITPREAADMDPQQRLALELAWEALE
DAAIRPDRVPHTGVYFANKFNDYRAVKLARGATAITPFTSTGDVEGVIANRVSYFLGFDGPSMTVNASCA
GSLVAVHLACQALRAGESALAIAGGVQLNLIPETTIGLSALGVLSARGRSHTFDASADGYVRGEGGAVVV
LKPLRQALRDGDRVYCTLLGSATNNNGRHRSMPASSADGQRQLLRRACARAGVDPSTIDYVEAHGTGTAV
GDRAELSALADVYGTGRAPGAPLIVGSIKTNIGHTEAAAGMAGLVKVALALSHRRIPRSLHFDHAPAELD
PAATGVRVADTEMAWPRHDGPARAAVSAFGFGGSNAHVIAEAAPPPDPVVDSGVSERLVVLSARTEAALR
EHARRLREHLVRDPDIRLGDLAHTLATRTAFQHRISTVADEVDKVVEFLDAYLDGAATDVDRDENAVLGL
VLSAALRPGTVDHRDLLHAAGTLFERGVEPDWAVVNPPGRPVSLPTYPFQRERHWADPQPPGSQAVPVVS
SETAIEAVESFATVRHSVAIASSAGSSEPRRSDAPGVAALAALVTTELAEVVGIAPDRVAARQNFEQLGI
GSVHAVELSARLGARLGIDVPAAVIWGCPTVELLAAELDIRLTPAYSEGVPPSTPASAADPRPTVVEEAL
STAELLALSRQLLGKL

**pCK-KPY178 (3: *S. coelicolor* MatB)**

MGIELLSIGLLIAMFIIATIQPINMGALAFAGAFVLGSMIIGMKTNEIFAGFPSDLFLTLVAVTYLFAIA
QINGTIDWLVECAVRLVRGRIGLIPWVMFLVAAIITGFGALGPAAVAILAPVALSFAVQYRIHPVMMGLM
VIHGAQAGGFSPISIYGGITNQIVAKAGLPFAPTSLFLSSFFFNLAIAVLVFFVFGGARVMKHDPASLGP
LPELHPEGVSASIRGHGGTPAKPIREHAYGTAADTATTLRLNNERITTLIGLTALGIGALVFKFNVGLVA
MTVAVVLALLSPKTQKAAIDKVSWSTVLLIAGIITYVGVMEKAGTVDYVANGISSLGMPLLVALLLCFTG
AIVSAFASSTALLGAIIPLAVPFLLQGHISAIGVVAAIAISTTIVDTSPFSTNGALVVANAPDDSREQVL
RQLLIYSALIAIIGPIVAWLVFVVPGLV

**pCK-KPY178** (**4:** *R. leguminosarum* MatC)

MSSLFPALSPAPTGAPADRPALRFGERSLTYAELAAAAGATAGRIGGAGRVAVWATPAMETGVAVVAALL
AGVAAVPLNPKSGDKELAHILSDSAPSLVLAPPDAELPPALGALERVDVDVRARGAVPEDGADDGDPALV
VYTSGTTGPPKGAVIPRRALATTLDALADAWQWTGEDVLVQGLPLFHVHGLVLGILGPLRRGGSVRHLGR
FSTEGAARELNDGATMLFGVPTMYHRIAETLPADPELAKALAGARLLVSGSAALPVHDHERIAAATGRRV
IERYGMTETLMNTSVRADGEPRAGTVGVPLPGVELRLVEEDGTPIAALDGESVGEIQVRGPNLFTEYLNR
PDATAAAFTEDGFFRTGDMAVRDPDGYVRIVGRKATDLIKSGGYKIGAGEIENALLEHPEVREAAVTGEP
DPDLGERIVAWIVPADPAAPPALGTLADHVAARLAPHKRPRVVRYLDAVPRNDMGKIMKRALNRD

**pCK-KPY222** (**1:** *Np* Modules 1-2-DEBS(4))

MADDGYDLRVLLTRALRRIQELETGPRQEPIAVIGMGCRFPGGAESPQQYWELLSSGRGAIVEVPENRWP
VADSAPRRARRRAGLLAGPIDAMDTEFLGIAPREAASMDPQQRLVLEVAWEAMEDAALAPNGPAAARTGV
FLGVSWQEYQRTITPDWVNSVDAHTLTGTMSSIVAGRVSYVLGLRGPAVAIDTACSSSLVAVHQACRSLR
AGECEVALAGGVNLLQSELTSEALARMGALSPDGHCRPFDARANGYVRGEGAGVVVLKPLSKALADGDPI
RALIRGSSVNHDGRSMGLTAPNPTAQRELLRDALADAGCAADQVGYVETHGTGTPLGDPIEIEALSQVLG
APRADGSVCLLGSVKSQIGHLEAAAGIAGLIKAVLQLEHERIPRQHDFATMNPKIALDGTALAVPTDDVA
WPRTDRPRLAGVSSFGFAGTNAHVLLEQAPEPSPTVDAEQPELLALSARTASALDALSRRYVEQLDTTQS
PLADIAATAALGRSHFAHRRAVVARTVAEAARKLAEPPGRSTDLDSGKPRVAMLFTGQGAQYAGMGAALD
RGYPAFRAALDRCDDILGPLPGGLRLRSVLFEANDEVLSRTEYAQPALFAIEWACAELWATFGVRPDIVL
GHSLGELVAATVAGVLDLETGLRLAAERGRLMQSVARPGAMAAVFGKPEEFAALLDELAAEVAVAAVNGP
GQFVVSGTAEAVRRILEDAARRSSRSVKLPGSTAFHSPLLDPMLDEFETRVAELVPAPRAAVIPLVSNVT
GTRIEAALDAAYWRAHARGTVRFADGVRSVADFGVDAYLEVGPHPVLLEFGRQADPDALWLPSMRRTSTD
DEVLLTGLGEMYCAGADIDWTAVIPPRARRRNGLPTYPFERERITVPIAGASSNTGYLAEHRFQDRAVVP
AAYLALRALRAAGGANRELVDFVAARPVPLEELADPIVSGADGHADVRFHVGNGARTAASGSVRAATSKR
PRDSNDLAEARRASSTEWEVADFYRRCADAGLSFGPRFRWITRLHVTGSSVLAELTRPAELAGEPDAALA
CLLDAAVQTGLALTARHSHAAQLPVGIGRLRVSGGETAAKAWALAERTGRTVRITVFDTAGAVVCEFDDV
WFAESGAAGQSQDRDLLSGLVRVPVWEPAAASATGRGPDATLVIGVGQQADRLAAALRAAGTDAVAVERT
TDPAAWPDDNRAVVYLADTEADDSADTARTETEFVLGVVQAMARAATAGPLYLVTMGATGALPGERVRPA
AATLWGLGAVVRTELPELRCRLIDLPVGARCDDRRLIDELASSAADFVVALRGRQRYVTRTESFDDAGEA
PRLERDAAYLITGGRGALGLTAARTLAECGAGTVVLVGRTDPDAHAQREMARIAALGTRVETATVDVADR
NALSGLLARFGGELPRLAGVVHCAGVLDDAMLIEQTPAHVDRVFAGKVAGAWHLHELTAEHSLELFVLFS
SVSATVGTAGQGNYAAANAYLDALARLRLAEGLPAVSIAWGPWAGTGMAGGLSDAARRAWERRGIAELAA
EDGARALRSLLAAEGVVAVFPAAADASTRQEAVADADEQDMEAPRAAADLLELVRTHAMTALGRSAAVSD
RTPLRELGLDSMMAVELRNALSADLGVRLPATLLFDHPTVAAVSEEIARLSAREGGAPRRTAQEQAEQRP
NTRISEQLTHSMYPTEPPVTPTAPNVSQFTNPESERAADDYADAIAIIGMGCRYPGGVRGPADLWHLLTT
ETDAVTEVPAHRWDVNAYFDPDPDAEGKTYTRWGGFVDGVEEFDAGFFDISVGEARMLDPQQRLLLETSW
EALEHAGYPASRLEGSRTGVFIGLMSNDYAERMARADLAPNAWFGTGNLSSVASGRLSYFFGANGPSLTV
DTACSSSLVTVHTAVRSLRSGECDLALAGGATVVLTPSLDIYFARARGLAADGRCKSFDARADGVVWSDG
AGVLVLKRLADARRDGDRVLGLVRGSAVNSDGRSQGLSAPNGPAQERVIRAALADAAVEAAQIDYVETHG
TGTALGDPIEVNALASVLGDGRTAAAPLWIGSVKSNLGHTQAAAGIANIIKVIESMRHERIPASLHFRSG
NPHIDWDSVAVRVVAEERPWRRSRRPRRAGVSAFGISGTNAHVVIEEAPPVAEAGPVDDSGPHLLVLSAK
STDALRELAARYRDFLAAHPETGLAEVCSSAALGRTHFGERVAITVDTAAQLRAALDNVASGITASGSVA
DLAESGDAGDLARRYVSGAEIDWAKVYSDRPRRHVDLPTYPFQRKRHWLELAARPDATPSSRALRVTPEM
FAGHRLRGVPTVPAAWLCTQVSGLAAQLPGTWVITGLRLVTPLEVTAEATVRVTLDLGSDGYDVRIVHAD
GTSIADAKLAVTEPNSFDGTAEDAATTDADARVWPAYYEKLARLGLHLGADYRCATEFRRISDTEAEAVF
RLPEHDAATLDPILLDACLHTVVLTGASADALFLPSAIDRAQLFGRPTVGDTTVRCRVRVTERGGRVVVG
DATLRTASGRLLARVSGVCLLAANTPVAEPEPHAPEAESSRIAQLLHHVDWTPAADDLPVRGSSAGRWLI
FSDGPVGDAIAAGLAARGAQTVVVRPEDIDDPASVIGMADTSSAHGVVFAWTAVASRTRIDAALRLARAL
DGSGRPPAALWLLTVNAQRVVAADRPDPDQALLWGFGRTLATEHPEFGCRLLDLPEEPTDGALIAEGMLG

AREAGEFAIRDGHWLAARLAAGPAPAEDYRLVAHRSGQFDRLRLESGARRAPGTGEVEIAVAATGLNAHD
IWVASGLHPDAAAALGAECAGRVVAVGAGVAEFAVGDRVVALANGSFARYVTVDHRLVAALPDQLSFAEG
ATVPFAFTTAWHALFDVGELRAGDRVLVHAPADGIGMAAVALAAHAGAQVFTTAALRRWADNAGLPVTQL
DAWRDHELAERVRVATAGRGLDVVVHALGGDLVVHGAEILCPSGRFVDVIDPVQTRVSARHDIDHREFDP
AAVDPQRLGAILRTVMALFEQGALHALPHAVHGIDDAATAFRAMSRREHPGRVVLLGRDAAPVAARRGDR
LTDGTYLVTGGFGGLGAAAARWLAARGAGHLVLVGRRAPDAAGSQLLESLRASGIRVAARQADVSDRAAV
AALLRSIEESGLPPLRGVVHCAGVLDDGVVTEQSWSRFETVLGPKYHAARHLDELTRDQPLDLFVTYSSV
AGVLGTPGQANYAAANAALDALAWRRRAEGLPALSIAWGPFADVGMAADRADITARLTRRGLPPLYANDA
DTALDHLLRLDSATPGVFDFVPQRWYANSSGPVTAGTTIPTATATATPIAGEVSTTPTTVPQSKAPTDSA
PWETIVQRVAMRVLDRASELDPDLPLHDTGVDSLTGMELRAALAAEIGMPLPAGLVFDYPTVRDIARFLQ
RKSFAASPAVDIGDRLDELEKALEALSAEDGHDDVGQRLESLLRRWNSRRADAPSTSAISEDASDDELFS
MLDQRFGGGEDLPNSSSVDKLAAA

**pCK-KPY222** (**2:** DEBS(5)-*Na* Module 3-DEBS(2))

MADLNWISAGRSGDNGMTEEKLRRYLKRTVTELDSVTARLREVEHRAGDPIVIVGMSCRFPGGVRTPEDF
WDLLWHGRDAITEVPPQRWDIDDYYDPDPAALGTMYTRWGGFVDGADEFDPAFFGIGAAEARAMDPQQRL
LLETSWEALERSGRAPGALAGSRTGVFVGLCFSEYPATPIRADDPTAIGAYSVIGSAPSVAAGRLAYVLG
LQGPAMTLDTACSSSLVALQAAADGLRAGRCDLALVGGVNLQLAPETTIGFCRLASLSPDGRSRAFDADA
NGYVRADGCGVLVLTRLSQARRNGDRVLAIVRGVAVNHDGRSNGLTAPNGPAQQRVITDALERAGLRPDA
VDYVETHGTGTPLGDPIEATALAEVYGANRAQPLLIGSVKTNIGHTEAAAGVAGVIKAVLMLRHRRIPPS
LHFHTPNPLIDWAGLPLEVVREARDWPRTEHAPAIGVSSFGMSGTNAHVILAQGDPAPAPRPESPAAQVL
VLSARDETALADIARRWADEIESARAAFGDLAYTAATARTHFEERLAVVADSGAAAARDLRAPIPIRGRV
ERTARAKVALLFTGQGSQYPGMARELFDSEPVFRDALRRCDAVLGPIADGVGLIDAVYGDHGPESLRRTV
FTQPVLFAVEWALWELWRAWGIAPAAVLGHSVGEYVAACAAGAMTVEDALRLVAVRGRLMQRLPDGGGMM
AVDASAEQVAPLVARHEPALSIAGYNGPAQVVVAGPRAALDLLREQLSADGIRSADLTVSHAFHSAAMDP
ILDDLAVAASRATMTDPAIPLISNLTGAPLRPGELGPRYWARHARQPVRFAQGLRAMAELGIDQFLEIGP
HPTLSALGGRVLGDSARFIASLRRGRGDRETMMTALGNLYVGGVAVDWEQVHRPHDRRRVPGPTYPFQRQ
RYWVESHLAAVPPSVDVDAVSEASLRVRRGIAAAADEQEEDPSSPEGTTYFNLTLGPDSPSYLADHVVDR
TVVVPGAWFVATIADIARGLLADDVVAIEDLVITSGLALTGPQPVRIVFEPTGDGFAVHVQTESSGTHAR
AQVRTGTHRLPDGAGLAEILDHCPEHLAPEDFYQRLARAGLPYGPAFRRVVGLRLGRHEALATLSDFDTD
TGTVHPALLDAAFQATYAAMSPRADEQTWVPFAVDQVLASAAVGTVRFGHVRVLEQTPQLCVADVRLLDE
EGSPVLVANGLRVMPAPTPNSAGPSSHERSVAHTYRIDWRETGSAAPASAAGRWLIIGAPGSFADSLAED
LRARGGECVIARPGTGFTFRAANEFEIDLGVPAAVRALLGAVDDGRAPLRGIVSAVALSERNAPAPVAAE
RLVVGALHLVQAAALAGEPAVWLLTRHGQRVEPADRVAPQQSALWGFARSAQLEHRAIGCHVIDIDQVTD
VRSVVDALVTPSAPRQLALRQGRRRVPRLVPLTVVPGEPTATGGTVLITGGLGGVGLRLARWFVDRGRPD
IVLAGRSAPDARAQAVIAELRGRGARIEARSLDVADRAAVAATLDWIDRELPPLAGVAHLAGVLDDGVIA
EQTAARVGRVLAPKTFGAWHLHELTAGRALEFFLLFSSAAAVVGLPGQSGYAAANAFLDGMAESRGASGS
PALSIAWGPWADTGMVERLDSAAVRRLNRRGYGLLRPEHAFDTLDRLIGSGPATVTTMPLTLSGVSDVDV
PDVLLDSVATTVEKDSSTAASADLSARMEALVRRCLGLAPGDDLDRPLHELGLDSVAALEIRDELSRSSG
HRLPATLAFEYPTPRAVVGLLVRLGGTEVRGEAPSALAGLDALEAALPEVPATEREELVQRLERMLAALR
PVAQAADASGTGANPSGDDLGEAGVDELLEALGRELDGD

**pKMG14** (**1:** DEBS(3)-*Na* Module 4-$KS_5$)

MTDSEKVAEYLRRATLDLRAARQRIRELESGDIAIVGVSGRYPGADGLDEYWRVLVEGRDCVTEIPSDRW
DHARYFVPDRHNPFTAYTKWGGFLDDVDCFDPLFFAISPREAEILDPQERLFLQTAWAALEDSGHSRADL
ARGGLRPEQAGVFVGVMWGTYQMFGAEESRLGRGTLPGSTFWSIPNRVSHALDFQGPSMAVDTACSSSLT
ALHLACQSIRTGECRLAVVGGVNLSLHPYKFVALSQGQFASTDGRCRSFGAGGDGYVASEGVGALVLRPL
ADAEADGDTIYGVIKGTAINHGGRVNGFTVPNPDVQAAVIRSALRVGGVDPASVSYVEAHGTGTALGDPI

EIAGLAKVFGSSGVAELPIGSVKSNVGHLEAAAGVAALTKVLLQLRHGTIVPSLHATPPNPNIDFAATPF
RIPTRPLPWPGGDPARPRRATVSSFGAGGANANVVIEEYVDRRPQTPFSGGQVVPVSARDRDRLAEYVAA
LDRFLAGQEPNLADLAYTMQVGRQAGGARAAFVARDIGRLRAAVRAWVDDVPGARSCRDAAEDSRIARWL
DGADIDWATVRDPGPRRRIPAPTYPFARERYWIPDVAPGPQPDAAAGHQSASGADNTWTPQPTSTPGSGP
ITQPTLNGKATLISESIPPQRVSAQSLALEPLSPRPSGGSATKAGLPESASAPRPTAAQRLLVPVWREAP
LPSGDLRSRRALLVYDARTPQHLVEALEAAAGDPGRLVRLSELPGDLHRDDFGAGAALGRALAARYPDID
CVLDVCDLVPAVSGRPTGDLGRMGFYQGVIEARAASLTLLHVTAGHPGADGPESAPLAGLVAALAEEIRD
VRTRSVHVADERDAVCADPTRLLAILGSECAATEDGAPRVRYRGGTREAVELTEATARDGLVVDPARTYV
VTGGTRGLGAAFAAALVERGARRLVILGREPLPPQTEWDRVAAGTGPEAEKVRRLVQLRDRGVAVRTYFG
SLTDETGLSVFFDRVRGELGPIGGVLHCAGSVSTESPAFVRKTGPAVATVLAPKLAGTAVLADVLAADRP
DFFVLFSSVSAVLPGLAVGLSDYAMANAHLDRFAEAQHALGRGWFRSINWPSFRDTGFGEVTSAAYRATG
LPTLSAAEGFELLDAALGLRAPVVLPYHGAALRLSTRTSRPTPVAAPAPTARPAAANGSAAAYEELLRIF
SDELKLAPTRFEGDKRFEEYGADSVLIASAVRRIEQVVEAPFDPALVLEFPTLDALAGLLTEQFPERFAP
EASPQPHPVSAERPRTPDTGTPRPSAATWAADARRGRAAEPVAVVGIGCRLPGGDTPDAFWRLLAAGASG
IREVDPARWDPARFYRPAGGPGLTNSKWGGFVDGVDLFDPEYFGVTDELAWQMDPLQRLLLETSVLATTD
AGYRREELAGKRVGVYAGSRAANYFNRIPVADRHTIIGIGQNFIAARISDYFDWHAGNVVLDSACSSSLL
SVHLACQALRAGEVDAALAGGVEVLLDEMPFVTLSAAGVLSPNGRCATFSEDADGFVPGEGAALLMLKRL
DDALADGDRVYAVLRGSAIGNDGHTMGITTPNMRAQIEVVTAALEVAGMSPDALSYLEAHGTGTMIGDPI
ELKGLATVLGDRSRGPEPCAVGSVKTNIGHLLSAAGIAGAVKTILAVHHAALPPSLHCDNLNPRFNFAGS
PLQVNRELRPWRPADGRPRVAGVSSFGFGGTNAHLVVEQAPDGHRPTRHPLDPPAFRKRSFMLPKTVPST
STEPAPRIVVPISAPAPSFLRVEPLR

**pKMG14** (**2:** *Na* 6×His-$DH_5$-$ACP_5$-$KR_5$)

MGSSHHHHHHSQDPNSSLSGIDLDSFATTLRAEDPVVRDHRVHGVRILPGVVYLDMILRAAMAAGYRPGE
VTIREALFQRPVALADDGAVRIAFRVDRPTGRVAIDVQALDAAGVPVGAVDPVAECVVVLENSNDQPQGI
DVAGFRSAAGARGMAELYAQTGGRGIDHREFMRATGHFARHDDVVTATLGLGAEAAAHLPGFLLHPTFLD
ASTIVPFIDPQRLGVDDDSAFIPIYFGDVRAWRATPEEVFVRAAVRGGGHGGDLLEADVALYAPDGAPVA
RYRLTSKRVRGSADIQRLLTRPEAPSPRQAAPRPSAHAPVASGDPRAVITADLAALVGGELGTDAAGIAT
DVGFYELGLDSGRLLRLTAELEARVGEPLYPTLLFEYQTIDALAGHLAATVGTNYRPAETAPDIEPGESA
VIAETAGGPADPVTAAPATDPVATPEAVVPDDLVVFGFDWESDTTAHPRREPGSLLVFDTAEHPYPQGAS
IVRVRPGGEFVSGSDTFVVDPADARHMELLVAQLTARRLRFDAAVYLWPETTDTDPQTLIERVYLPLRDL
LRAVAGDSEPPMRLVLALTPATPAGELVCALLGGSARTLRLEQPQPLLRAIQRDPARPIADIVATELGID
DRDPATEIRYHGATRQIRRTRLLTGSAPKSGDASLRPGSVVVITGGFGGIGRHLARRLAADYGARLVLVS
RRPLDAPAAGLLAELGDLGAEAVHVRADVSVAADAERVAAIAHATFGPVDVLYHGAGQLRDGLLSGSGRD
DAKPVLAPKIQGVEHLTRALPEARLTVLFSSVSAVLGNPGQADYAAANRYLDVFAEQRDRAGNRTISIGW
PLWAEGGMRPPAEQIAALRRAGVGLLPTATGLDLLLSCATLDRAQLAAVWGRSETLRERLQPEAPPPAVA
PAPVDEHRASVEEPKLAAA

**pKMG15** (**1:** DEBS(3)-*Na* Module 4-$KS_5$)

see **pKMG14** (**1:** DEBS(3)-*Na* Module 4-$KS_5$) sequence

**pKMG15** (**2:** *Na* 6×His-$DH_5$-$ACP_5$)

MGSSHHHHHHSQDPNSSLSGIDLDSFATTLRAEDPVVRDHRVHGVRILPGVVYLDMILRAAMAAGYRPGE
VTIREALFQRPVALADDGAVRIAFRVDRPTGRVAIDVQALDAAGVPVGAVDPVAECVVVLENSNDQPQGI
DVAGFRSAAGARGMAELYAQTGGRGIDHREFMRATGHFARHDDVVTATLGLGAEAAAHLPGFLLHPTFLD
ASTIVPFIDPQRLGVDDDSAFIPIYFGDVRAWRATPEEVFVRAAVRGGGHGGDLLEADVALYAPDGAPVA

RYRLTSKRVRGSADIQRLLTRPEAPSPRQAAPRPSAHAPVASGDPRAVITADLAALVGGELGTDAAGIAT
DVGFYELGLDSGRLLRLTAELEARVGEPLYPTLLFEYQTIDALAGHLAATVKLAAA

**pJK52** (*Np* 6×His-*t*AT-TEII)

MGSSHHHHHHSSGLVPRGSHMSTEPSECTIVFPGQGAQRTGMGADFCAEFPLARDTFAEASAAVGEDLLR
ICVERDPRLHRTEYTQPCLLTMEIAAYRVLVAEYGVRPVAFGGHSLGEYAALVAAGVFDLADGVRLVRTR
GALMQRAVPEGEGAMAALILPDIADYGVAELVAAAGAEVANDNSTDQLVISGAADAVSAARAVLAERLPD
LRFVPLRVSAPFHSRWMRGIEAEFAEHLAACAPRAHADRAVAVTSNYTGEFHRPETLAEHLVRQISAPVR
WTDNMRALLRSGTHRYEIGPSAPLSKFFATLGAPVTRIATVGDLRALPIQAGSRDREAKGPAAPLLPATL
TQTTTQPARATPPSDDTGVLTIHRRTAGTPSLRLFCWPFAGGKAAAYTPWRRHLPDWVELCVIELPARQR
HLAQTPIRRFTDLVDASLAQVLPLTDLPFAFFGHSLGALTAYEVARGLPAGIEPRALFLGAVAAPHLPRP
GRLSGLPDHEFIAAVGHYGGIPPEVRDTPEVMALFLPALRSDFEIFDDYRFAPAAAPSCPAHLFGGRDDR
QVATTQLEAWADVLPGLRSTELMPGGHFFLVEHREALLGSLTDKLTAVHPDVVPA

**pCK-KPY059** (MBP-*Na* Module L-Twin-Strep-tag)

MRGSKIEEGKLVIWINGDKGYNGLAEVGKKFEKDTGIKVTVEHPDKLEEKFPQVAATGDGPDIIFWAHDR
FGGYAQSGLLAEITPDKAFQDKLYPFTWDAVRYNGKLIAYPIAVEALSLIYNKDLLPNPPKTWEEIPALD
KELKAKGKSALMFNLQEPYFTWPLIAADGGYAFKYENGKYDIKDVGVDNAGAKAGLTFLVDLIKNKHMNA
DTDYSIAEAAFNKGETAMTINGPWAWSNIDTSKVNYGVTVLPTFKGQPSKPFVGVLSAGINAASPNKELA
KEFLENYLLTDEGLEAVNKDKPLGAVALKSYEEELAKDPRIAATMENAQKGEIMPNIPQMSAFWYAVRTA
VINAASGRQTVDEALKDAQTRITKVPRPPARTSPERKRGFRKTAQVKSPAHVPTPQQDRQIHRTACCASA
RSVDRGQVSSVKKTKRHARSRPDRVGAEPIAVIGMSCRFAGGADSPEALWRLLIEETDAIADEPPPGRWS
GDHRGRDRAAAGKTVSVAGGYVADVAGFDAEFFGITPREAADMDPQQRLALELAWEALEDAAIRPDRVPH
TGVYFANKFNDYRAVKLARGATAITPFTSTGDVEGVIANRVSYFLGFDGPSMTVNASCAGSLVAVHLACQ
ALRAGESALAIAGGVQLNLIPETTIGLSALGVLSARGRSHTFDASADGYVRGEGGAVVVLKPLRQALRDG
DRVYCTLLGSATNNNGRHRSMPASSADGQRQLLRRACARAGVDPSTIDYVEAHGTGTAVGDRAELSALAD
VYGTGRAPGAPLIVGSIKTNIGHTEAAAGMAGLVKVALALSHRRIPRSLHFDHAPAELDPAATGVRVADT
EMAWPRHDGPARAAVSAFGFGGSNAHVIAEAAPPPDPVVDSGVSERLVVLSARTEAALREHARRLREHLV
RDPDIRLGDLAHTLATRTAFQHRISTVADEVDKVVEFLDAYLDGAATDVDRDENAVLGLVLSAALRPGTV
DHRDLLHAAGTLFERGVEPDWAVVNPPGRPVSLPTYPFQRERHWADPQPPGSQAVPVVSSETAIEAVESF
ATVRHSVAIASSAGSSEPRRSDAPGVAALAALVTTELAEVVGIAPDRVAARQNFEQLGIGSVHAVELSAR
LGARLGIDVPAAVIWGCPTVELLAAELDIRLTPAYSEGVPPSTPASAADPRPTVVEEALSTAELLALSRQ
LLGKLGGGSGGGSGGSAWSHPQFEKGGGSGGGSGGSAWSHPQFEK

**pCK-KPY293** (DEBS(5)-*Na* Module 3-DEBS(2)-StrepII)

MADLNWISAGRSGDNGMTEEKLRRYLKRTVTELDSVTARLREVEHRAGDPIVIVGMSCRFPGGVRTPEDF
WDLLWHGRDAITEVPPQRWDIDDYYDPDPAALGTMYTRWGGFVDGADEFDPAFFGIGAAEARAMDPQQRL
LLETSWEALERSGRAPGALAGSRTGVFVGLCFSEYPATPIRADDPTAIGAYSVIGSAPSVAAGRLAYVLG
LQGPAMTLDTACSSSLVALQAAADGLRAGRCDLALVGGVNLQLAPETTIGFCRLASLSPDGRSRAFDADA
NGYVRADGCGVLVLTRLSQARRNGDRVLAIVRGVAVNHDGRSNGLTAPNGPAQQRVITDALERAGLRPDA
VDYVETHGTGTPLGDPIEATALAEVYGANRAQPLLIGSVKTNIGHTEAAAGVAGVIKAVLMLRHRRIPPS
LHFHTPNPLIDWAGLPLEVVREARDWPRTEHAPAIGVSSFGMSGTNAHVILAQGDPAPAPRPESPAAQVL
VLSARDETALADIARRWADEIESARAAFGDLAYTAATARTHFEERLAVVADSGAAAARDLRAPIPIRGRV
ERTARAKVALLFTGQGSQYPGMARELFDSEPVFRDALRRCDAVLGPIADGVGLIDAVYGDHGPESLRRTV
FTQPVLFAVEWALWELWRAWGIAPAAVLGHSVGEYVAACAAGAMTVEDALRLVAVRGRLMQRLPDGGGMM
AVDASAEQVAPLVARHEPALSIAGYNGPAQVVVAGPRAALDLLREQLSADGIRSADLTVSHAFHSAAMDP
ILDDLAVAASRATMTDPAIPLISNLTGAPLRPGELGPRYWARHARQPVRFAQGLRAMAELGIDQFLEIGP

HPTLSALGGRVLGDSARFIASLRRGRGDRETMMTALGNLYVGGVAVDWEQVHRPHDRRRVPGPTYPFQRQ
RYWVESHLAAVPPSVDVDAVSEASLRVRRGIAAAADEQEEDPSSPEGTTYFNLTLGPDSPSYLADHVVDR
TVVVPGAWFVATIADIARGLLADDVVAIEDLVITSGLALTGPQPVRIVFEPTGDGFAVHVQTESSGTHAR
AQVRTGTHRLPDGAGLAEILDHCPEHLAPEDFYQRLARAGLPYGPAFRRVVGLRLGRHEALATLSDFDTD
TGTVHPALLDAAFQATYAAMSPRADEQTWVPFAVDQVLASAAVGTVRFGHVRVLEQTPQLCVADVRLLDE
EGSPVLVANGLRVMPAPTPNSAGPSSHERSVAHTYRIDWRETGSAAPASAAGRWLIIGAPGSFADSLAED
LRARGGECVIARPGTGFTFRAANEFEIDLGVPAAVRALLGAVDDGRAPLRGIVSAVALSERNAPAPVAAE
RLVVGALHLVQAAALAGEPAVWLLTRHGQRVEPADRVAPQQSALWGFARSAQLEHRAIGCHVIDIDQVTD
VRSVVDALVTPSAPRQLALRQGRRRVPRLVPLTVVPGEPTATGGTVLITGGLGGVGLRLARWFVDRGRPD
IVLAGRSAPDARAQAVIAELRGRGARIEARSLDVADRAAVAATLDWIDRELPPLAGVAHLAGVLDDGVIA
EQTAARVGRVLAPKTFGAWHLHELTAGRALEFFLLFSSAAAVVGLPGQSGYAAANAFLDGMAESRGASGS
PALSIAWGPWADTGMVERLDSAAVRRLNRRGYGLLRPEHAFDTLDRLIGSGPATVTTMPLTLSGVSDVDV
PDVLLDSVATTVEKDSSTAASADLSARMEALVRRCLGLAPGDDLDRPLHELGLDSVAALEIRDELSRSSG
HRLPATLAFEYPTPRAVVGLLVRLGGTEVRGEAPSALAGLDALEAALPEVPATEREELVQRLERMLAALR
PVAQAADASGTGANPSGDDLGEAGVDELLEALGRELDGDGSWSHPQFEK

**pCK-KPY294** (DEBS(3)-*Na* Module 4-$KS_5$-StrepII-tag)

MTDSEKVAEYLRRATLDLRAARQRIRELESGDIAIVGVSGRYPGADGLDEYWRVLVEGRDCVTEIPSDRW
DHARYFVPDRHNPFTAYTKWGGFLDDVDCFDPLFFAISPREAEILDPQERLFLQTAWAALEDSGHSRADL
ARGGLRPEQAGVFVGVMWGTYQMFGAEESRLGRGTLPGSTFWSIPNRVSHALDFQGPSMAVDTACSSSLT
ALHLACQSIRTGECRLAVVGGVNLSLHPYKFVALSQGQFASTDGRCRSFGAGGDGYVASEGVGALVLRPL
ADAEADGDTIYGVIKGTAINHGGRVNGFTVPNPDVQAAVIRSALRVGGVDPASVSYVEAHGTGTALGDPI
EIAGLAKVFGSSGVAELPIGSVKSNVGHLEAAAGVAALTKVLLQLRHGTIVPSLHATPPNPNIDFAATPF
RIPTRPLPWPGGDPARPRRATVSSFGAGGANANVVIEEYVDRRPQTPFSGGQVVPVSARDRDRLAEYVAA
LDRFLAGQEPNLADLAYTMQVGRQAGGARAAFVARDIGRLRAAVRAWVDDVPGARSCRDAAEDSRIARWL
DGADIDWATVRDPGPRRRIPAPTYPFARERYWIPDVAPGPQPDAAAGHQSASGADNTWTPQPTSTPGSGP
ITQPTLNGKATLISESIPPQRVSAQSLALEPLSPRPSGGSATKAGLPESASAPRPTAAQRLLVPVWREAP
LPSGDLRSRRALLVYDARTPQHLVEALEAAAGDPGRLVRLSELPGDLHRDDFGAGAALGRALAARYPDID
CVLDVCDLVPAVSGRPTGDLGRMGFYQGVIEARAASLTLLHVTAGHPGADGPESAPLAGLVAALAEEIRD
VRTRSVHVADERDAVCADPTRLLAILGSECAATEDGAPRVRYRGGTREAVELTEATARDGLVVDPARTYV
VTGGTRGLGAAFAAALVERGARRLVILGREPLPPQTEWDRVAAGTGPEAEKVRRLVQLRDRGVAVRTYFG
SLTDETGLSVFFDRVRGELGPIGGVLHCAGSVSTESPAFVRKTGPAVATVLAPKLAGTAVLADVLAADRP
DFFVLFSSVSAVLPGLAVGLSDYAMANAHLDRFAEAQHALGRGWFRSINWPSFRDTGFGEVTSAAYRATG
LPTLSAAEGFELLDAALGLRAPVVLPYHGAALRLSTRTSRPTPVAAPAPTARPAAANGSAAAYEELLRIF
SDELKLAPTRFEGDKRFEEYGADSVLIASAVRRIEQVVEAPFDPALVLEFPTLDALAGLLTEQFPERFAP
EASPQPHPVSAERPRTPDTGTPRPSAATWAADARRGRAAEPVAVVGIGCRLPGGDTPDAFWRLLAAGASG
IREVDPARWDPARFYRPAGGPGLTNSKWGGFVDGVDLFDPEYFGVTDELAWQMDPLQRLLLETSVLATTD
AGYRREELAGKRVGVYAGSRAANYFNRIPVADRHTIIGIGQNFIAARISDYFDWHAGNVVLDSACSSSLL
SVHLACQALRAGEVDAALAGGVEVLLDEMPFVTLSAAGVLSPNGRCATFSEDADGFVPGEGAALLMLKRL
DDALADGDRVYAVLRGSAIGNDGHTMGITTPNMRAQIEVVTAALEVAGMSPDALSYLEAHGTGTMIGDPI
ELKGLATVLGDRSRGPEPCAVGSVKTNIGHLLSAAGIAGAVKTILAVHHAALPPSLHCDNLNPRFNFAGS
PLQVNRELRPWRPADGRPRVAGVSSFGFGGTNAHLVVEQAPDGHRPTRHPLDPPAFRKRSFMLPKTVPST
STEPAPRIVVPISAPAPSFLRVEPLRGSWSHPQFEK

**pJK65** (*Np* $DH_5$-$ACP_5$-$KR_5$-6×His)

MASRGIDLDSFATTLHADDPVVRDHRVHGVRILPGVVYLDMILRAAMAAGYQPGDLTIREALFQRPVALA
DDGAVRIAFRVDRPTGRIAIDVQRLDHAGAPYGPVDSVAECVVVRSSSPTEPPASDTAVGSAQARGMAEM
YAQSGSRGIEHRDFMRATGHFARSGDIVTATMGLGPEAAAHLPGFLLHPTFLDASTIVPFVDPQRLGADD

DSAFIPIYFGDVHAWQATPEQVFVRASVRGGGHGGDLLEADVALYAPDGSPVVRYRLTSKRVRGSADIHR
LLTPTEAPAPQQPVPASSTVMQYDPAATGPVSAAQYGPPAASPAQGPSTKDPRALITADLAALVAAELGA
DAAGIATDVGFYELGLDSGRLLRLTAELEGRVGEPLYPTLLFEYQTIDALAGHLAETVGANYRPDAATTA
HQPGAGTTPAEFATQSTGLSRAPQVAAPPSGSTDPGPAPHAATPPGSNALGSTPQAVTPASGSTDLGPAP
YAATPTTGSTGQGPAPQAVTLTTGSTDPAPAPQAAVPTTPSTGSVPQAAAPKAGSTEFRSAQSDAPIADP
PFASSTTVAPASAARDELVTFGFDWETDATAHPRREAIGLLVFDTADHPYRHAGAAAVVRVRPGDEFVSG
AVDFAVNPTDPRHMGLLVEQLTARGLRFDAVLYLWPESIGDDDSRDLIERVYLPLRDLVRATAGLSESPV
RLLLAVTRATPAGELSCALLGGFARTLRLEQPQPILRAVQCDPGAAIDDIVAAELGVEERHPATEIRYRG
GRREIRSPRLDTDSAGATGDPVLRRGSVVVVTGGFGGIARHLARRLATDYSARLVLVSRRPLDAAGAGLL
AELNDLGAEAVHVRADVSVAADAERVAAIAHATFGPVDVLYHGAGQLRDGLVSGSGGDEAISVLAPKVHG
VRHLTRALPEARLTVLFSSVSAVLGNPGQADYAAANRYLDVFAEQRDQAGTRTVSVNWPLWAEGGMHPPA
EQIAALRRAGIGLLPTATGLDLLLSCAALDRAQLAAVWGDPEVVRARLRPAASQARAAVESVTADEHPAP
ARPVGVEKLAAALEHHHHHH

**pJK66** (*Np* $DH_5$-$ACP_5$-6×His)

MASRGIDLDSFATTLHADDPVVRDHRVHGVRILPGVVYLDMILRAAMAAGYQPGDLTIREALFQRPVALA
DDGAVRIAFRVDRPTGRIAIDVQRLDHAGAPYGPVDSVAECVVVRSSSPTEPPASDTAVGSAQARGMAEM
YAQSGSRGIEHRDFMRATGHFARSGDIVTATMGLGPEAAAHLPGFLLHPTFLDASTIVPFVDPQRLGADD
DSAFIPIYFGDVHAWQATPEQVFVRASVRGGGHGGDLLEADVALYAPDGSPVVRYRLTSKRVRGSADIHR
LLTPTEAPAPQQPVPASSTVMQYDPAATGPVSAAQYGPPAASPAQGPSTKDPRALITADLAALVAAELGA
DAAGIATDVGFYELGLDSGRLLRLTAELEGRVGEPLYPTLLFEYQTIDALAGHLAETVKLAAALEHHHHH
H

#### Biosynthesis of 3 and 4 in *E. coli*

##### Media

***LB Broth***: 25.0 g/L LB Broth, Miller granulated powder (Thermo Fisher Scientific) (supplemented with 100 mg/L carbenicillin disodium (Gold Biotechnology), 50 mg/L kanamycin monosulfate (Gold Biotechnology) and 50 mg/L streptomycin sulfate (Gold Biotechnology))

Sterilized in an autoclave before addition of antibiotics

***Terrific Broth***: 47.6 g/L Terrific Broth (modified) powder (MilliporeSigma), 10 g/L glycerol (MilliporeSigma) (supplemented with 100 mg/L carbenicillin disodium (Gold Biotechnology), 50 mg/L kanamycin monosulfate (Gold Biotechnology) and 50 mg/L streptomycin sulfate (Gold Biotechnology))

Sterilized by a 0.2 µm PES bottle filter attachment (Thermo Fisher, # 595-4520)

##### Culture(s)

A single colony of *E. coli* BAP1 [pCK-KPY178/pCK-KPY222/pKMG14] was used to inoculate an overnight seed culture of LB Broth (15 mL) grown at 30°C in a 50 mL Falcon tube. The overnight seed culture was pelleted by centrifugation at 4000 × *g* for 15 min at room temperature and re-suspended in LB Broth (15 mL). 12 mL of starter culture were used to inoculate 1 L of Terrific Broth in a 2.5 L Tunair shake flask (IBI Scientific) and agitated at 30°C (200 rpm) until the $OD_{600}$ reached ~0.2. After the addition of 10 mL/L culture of 500 mM sodium malonate, pH 7.4 (MilliporeSigma), 1 mL/L culture of 50 mM calcium D-pantothenate (MilliporeSigma) and 0.5 mL/L culture of 200 mM isopropyl β-D-1-thiogalactopyranoside (IPTG, Gold Biotechnology), the cultures were agitated (200 rpm) at 16°C for an additional 72 hrs.

##### Extraction

***HPLC A***: 99.9% (v/v) water, 0.1% (v/v) formic acid (Thermo Fisher Scientific) HPLC Grade

***HPLC B***: 99.9% (v/v) acetonitrile, 0.1% (v/v) formic acid (Thermo Fisher Scientific) HPLC Grade

1-3 L of culture were emptied into centrifuge bottles. The Tunair shake flask was then rinsed with ~200 mL of distilled $H_2O$ to ensure maximum yield and collection of foam that arose during culturing. Bottles were then centrifuged at 5000 × *g* for 15 min at 16°C. Supernatant and pellets were separated and frozen at -20°C. Pellet and supernatant were processed separately. Compounds **3** and **4** were identified in the supernatant, judged by crude LC-MS (Waters SQ Detector 2 LC-MS system**) extract comparative analysis. With these results the following protocol was adopted for the extraction of **3** and **4**. [1]

A 60-mL Solid-Phase Extraction column (SPE) (Discovery® DSC-18 SPE Tube) was retrofitted onto either a 50 mL Falcon tube with a vacuum syringe line or a 2 L Erlenmeyer flask with an in-house vacuum line depending on the extraction step (see below). The following were gently passed through the SPE column (flow rate ~1 drop/s):

1. 25 mL of HPLC grade Methanol (Fisher Scientific) - **Wash**
2. 25 mL of HPLC A - **Equilibration**
3. 500 mL of *E. coli* supernatant - **Load**
4. 5 × 5 mL HPLC B – **Elution(s)**

**Notes:** The *E. coli* supernatant was filtered using a 0.2 µm PES membrane filtering unit (Thermo Scientific Nalgene Rapid-Flow) prior to loading on the SPE column. 1 SPE column was used/500 mL of *E. coli* supernatant processed. <u>Samples were protected from light at all times.</u>

Elution fractions 1-5 identified by LC-MS** to contain **3** and **4** were pooled and dried under a gentle $N_2$ stream. Once a final volume of 10 mL was achieved, 10 mL of HPLC A was added, mixed and then dried *in vacuo*. The sample was removed from vacuum and dissolved in 2.4 mL 75:25 (% v/v) HPLC A:HPLC B and filtered through a 0.45 µm PTFE membrane (VWR). This mixture was separated with a gradient elution method (70:30 % HPLC A:HPLC B to 45:55 % HPLC A:HPLC B over 1 hr at 1 mL/min) on an Agilent 1260 Infinity LC system equipped with an Agilent Eclipse XDB-C8 column (5 µm, 250 mm x 9.4 mm). Fractions identified by LC-MS (**Figure S2-4**) to contain **3** and **4** were diluted with $D_2O$, applied to a 200 mg C18 solid-phase extraction column (Thermo-Fisher, #60108-303), washed with $D_2O$ to remove HPLC A and HPLC B solvents, and eluted with 0.25-0.75 mL $CD_3OD$ for NMR experiments. Unfortunately, since the compounds were unable to be lyophilized without compromising their quality, the exact quantity (mass) obtained is unknown.

**Compounds were separated with a gradient elution method (98/2 % HPLC A/HPLC B to 5/95 % HPLC A/HPLC B over 4 min at 0.3 mL/min) equipped with an Agilent InfinityLab Poroshell 120 SB-C18 column (2.7 µm, 2.1 x 50 mm) and operating in both positive and negative ion mode.

[1] Wang B, Guo F, Huang C, Zhao H. Unraveling the iterative type I polyketide synthases hidden in *Streptomyces*. *Proc Natl Acad Sci U S A*. **2020**, 117, 8449-8454.

#### Structural Elucidation of 3 and 4

Samples were prepared in 0.25-0.75 mL of methanol-$d_4$ (Cambridge Isotope Laboratories) in standard borosilicate glass NMR tubes (5 mm x 8 inch) (Fisher Scientific) or in matched symmetrical Shigemi tubes (Wilmad Labglass).

NMR spectra of **3** were acquired on a 900 MHz Bruker AVANCE II spectrometer (Central California 900 MHz NMR Facility, California Institute for Quantitative Biosciences, University of California, Berkeley) with a 5 mm TCI (H{CN}, Z-gradient) cryoprobe, running TopSpin v3.2. Sample temperature was regulated at 25°C. Experiments acquired include: $^{1}$H 1D, $^{13}$C 1D, COSY, HSQC, HMBC, and ROESY. This instrument was also used to obtain the $^{13}$C 1D spectra of **4**. The remaining NMR spectra of **4** were acquired on a 600 MHz Varian Inova spectrometer (NMR facility, Department of Chemistry, Stanford University) with a 5 mm (H{CN}, Z-gradient) probe. Sample temperature was regulated at 25°C. Experiments acquired include: $^{1}$H 1D, COSY, HSQC, HMBC, and ROESY. Data files were analyzed using Mnova software. Automatic baseline corrections, phasing, and referencing (when applicable; $CD_3OD$; H 3.31 ppm, C 49.0 ppm) were used to correct offsets in the acquired data. Additionally, t1 noise was reduced using processing in Mnova.

Samples were protected from light at all times.

NMR Experimental Parameters*: Compound **3**

| | **nt** | **(np, ni)** | **sw** | **d1** | **at** |
|---|---|---|---|---|---|
| **$^{1}$H 1D** | 16 | - | 12626.3 | 2.00 | 5.1905 |
| **$^{13}$C 1D** | 9900 | - | 59523.8 | 2.80 | 0.3588 |
| **COSY** | 4 | (4096, 400) | (12626.3, 12594.5) | 1.50 | 0.1622 |
| **HSQC** | 32 | (2048, 128) | (12626.3, 37313.4) | 1.50 | 0.0811 |
| **HMBC** | 64 | (4096, 64) | (12626.3, 54347.8) | 1.50 | 0.1622 |
| **ROESY** | 8 | (4096, 400) | (12626.3, 12594.5) | 2.00 | 0.1622 |
| **(300 ms mixing)** | | | | | |

*Varian Inova/Agilent parameter terminology used

NMR Experimental Parameters*: Compound **4**

| | nt | (np, ni) | sw | d1 | at |
|---|---|---|---|---|---|
| **$^{1}$H 1D** | 1 | - | 5950.6 | 1.00 | 5.5067 |
| **$^{13}$C 1D**** | 1218 | - | 59523.8 | 2.80 | 0.3599 |
| **COSY** | 4 | (2048, 400) | (5950.6, 5950.6) | 1.50 | 0.1721 |
| **HSQC** | 16 | (2048, 200) | (5950.6, 25641.0) | 1.50 | 0.1721 |
| **HMBC** | 80 | (1786, 300) | (5950.6, 36199.1) | 1.50 | 0.1501 |
| **ROESY** | 8 | (1786, 200) | (5950.6, 5950.6) | 3.00 | 0.1501 |
| **(200 ms mixing)** | | | | | |

*Varian Inova/Agilent parameter terminology used; **Acquired on 900 MHz instrument

#### *In Vitro* Analysis of Modules L-5, *t*AT-TEII

##### Media

*LB Broth*: 25.0 g/L LB Broth, Miller granulated powder (Thermo Fisher Scientific)

Supplemented with either 100 mg/L carbenicillin disodium (Gold Biotechnology) or 50 mg/L kanamycin monosulfate (Gold Biotechnology) – see "Expression vectors for *In Vitro* Assays" list

All LB Broth was autoclaved for sterilization

##### Buffer(s)

*Ni-NTA Lysis/Wash Buffer*: 50 mM sodium phosphate (Thermo Fisher Scientific), 500 mM sodium chloride (Thermo Fisher Scientific), 20 mM imidazole (MilliporeSigma), 1 mg/mL chicken egg white lysozyme (Alfa Aesar), 10% (v/v) glycerol (Fisher Scientific), pH 7.4

*Ni-NTA Elution Buffer*: 50 mM sodium phosphate (Thermo Fisher Scientific), 20 mM sodium chloride (Thermo Fisher Scientific), 500 mM imidazole (MilliporeSigma), 10% (v/v) glycerol (Fisher Scientific), pH 7.4

*Strep-Tactin Lysis Buffer*: 50 mM sodium phosphate (Thermo Fisher Scientific), 500 mM sodium chloride (Thermo Fisher Scientific), 10% (v/v) glycerol (Fisher Scientific), 1 mg/mL chicken egg white lysozyme (Alfa Aesar), 4 mg/L chicken egg white avidin (MilliporeSigma), pH 7.4

*Strep-Tactin Wash Buffer*: 50 mM sodium phosphate (Thermo Fisher Scientific), 500 mM sodium chloride (Thermo Fisher Scientific), 10% (v/v) glycerol (Fisher Scientific), pH 7.8

*Strep-Tactin Elution Buffer*: 50 mM sodium phosphate (Thermo Fisher Scientific), 500 mM sodium chloride (Thermo Fisher Scientific), 10% (v/v) glycerol (Fisher Scientific), 5 mM d-desthiobiotin (MilliporeSigma), pH 7.8

*FPLC A*: 50 mM sodium phosphate (Thermo Fisher Scientific), 10% (v/v) glycerol (Fisher Scientific), pH 7.4

*FPLC B*: 50 mM sodium phosphate (Thermo Fisher Scientific), 1 M sodium chloride (Thermo Fisher Scientific), 10% (v/v) glycerol (Fisher Scientific), pH 7.4

##### Culture(s)

A single colony of *E. coli* BAP1 housing the appropriate plasmid was used to inoculate an overnight seed culture of LB Broth (10-80 mL) supplemented with either Kanamycin (50 mg/L) or Carbenicillin (100 mg/L) and grown at 37°C. The overnight seed culture was pelleted by centrifugation at 4000 × *g* for 15 min at room temperature and re-suspended in fresh LB Broth (10-80 mL). 4-5 mL of seed culture was added into each 2.5 L Tunair shake flask (2-8 flasks with 1 L fresh LB Broth in each flask) and agitated (200 rpm) at 37°C until the $OD_{600}$ reached 0.2-0.3. At this point, the incubator temperature was set to 18°C, and the flasks were allowed to shake for 1 hr before the addition of 0.5-1.0 mL/L culture of 200 mM IPTG at an $OD_{600}$ of 0.5-0.7. The cultures were further agitated (180 rpm) at 18°C for an additional 15-17 hrs. Cells were harvested by centrifugation at 5000 × *g* for 15 min at 4°C, frozen in liquid nitrogen and stored at -80°C.

| Protein | Scale (L) | Approximate Yield (mg/L) | IPTG $[\mu M]_{Final}$ |
|---|---|---|---|
| JK52 | 4 | 15 mg/L | 200 |
| CK-KPY059 | 8 | 10 mg/L | 100 |
| CK-KPY102 | 8 | 3 mg/L | 100 |
| CK-KPY293 | 8 | 1 mg/L | 100 |
| CK-KPY294 | 8 | 5 mg/L | 100 |
| JK65 | 4 | 5 mg/L | 200 |
| JK66 | 4 | 10 mg/L | 200 |

#### Protein Purification

Thawed cells were re-suspended either in Ni-NTA Lysis Buffer or Strep-Tactin Lysis Buffer, incubated at 4°C for 1 hr and sonicated. Lysates were clarified twice by centrifugation at 25,000 × *g* for 1 hr at 4°C. Strep-II tagged protein lysates were incubated overnight while rotating end-over-end with 0.5 mL/L culture Strep-Tactin Sepharose resin (IBA Lifesciences). Whereas, 6×His tagged protein lysates were incubated for 1-2 hrs while rotating end-over-end with 2 mL/L culture HisPur Ni-NTA resin (Thermo Fisher Scientific). The protein-bound resin was washed (25 mL×3) with either Ni-NTA Wash Buffer or Strep-Tactin Wash Buffer. Protein was eluted off the resin with either Ni-NTA

Elution Buffer (5-10 mL × 2) or Strep-Tactin Elution Buffer (2.5 mL × 6) and further purified with a gradient elution method (FPLC A to FPLC B over 20 column volumes at 4 mL/min, 4 column volume wash with FPLC A prior to gradient) on an ÄKTA pure chromatography system (GE Healthcare Life Sciences) equipped with a HiTrap Q HP anion exchange chromatography column (5 mL). Fractions identified by SDS-PAGE to contain proteins of interest were pooled, concentrated using Amicon Ultra Centrifugal Filters (MilliporeSigma), frozen in liquid nitrogen and stored at -80°C.

All steps were carried out at 4°C or on ice.

#### *In Vitro* Reconstitution of Modules L, 1-5 and *t*AT-TEII

**Reaction Components**: 100 mM HEPES (Thermo Fisher Scientific), pH 7.5, 2.5 mM coenzyme A (MilliporeSigma), 10 mM adenosine 5′-triphosphate (MilliporeSigma), 5 mM NADPH (MilliporeSigma), 10 mM magnesium chloride (Thermo Fisher Scientific), 10 mM tris(2-carboxyethyl)phosphine (Thermo Fisher Scientific), 5 μM MatB, and 4-8 μM of all remaining proteins depending on scale (*t*AT-TEII, MBP-module L, module 1-2-DEBS(4), DEBS(5)-module 3-DEBS(2), DEBS(3)-module 4-$KS_5$, $DH_5$-$ACP_5$-$KR_5$).

Malonate ($^{12}C_3$, $^{13}C_{1,3}$, $^{13}C_2$, or $^{13}C_3$) (Millipore Sigma) was added to start the reaction (5 mM final concentration), which was incubated overnight (24 hrs) at room temperature in the dark. Reactions were quenched by the addition of LC-MS grade methanol (Thermo Fisher Scientific) (1:1 reaction volume/methanol). The mixture was vortexed followed by centrifugation at 16,000 × *g* for 10 min at room temperature. The supernatant was transferred to a 0.2 μm PTFE spin filter (Thermo Scientific™) and centrifuged at 16,000 × *g* for 1 min before transferring to a LC-MS vial. For LC-MS analysis, compounds were separated with a gradient elution method (95:5 % HPLC A:HPLC B to 5:95 % HPLC A:HPLC B over 4 min at 0.6 mL/min) on an Agilent Infinity 1290 II HPLC/6545 Q-TOF MS system equipped with an Agilent ZORBAX RRHD Extend-C18 column (1.8 μm, 2.1 x 50 mm) operating in negative or positive ion mode.

In this and all assays utilizing both malonate and coenzyme A, the *Streptomyces coelicolor* malonyl-CoA synthetase MatB generated malonyl-CoA *in situ*.

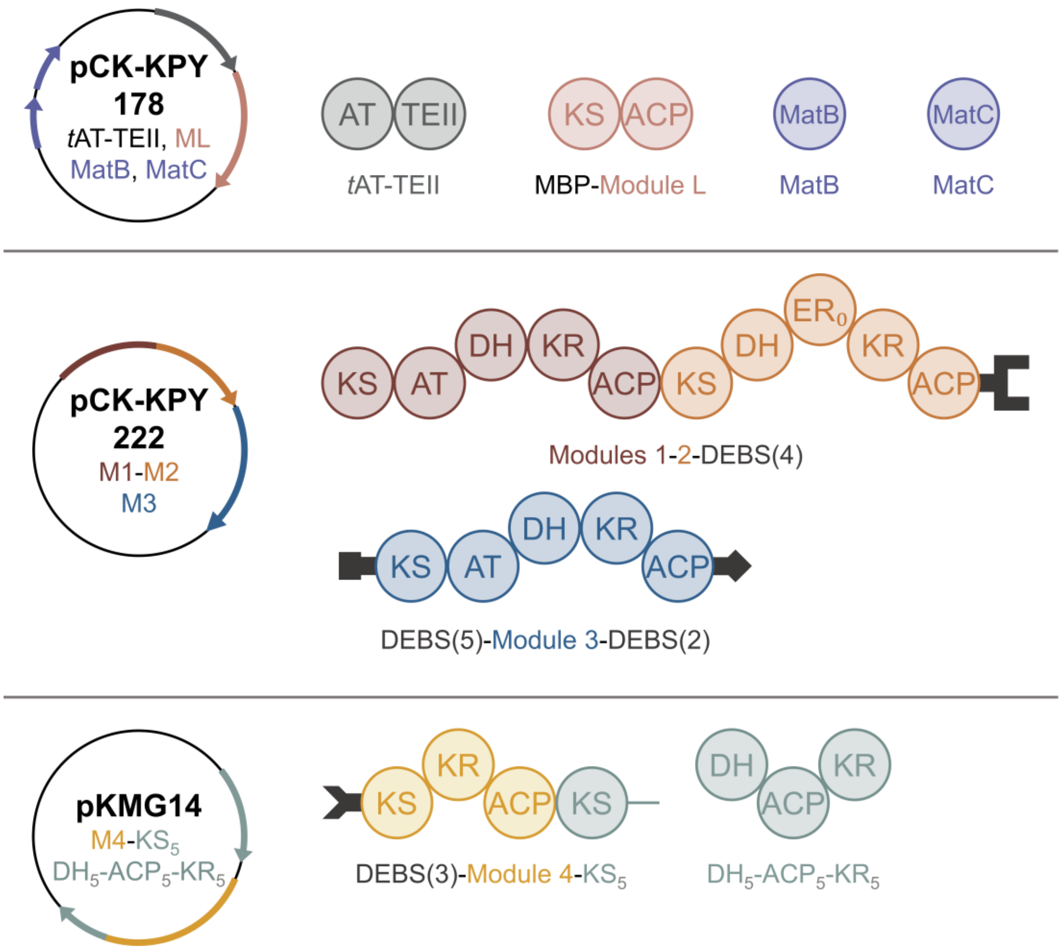

**Figure S1**. Plasmids used for biosynthesis of **3** and **4** in *E. coli* BAP1.

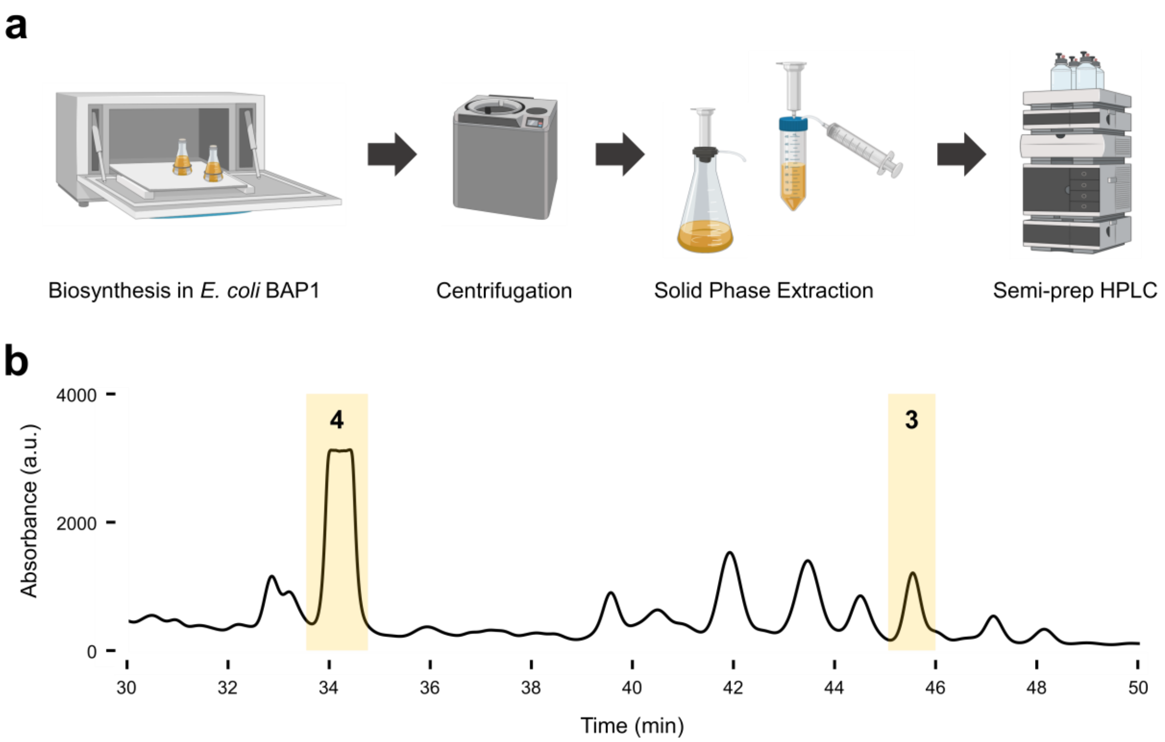

**Figure S2**. Isolation of **3** and **4** from *E. coli* BAP1 [pCK-KPY178/pCK-KPY222/pKMG14] supernatant. (a) Two step extraction: solid phase extraction column (SPE) followed by semi-preparative HPLC purification. (b) Extracted UV chromatogram (absorbance at 280 nm ± 10 nm) from HPLC purification of **3** and **4**. Data acquired on an Agilent 1260 Infinity LC system, and are representative of at least three independent rounds of compound biosynthesis and isolation.

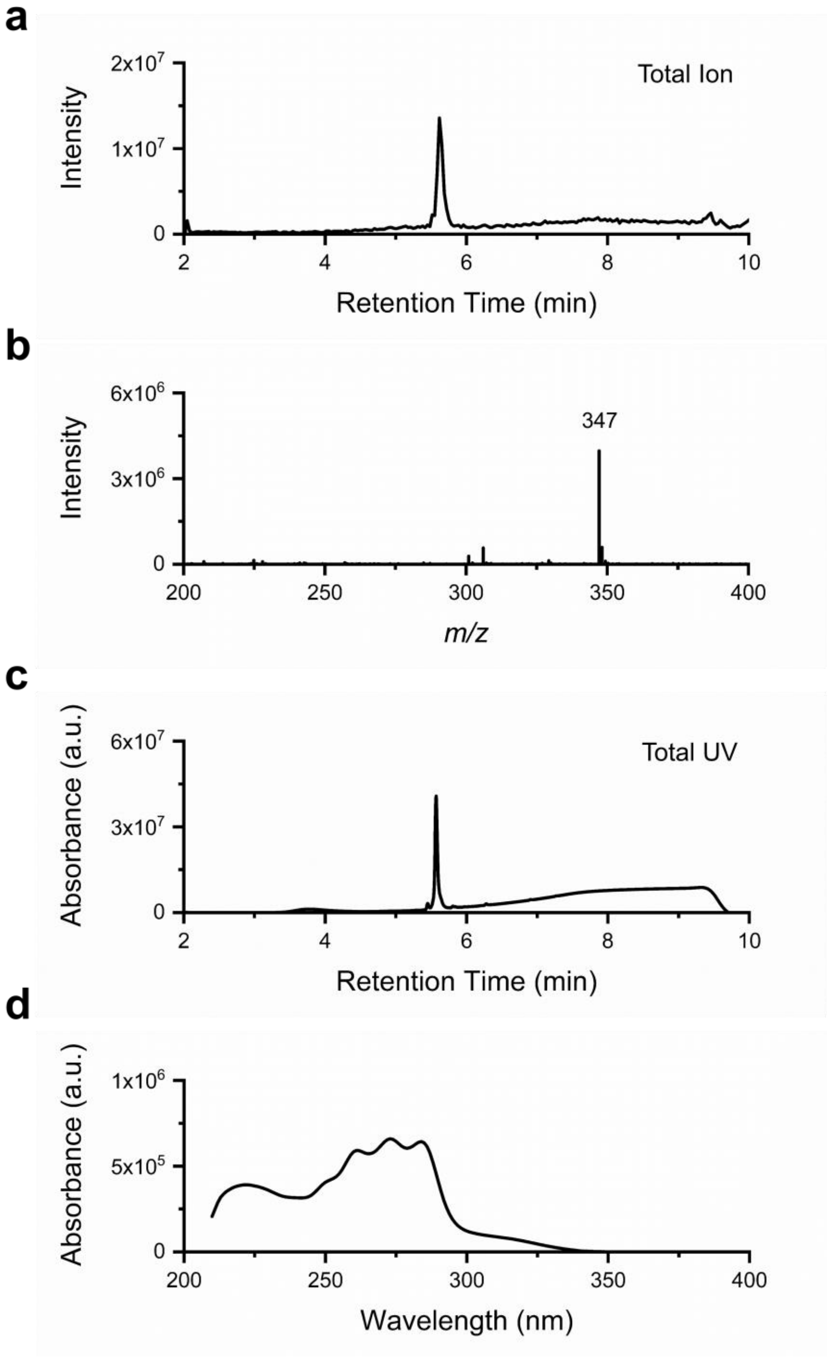

**Figure S3**. (a) Total ion chromatogram (ESI-), (b) MS spectrum (ESI-), (c) total UV chromatogram and (d) UV spectrum of purified **3**. Data were acquired on a Waters SQ Detector 2 LC-MS system and are representative of at least three independent rounds of compound biosynthesis and isolation.

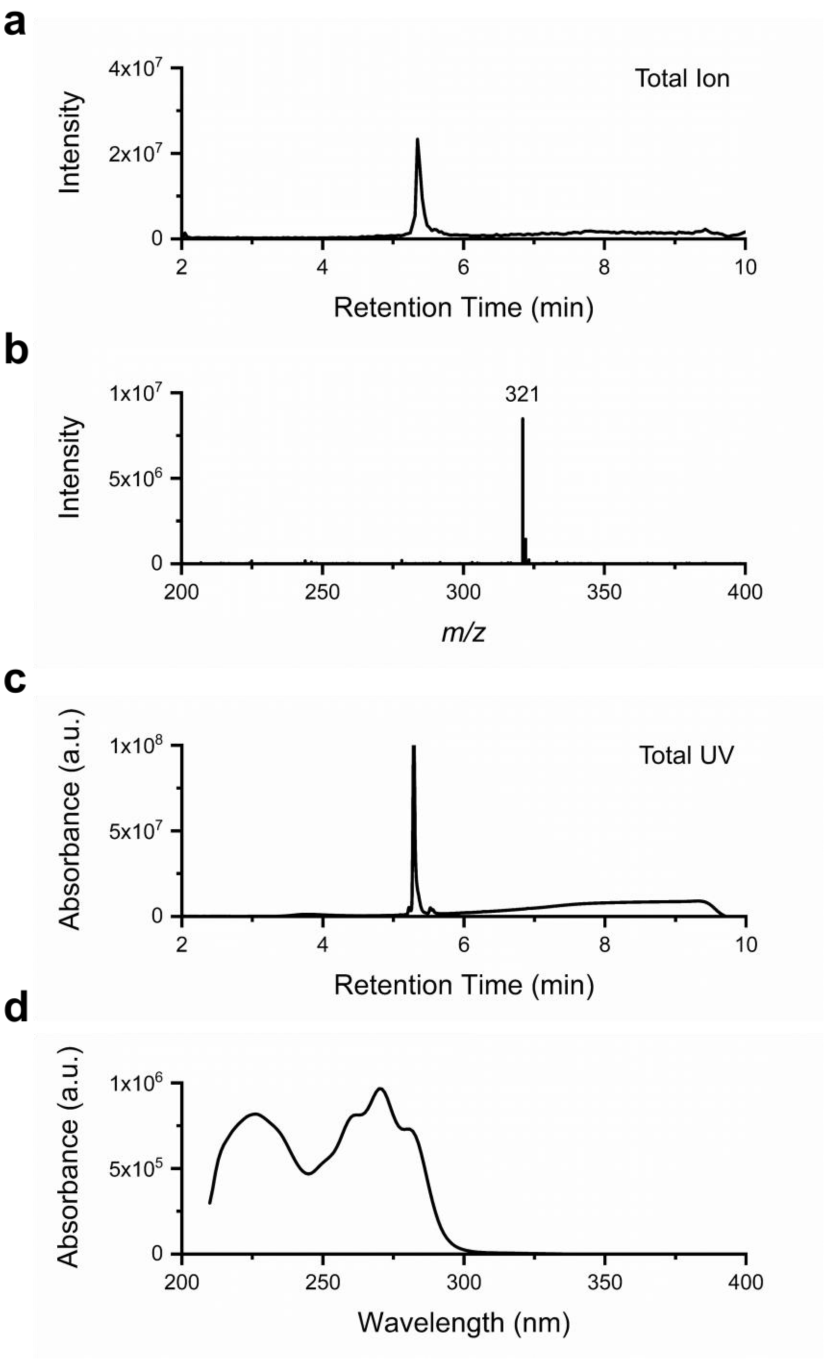

**Figure S4**. (a) Total ion chromatogram (ESI-), (b) MS spectrum (ESI-), (c) total UV chromatogram and (d) UV spectrum of purified **4**. Data were acquired on a Waters SQ Detector 2 LC-MS system and are representative of at least three independent rounds of compound biosynthesis and isolation.

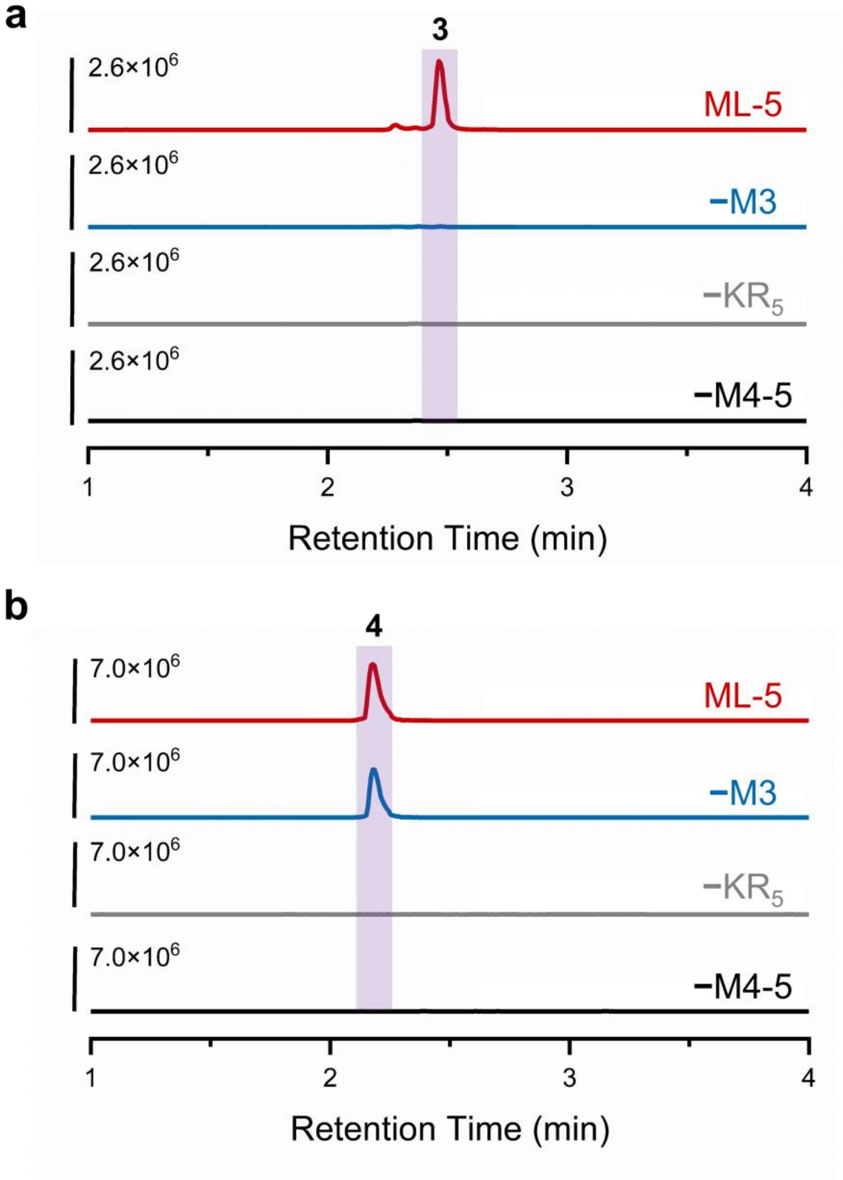

**Figure S5**. (a) ESI(-) EIC of **3** (*m/z* 347.1864 ± 0.1) and (b) **4** (*m/z* 321.1707 ± 0.1) in various engineered *E. coli* BAP1 strains. Removal of module 3 [pCK-KPY178/pCK-KPY102/pKMG14 strain] (blue curve) abolished production of **3** but not **4**, similar to the biosynthesis of **1** and **2**. Additionally, when $KR_5$ [pCK-KPY178/pCK-KPY222/pKMG15] (grey curve) or Module 4-$KS_5$ and $DH_5$-$ACP_5$-$KR_5$ [pCK-KPY178/pCK-KPY222] (black curve) are removed, **3** and **4** are no longer present. Data were acquired on a Waters SQ Detector 2 LC-MS system and are representative of at least three independent rounds of compound biosynthesis and isolation.

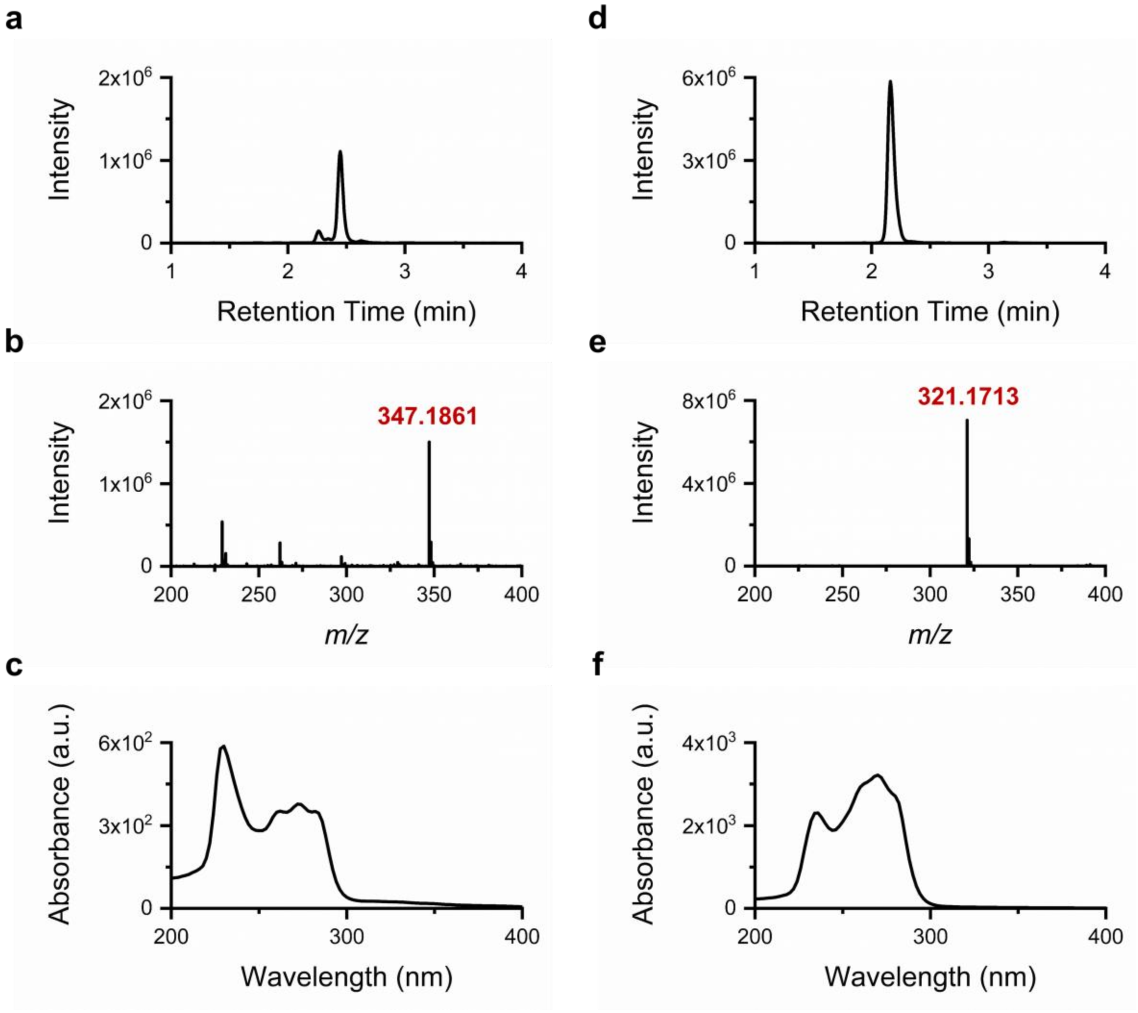

**Figure S6**. HPLC-HRMS (a) ESI(−) EIC 347.1864 (± 0.1), (b) mass spectrum, and (c) UV spectrum of **3** extracted from *E. coli* BAP1 [pCK-KPY178/pCK-KPY222/pKMG14] supernatant. **3** (*m/z*: $[M-H]^-$ calcd. for $C_{20}H_{27}O_5^-$ 347.1864, obsd. 347.1845, 5.5 ppm). (d) ESI(−) EIC 321.1707 (± 0.1), (e) mass spectrum, and (f) UV spectrum of **4** extracted from *E. coli* BAP1 [pCK-KPY178/pCK-KPY222/pKMG14] supernatant. **4** (*m/z*: $[M-H]^-$ calcd. for $C_{18}H_{25}O_5^-$ 321.1707, obsd. 321.1704, 0.9 ppm). Data were acquired on an Agilent Q-TOF 6545 LC-MS system and are representative of at least three independent rounds of compound biosynthesis. HRMS calculated values based on ChemDraw software predictions.

**Table S1**. NMR assignments* for **3** and **4** in $CD_3OD$ (900 MHz and 600 MHz, respectively)

| | **3** | | **4** | |
|---|---|---|---|---|
| Carbon | δC | δH | δC | δH |
| 1 | 175.93 | - | 176.40 | - |
| 2 | 43.40 | 2.48 | 43.19 | 2.46 |
| 3 | 66.02 | 4.26 | 65.94 | 4.26 |
| 4 | 44.93 | 1.65 | 44.68 | 1.65 |
| 5 | 69.40 | 4.40 | 69.42 | 4.40 |
| 6 | 137.74 | 5.74 | 137.66 | 5.73 |
| 7 | 126.06 | 6.73 | 125.87 | 6.73 |
| 8 | 128.32 | 5.91 | 128.20 | 5.90 |
| 9 | 130.57 | 5.96 | 130.36 | 5.95 |
| 10 | 129.12 | 6.61 | 128.86 | 6.60 |
| 11 | 132.34 | 5.72 | 132.18 | 5.70 |
| 12 | 41.80 | 2.37 | 41.69 | 2.36 |
| 13 | 72.94 | 4.16 | 72.74 | 4.12 |
| 14 | 135.40 | 5.66 | 133.44 | 5.53 |
| 15 | 131.86 | 6.21 | 131.68 | 6.16 |
| 16 | 130.57 | 6.09 | 131.85 | 6.04 |
| 17 | 134.27 | 6.19 | 130.36 | 5.71 |
| 18 | 132.66 | 6.08 | 17.95 | 1.73 |
| 19 | 131.05 | 5.73 | | |
| 20 | 18.37 | 1.75 | | |

*Assignments based on HSQC shifts referenced to $CD_3OD$ (49, 3.31 ppm), except C1 based on $^{13}C$ 1D data (900 MHz) referenced to $CD_3OD$ (49 ppm)

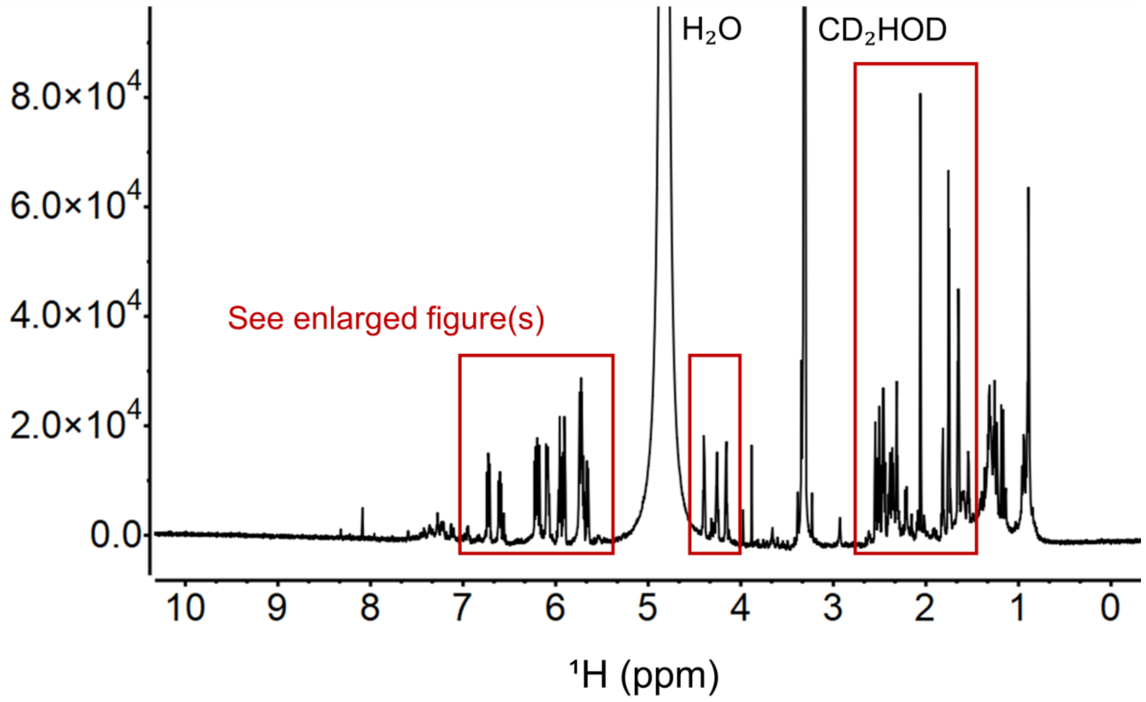

**Figure S7**. $^{1}H$ NMR spectrum (x-axis: δH, 10.0-0.0 ppm) of **3** in $CD_3OD$. Spectrum acquired on a 900 MHz Bruker AVANCE II spectrometer.

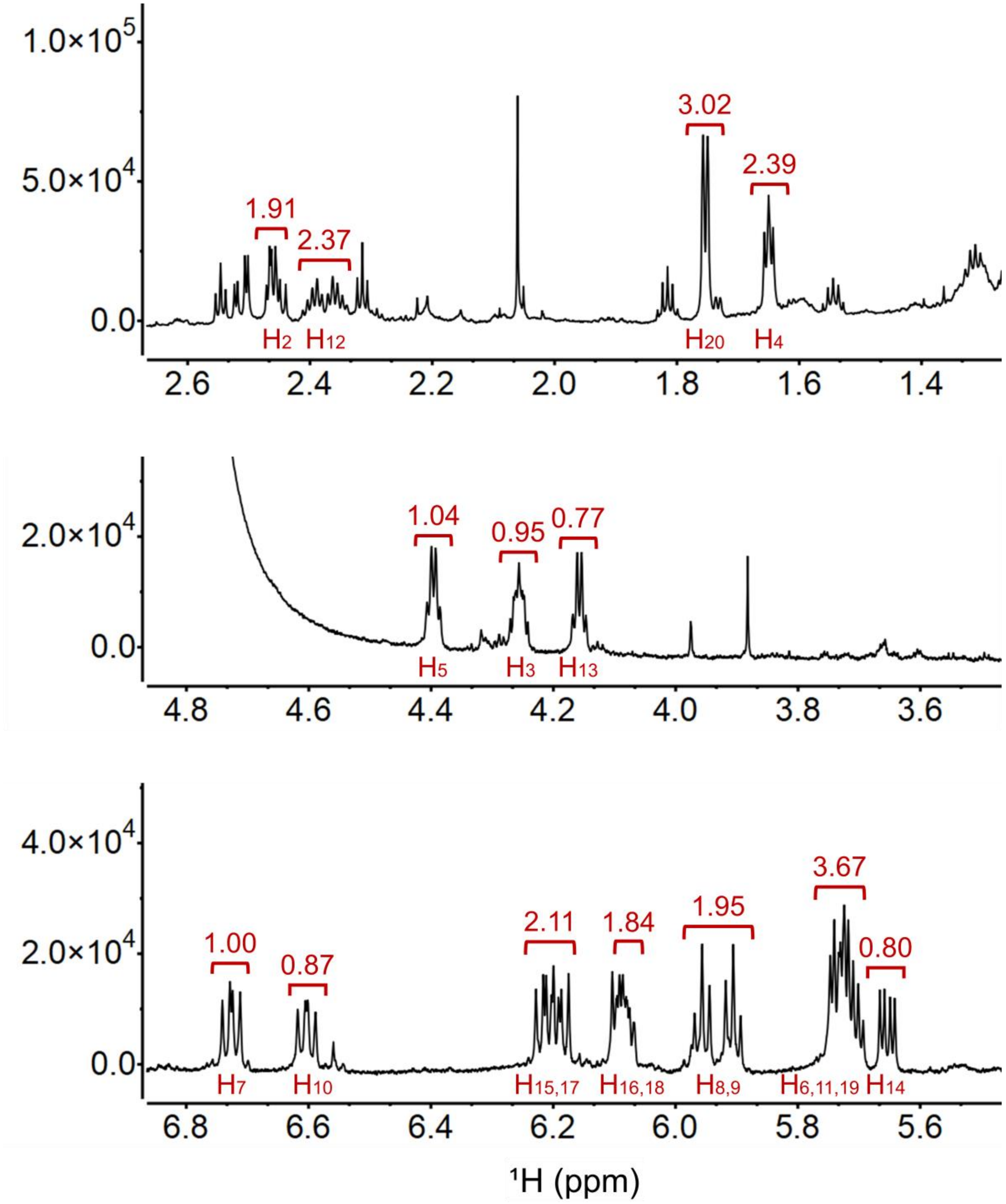

**Figure S8**. Enlarged regions of $^{1}H$ NMR spectrum of **3** in $CD_3OD$. Integration values are represented above peaks (red); proton assignments listed below peaks. Spectrum acquired on a 900 MHz Bruker AVANCE II spectrometer.

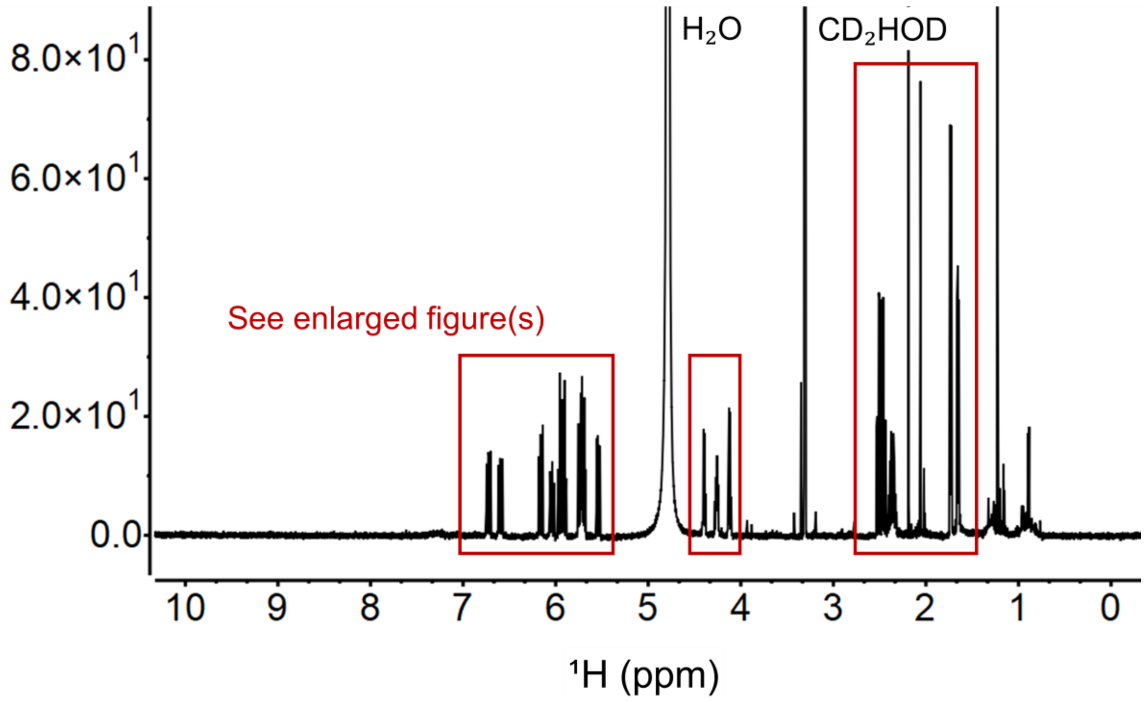

**Figure S9**. $^1$H NMR spectrum (x-axis: δH, 10.0-0.0 ppm) of **4** in $CD_3OD$. Spectrum acquired on a 600 MHz Varian Inova spectrometer.

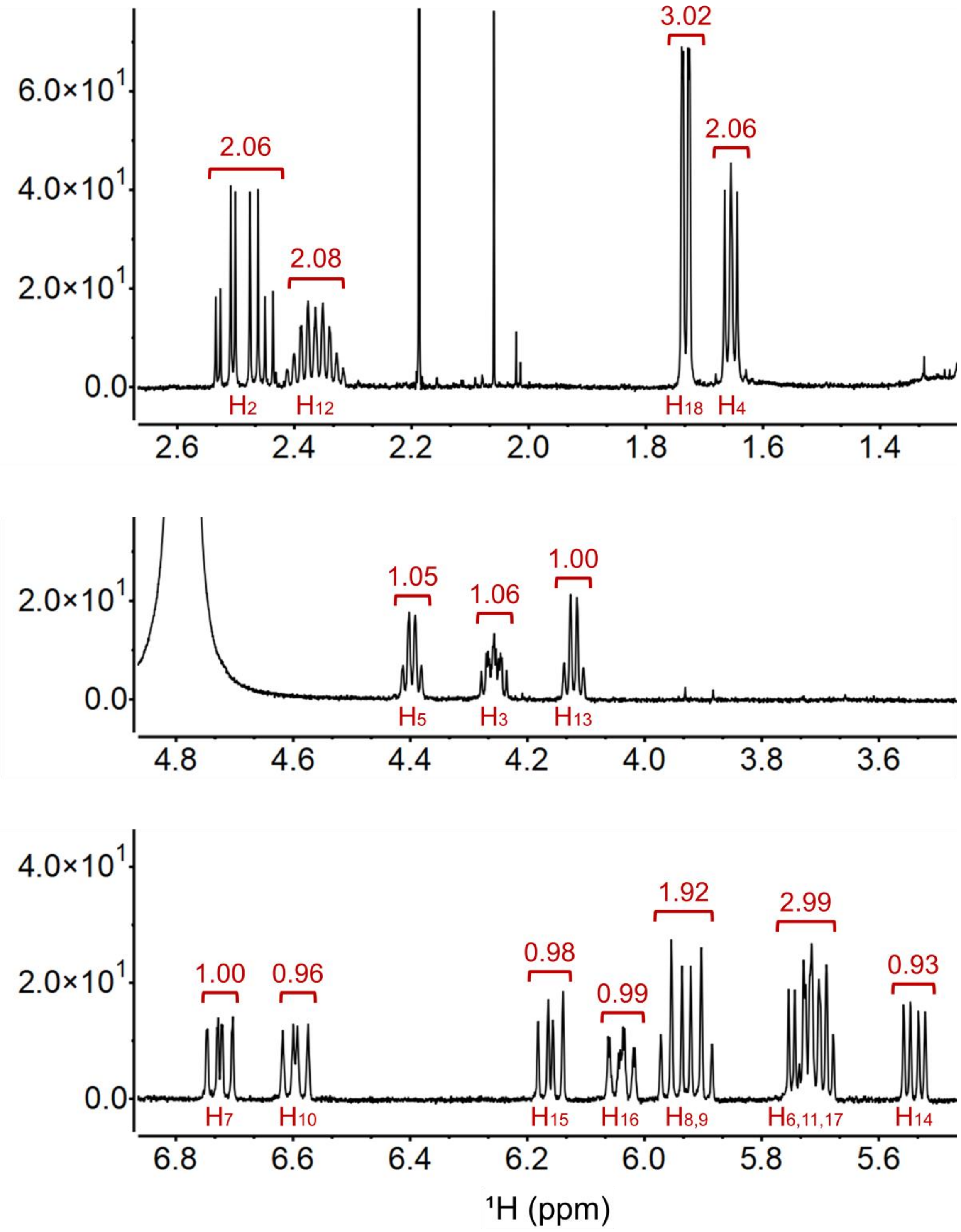

**Figure S10**. Enlarged regions of $^{1}H$ NMR spectrum of **4** in $CD_3OD$. Integration values represented above peaks (red); proton assignments listed below peaks. Spectrum acquired on a 600 MHz Varian Inova spectrometer.

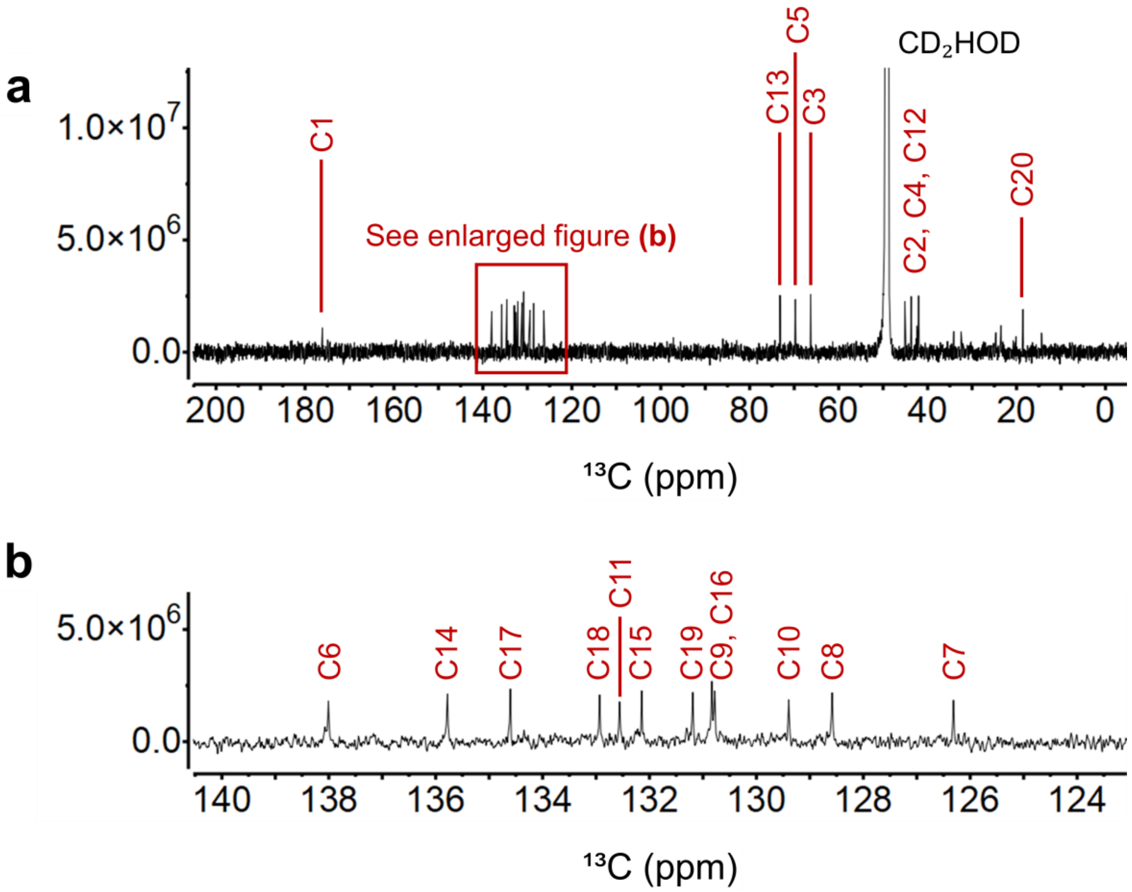

**Figure S11**. (a) $^{13}C$ NMR spectrum (x-axis: δC, 200-0 ppm) of **3** in $CD_3OD$. (b) Enlargement of downfield region for clarity. Spectrum acquired on a 900 MHz Bruker AVANCE II spectrometer.

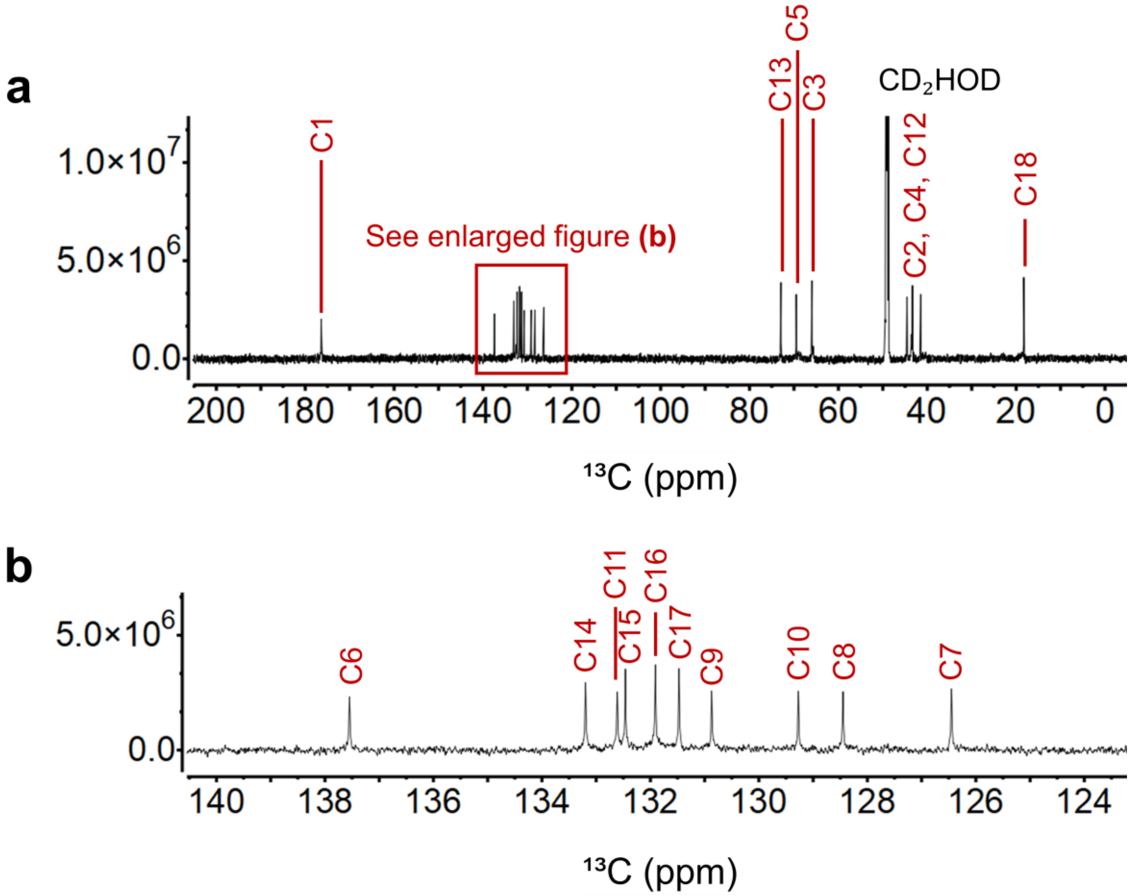

**Figure S12**. (a) $^{13}$C NMR spectrum (x-axis: δC, 200-0 ppm) of **4** in $CD_3OD$. (b) Enlargement of downfield region for clarity. Spectrum acquired on a 900 MHz Bruker AVANCE II spectrometer.

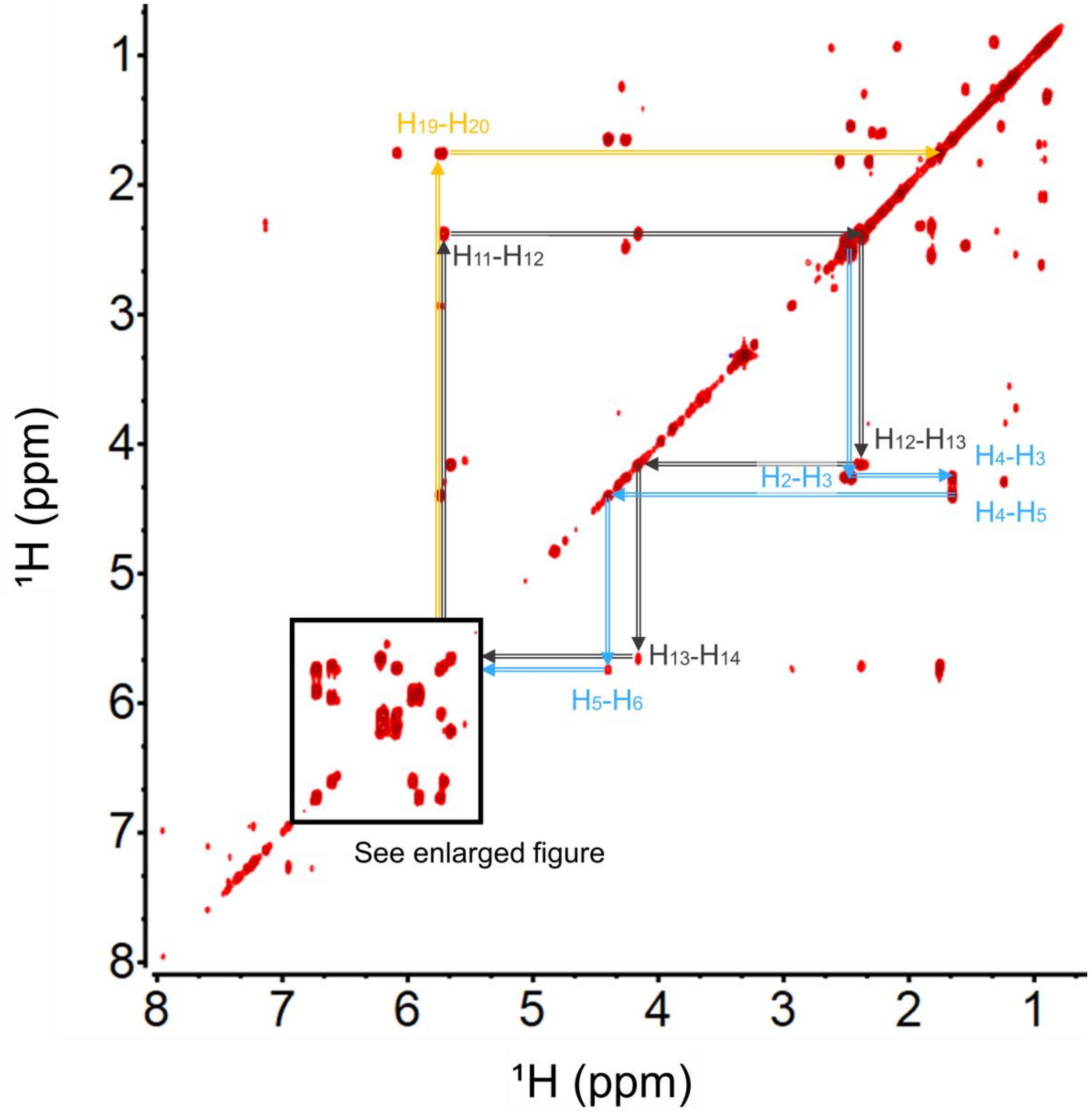

**Figure S13**. $^1$H-$^1$H COSY NMR spectrum (positive signals in red; x-axis: δH, 8.0-1.0 ppm; y-axis: δH, 8.0-1.0 ppm) of **3** in $CD_3OD$. Spectrum acquired on a 900 MHz Bruker AVANCE II spectrometer.

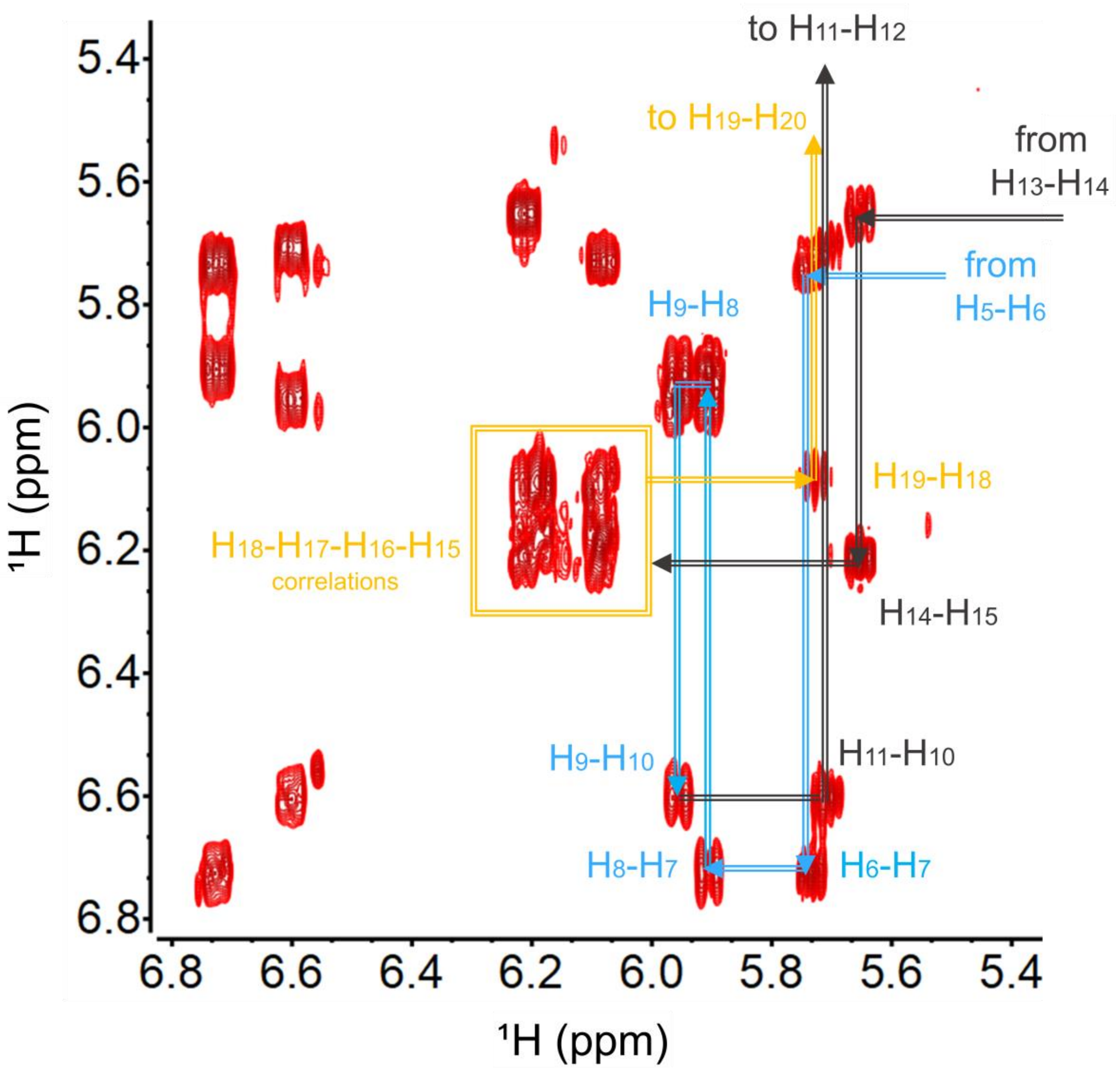

**Figure S14**. Enlarged region of $^1H$-$^1H$ COSY NMR spectrum (positive signals in red; x-axis: δH, 6.8-5.4 ppm; y-axis: δH, 6.8-5.4 ppm) of **3** in $CD_3OD$ (for assignment clarity). Spectrum acquired on a 900 MHz Bruker AVANCE II spectrometer.

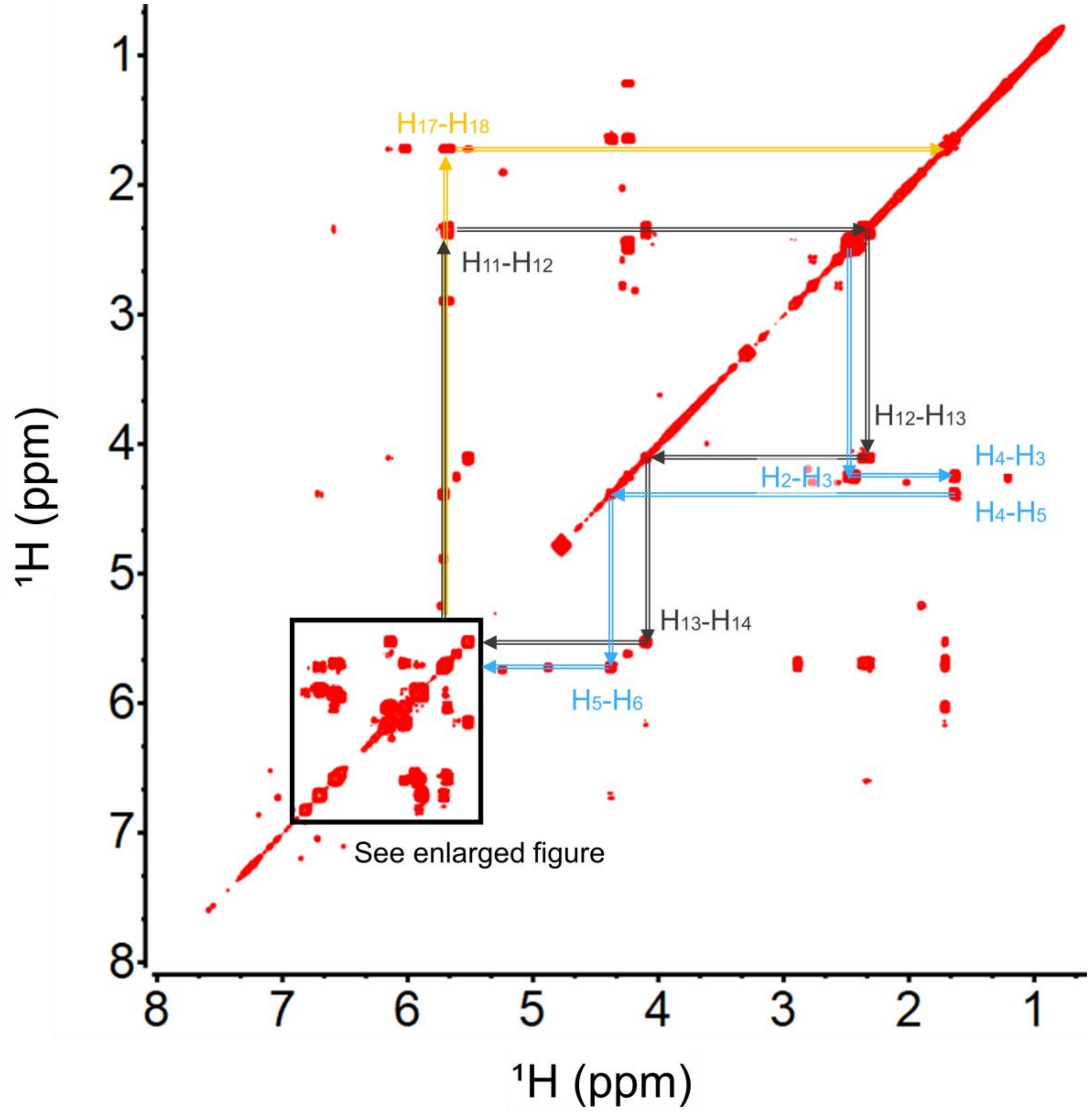

**Figure S15**. $^1$H-$^1$H COSY NMR spectrum (positive signals in red; x-axis: δH, 8.0-1.0 ppm; y-axis: δH, 8.0-1.0 ppm) of **4** in $CD_3OD$. Spectrum acquired on a 600 MHz Varian Inova spectrometer.

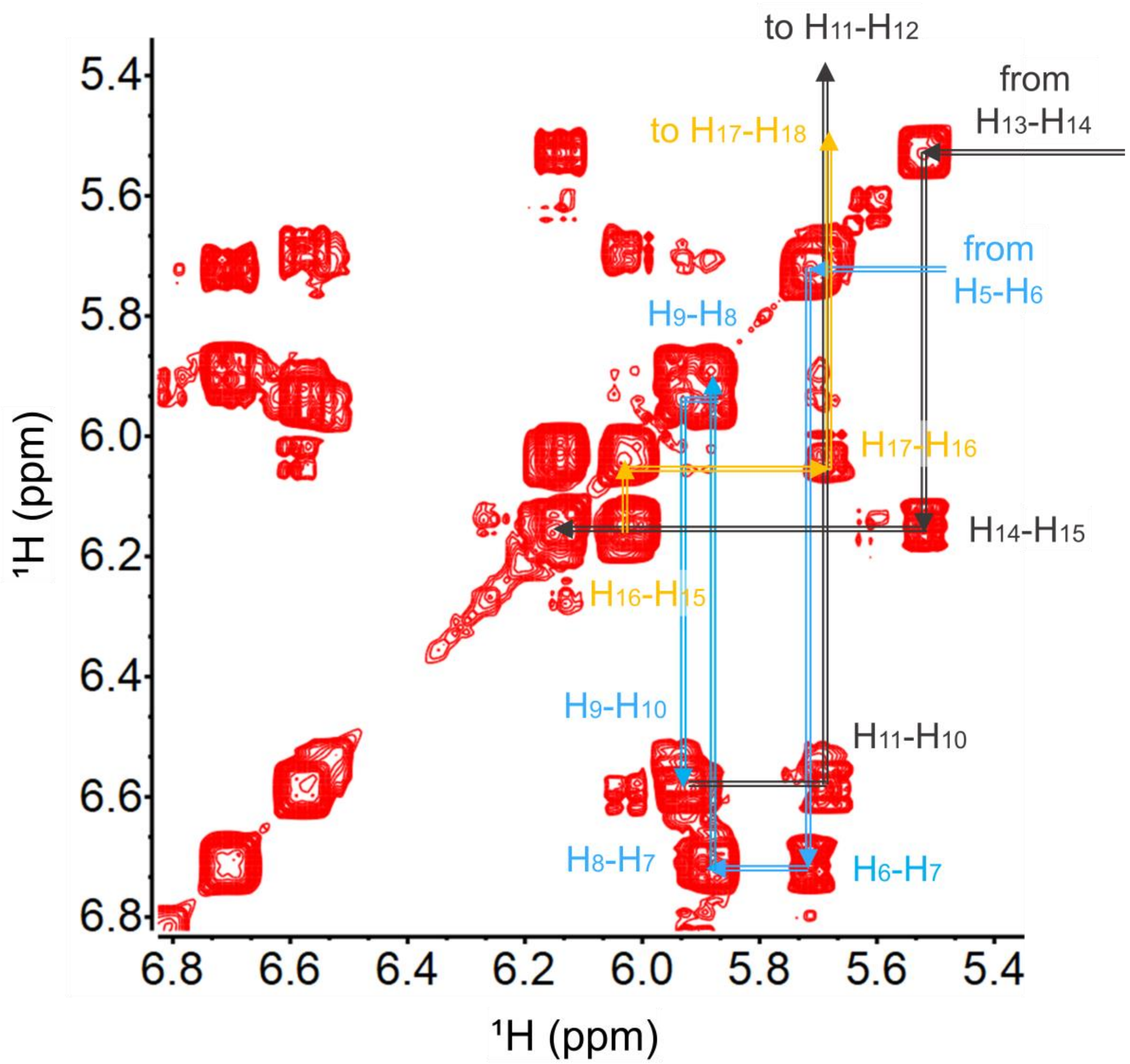

**Figure S16**. Enlarged region of ${}^1H$-${}^1H$ COSY NMR spectrum (positive signals in red; x-axis: δH, 6.8-5.4 ppm; y-axis: δH, 6.8-5.4 ppm) of **4** in $CD_3OD$ (for assignment clarity). Spectrum acquired on a 600 MHz Varian Inova spectrometer.

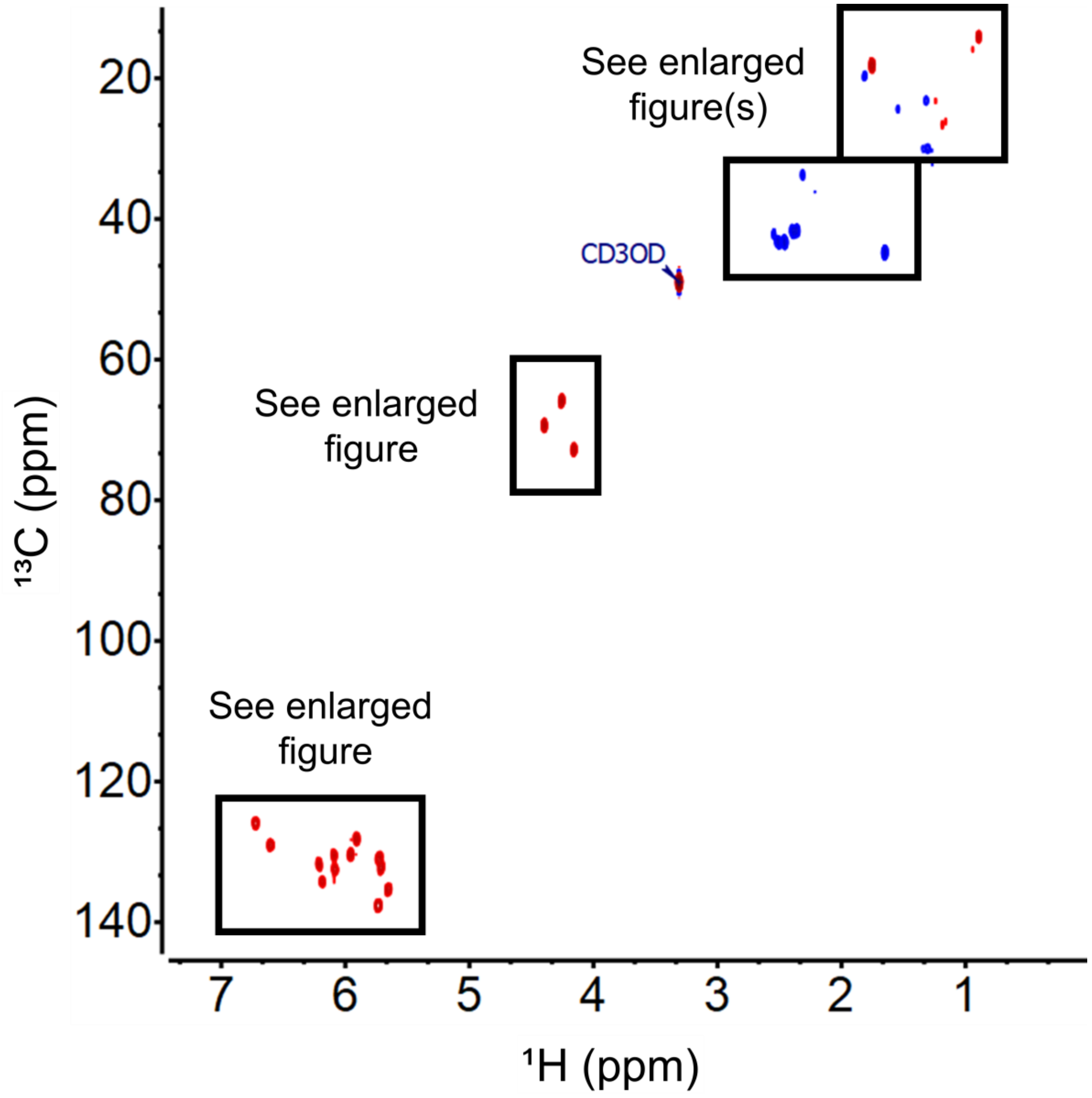

**Figure S17**. $^{1}H$-$^{13}C$ HSQC spectrum (positive signals in red; negative signals in blue; x-axis: δH, 7.4-0.0 ppm; y-axis: δC, 140.0-12.0 ppm) of **3** in $CD_3OD$. Spectrum acquired on a 900 MHz Bruker AVANCE II spectrometer.

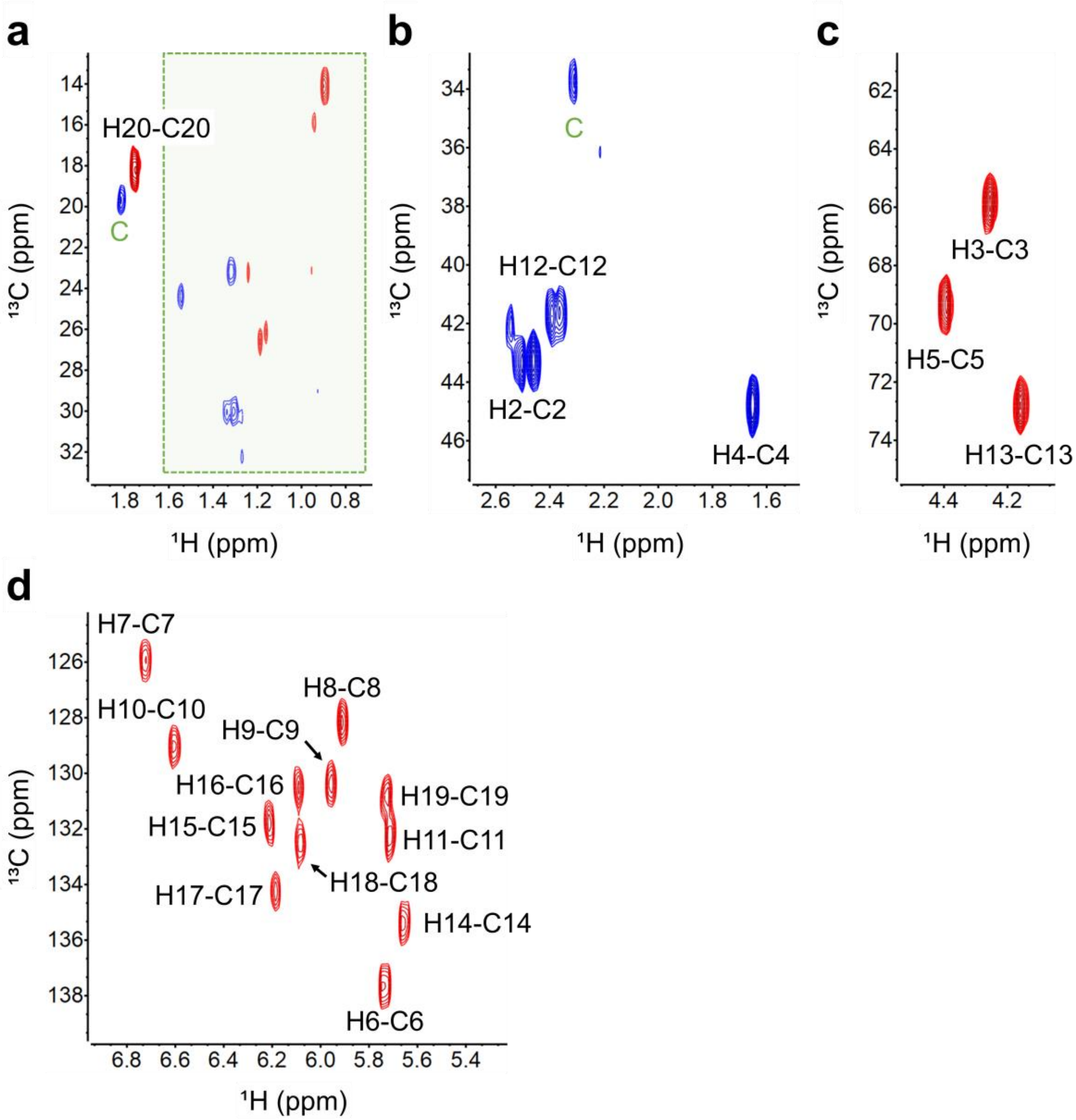

**Figure S18**. Enlarged regions of $^{1}H$-$^{13}C$ HSQC spectrum (positive signals in red (CH- and $CH_3$- groups); negative signals in blue ($CH_2$- groups)) of **3** in $CD_3OD$. (a) x-axis: δH, 1.9-0.7 ppm; y-axis: δC, 34.0-12.0 ppm; (b) x-axis: δH, 2.7-1.5 ppm; y-axis: δC, 48.0-32.0 ppm; (c) x-axis: δH, 4.6-4.0 ppm; y-axis: δC, 76.0-60.0 ppm; (d) x-axis: δH, 7.0-5.2 ppm; y-axis: δC, 140.0-124.0 ppm. Key: C, contaminant; additionally, green boxes represent groups of contaminate peaks. Spectrum acquired on a 900 MHz Bruker AVANCE II spectrometer.

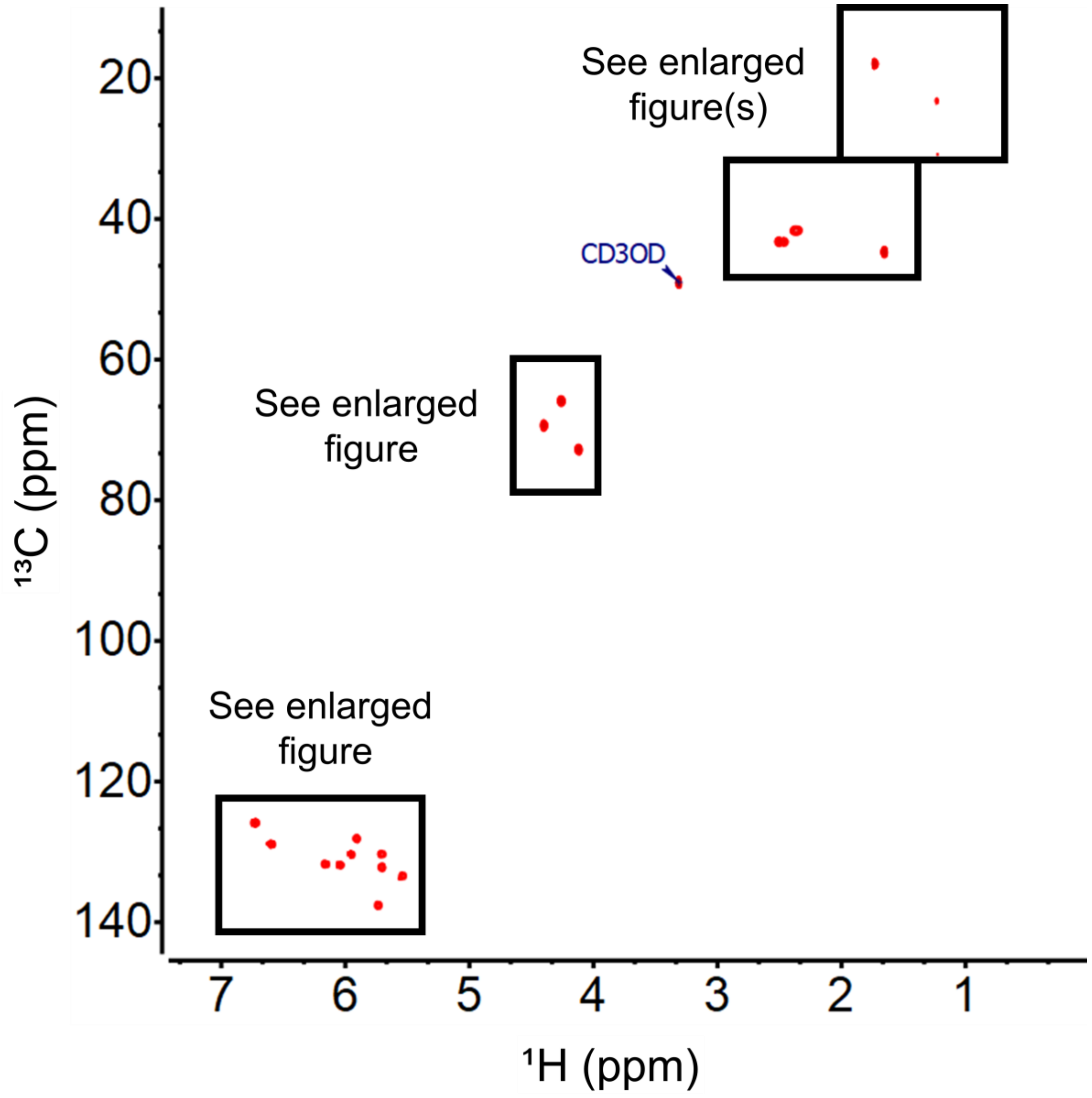

**Figure S19**. $^{1}$H-$^{13}$C HSQC-SE* spectrum (positive signals in red; x-axis: δH, 7.4-0.0 ppm; y-axis: δC, 140.0-12.0 ppm) of **4** in $CD_3OD$. Spectrum acquired on a 600 MHz Varian Inova spectrometer. *Sensitivity Enhanced HSQC (HSQC-SE) does not include multiplicity editing (CH, $CH_2$, $CH_3$, are all positive)

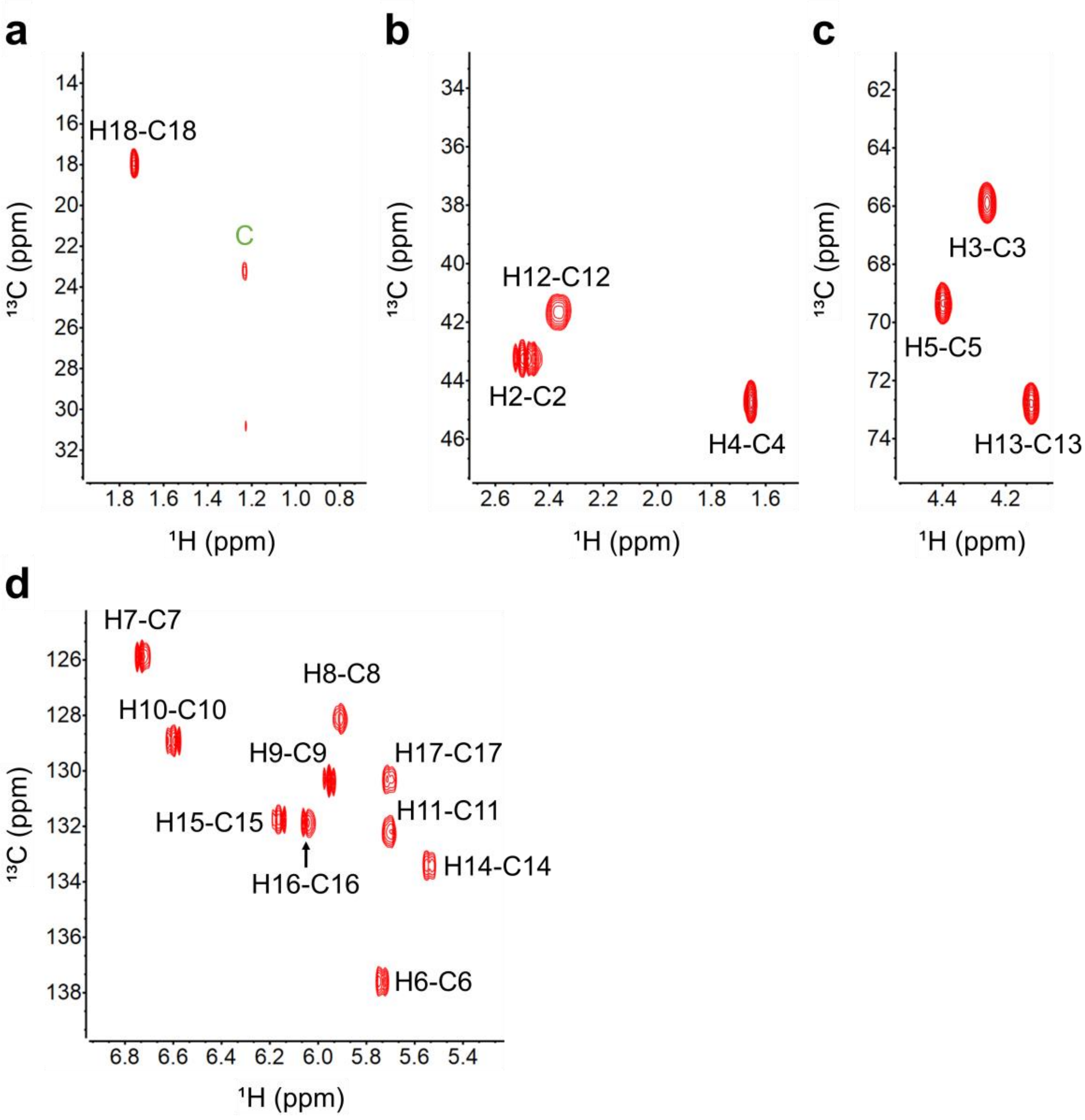

**Figure S20**. Enlarged regions of $^{1}H$-$^{13}C$ HSQC-SE* spectrum (positive signals in red) of **4** in $CD_3OD$. (a) x-axis: δH, 1.9-0.7 ppm; y-axis: δC, 34.0-12.0 ppm; (b) x-axis: δH, 2.7-1.5 ppm; y-axis: δC, 48.0-32.0 ppm; (c) x-axis: δH, 4.6-4.0 ppm; y-axis: δC, 76.0-60.0 ppm; (d) x-axis: δH, 7.0-5.2 ppm; y-axis: δC, 140.0-124.0 ppm. Key: C, contaminant. Spectrum acquired on a 600 MHz Varian Inova spectrometer. *Sensitivity Enhanced HSQC (HSQC-SE) does not include multiplicity editing (CH, $CH_2$, $CH_3$, are all positive)

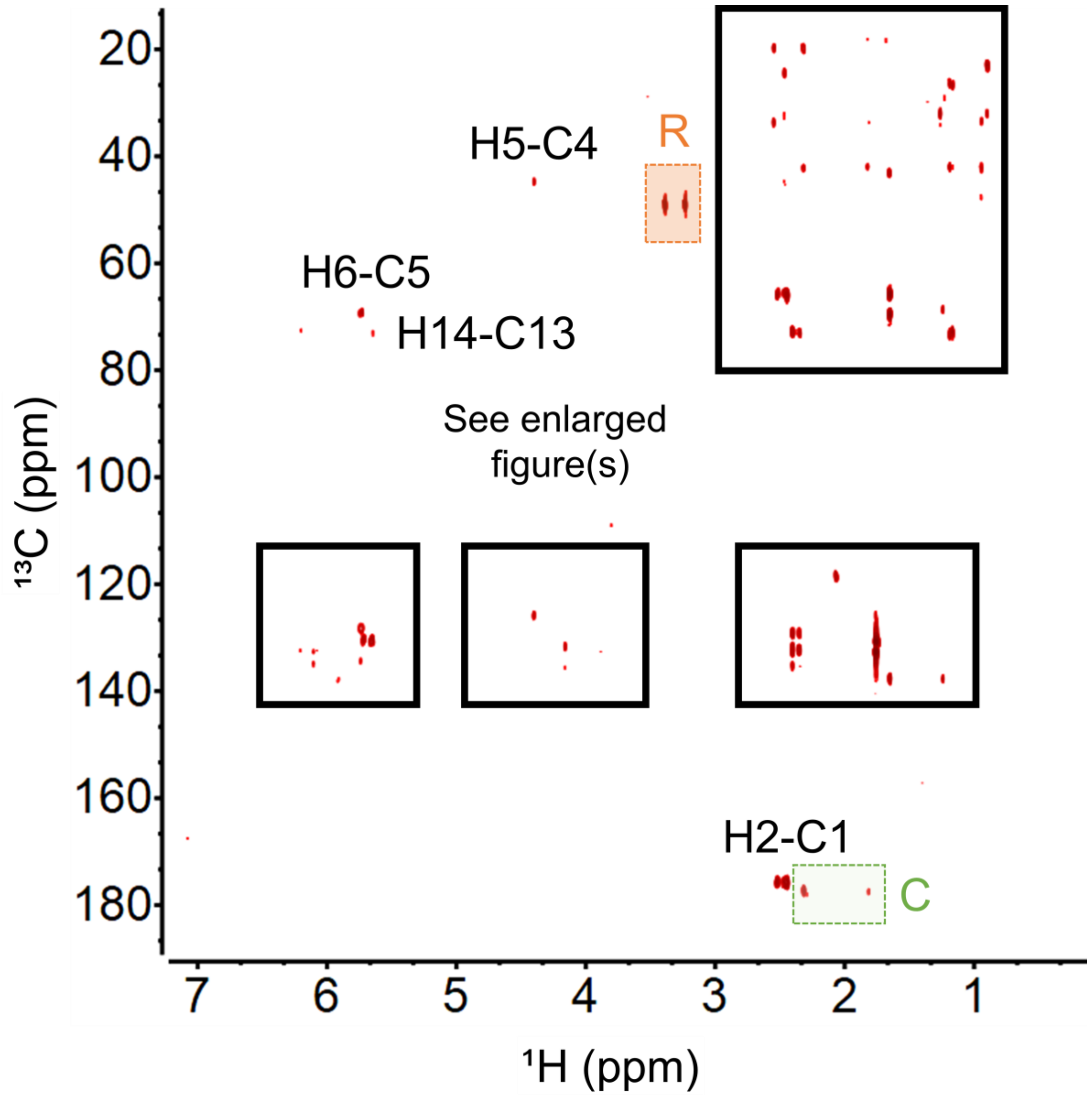

**Figure S21.** $^{1}$H-$^{13}$C HMBC spectrum (positive signals in red; x-axis: δH, 7.2-0.0 ppm; y-axis: δC, 187.0-14.0 ppm) of **3** in $CD_3OD$. Key: C, contaminant; R, residual solvent signal; additionally, green boxes represent groups of contaminate peaks. Spectrum acquired on a 900 MHz Bruker AVANCE II spectrometer.

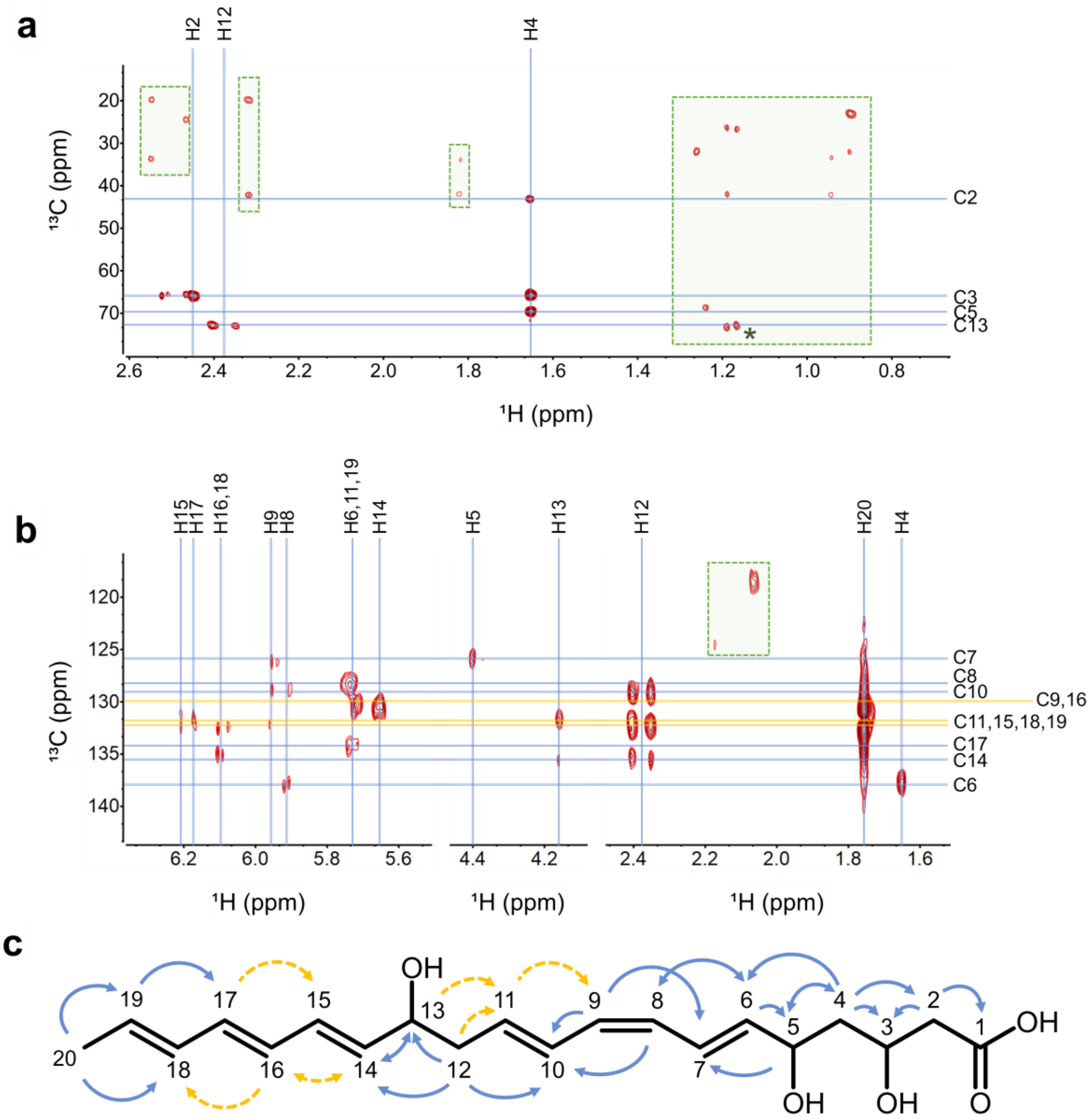

**Figure S22.** (a,b) Enlarged regions of $^{1}H$-$^{13}C$ HMBC spectrum (positive signals in red) of **3** in $CD_3OD$. (c) NMR correlations seen in (a),(b), and Figure S21; blue arrows represent strong confidence in assignment whereas yellow-dashed arrows represent areas of degenerate (overlapping) signals where best judgement was applied. Key: C, contaminant; additionally, green boxes represent groups of contaminate peaks. Spectrum acquired on a 900 MHz Bruker AVANCE II spectrometer.

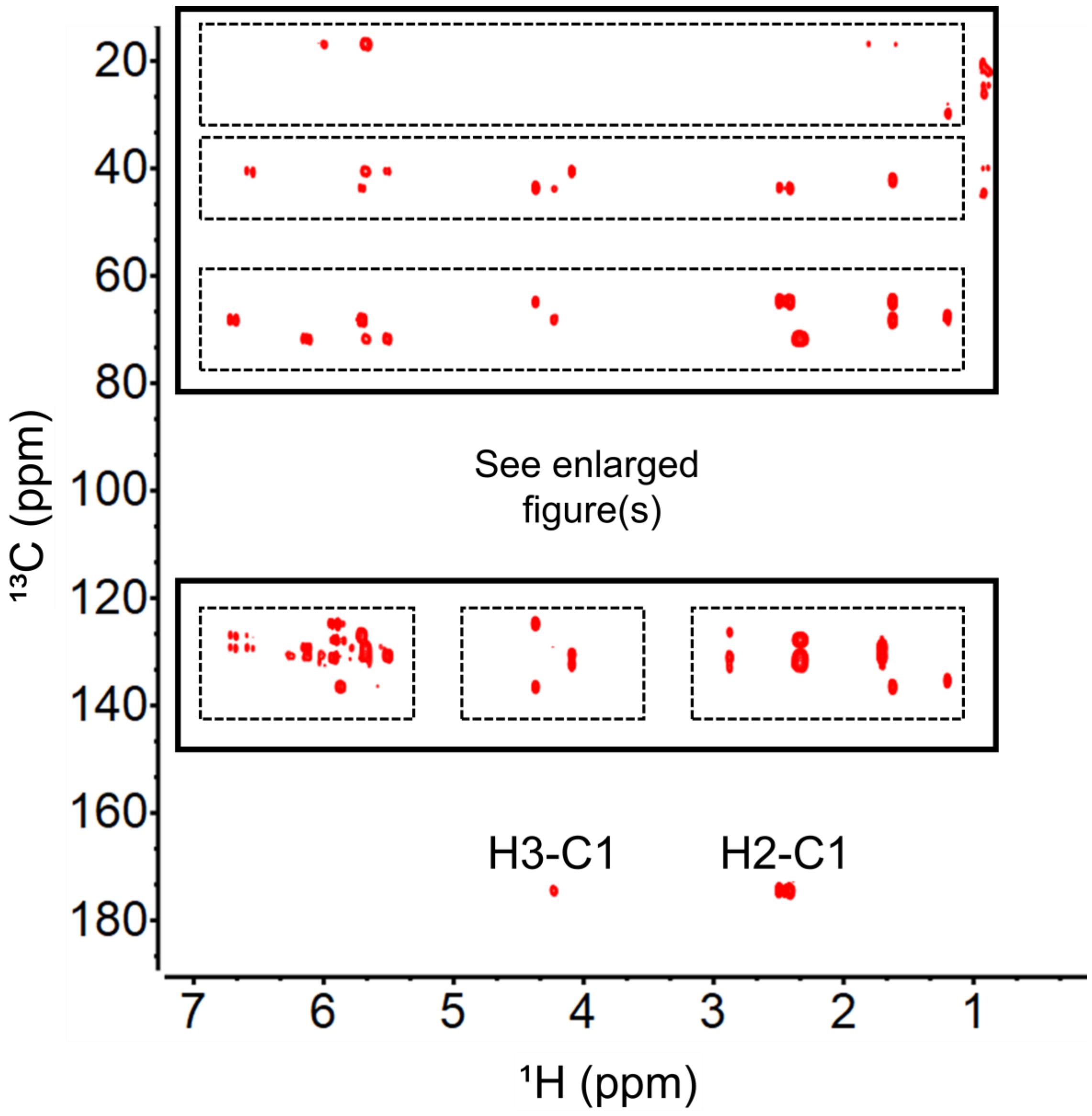

**Figure S23.** $^{1}H$-$^{13}C$ HMBC spectrum (positive signals in red; x-axis: δH, 7.2-0.0 ppm; y-axis: δC, 187.0-14.0 ppm) of **4** in $CD_3OD$. Spectrum acquired on a 600 MHz Varian Inova spectrometer.

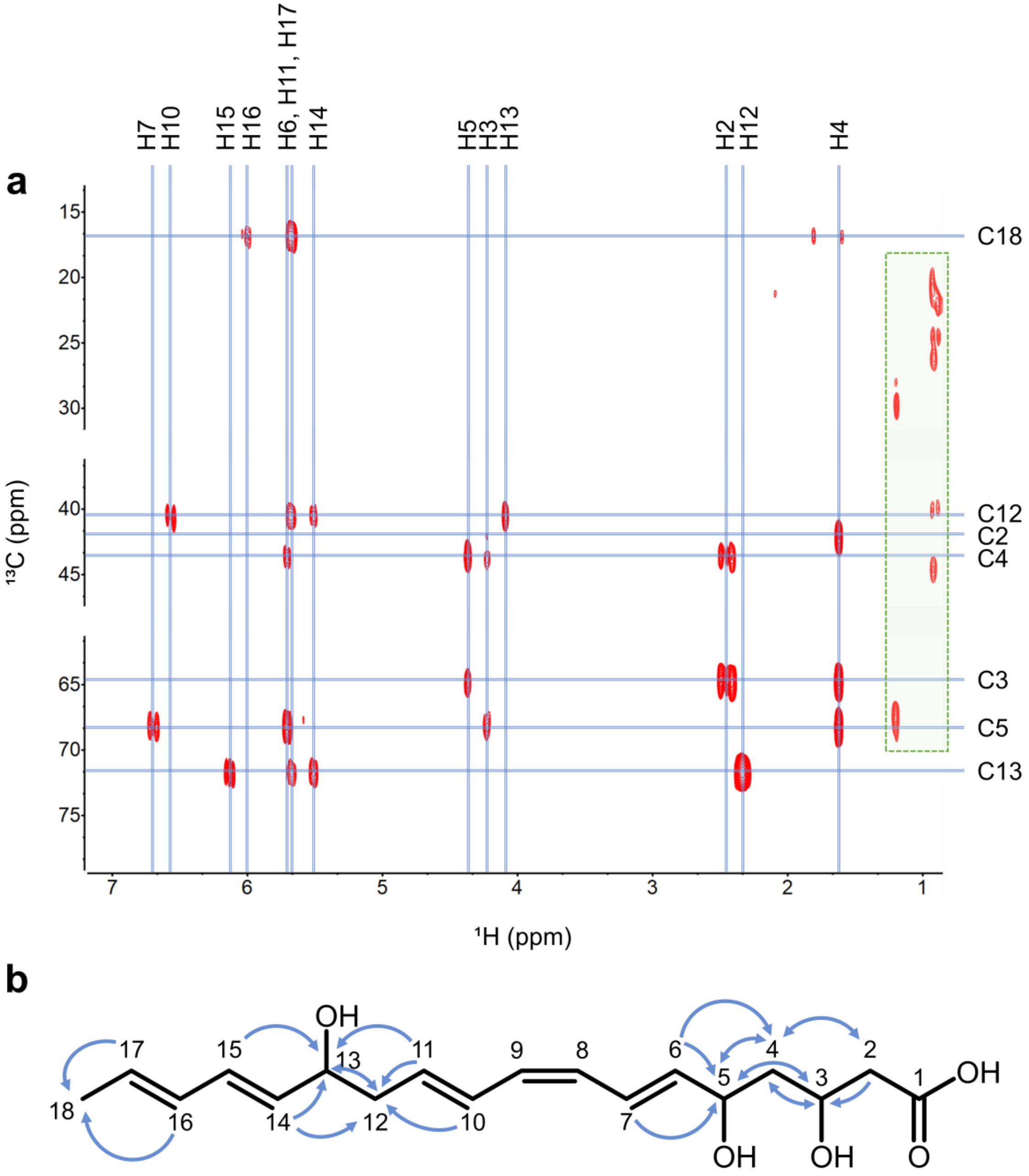

**Figure S24.** (a) Enlarged regions of $^1$H-$^{13}$C HMBC spectrum (positive signals in red) of **4** in $CD_3OD$. (b) NMR correlations seen in (a); blue arrows represent strong confidence in assignment. Key: C, contaminant; additionally, green boxes represent groups of contaminate peaks. Spectrum acquired on a 600 MHz Varian Inova spectrometer.

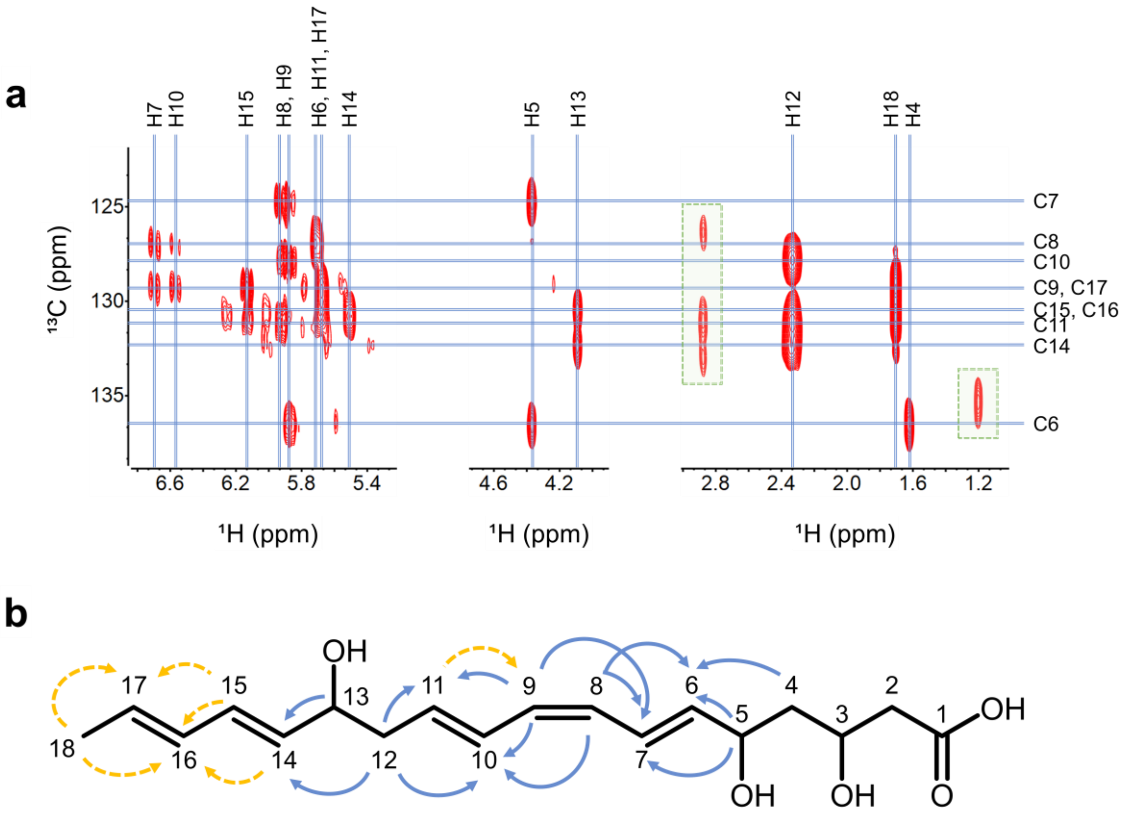

**Figure S25.** (a) Enlarged regions of $^{1}H$-$^{13}C$ HMBC spectrum (positive signals in red) of **4** in $CD_3OD$ (b) NMR correlations seen in (a); blue arrows represent strong confidence in assignment whereas yellow-dashed arrows represent areas of degenerate (overlapping) signals where best judgement was applied. .Key: C, contaminant; additionally, green boxes represent groups of contaminate peaks. Spectrum acquired on a 600 MHz Varian Inova spectrometer.

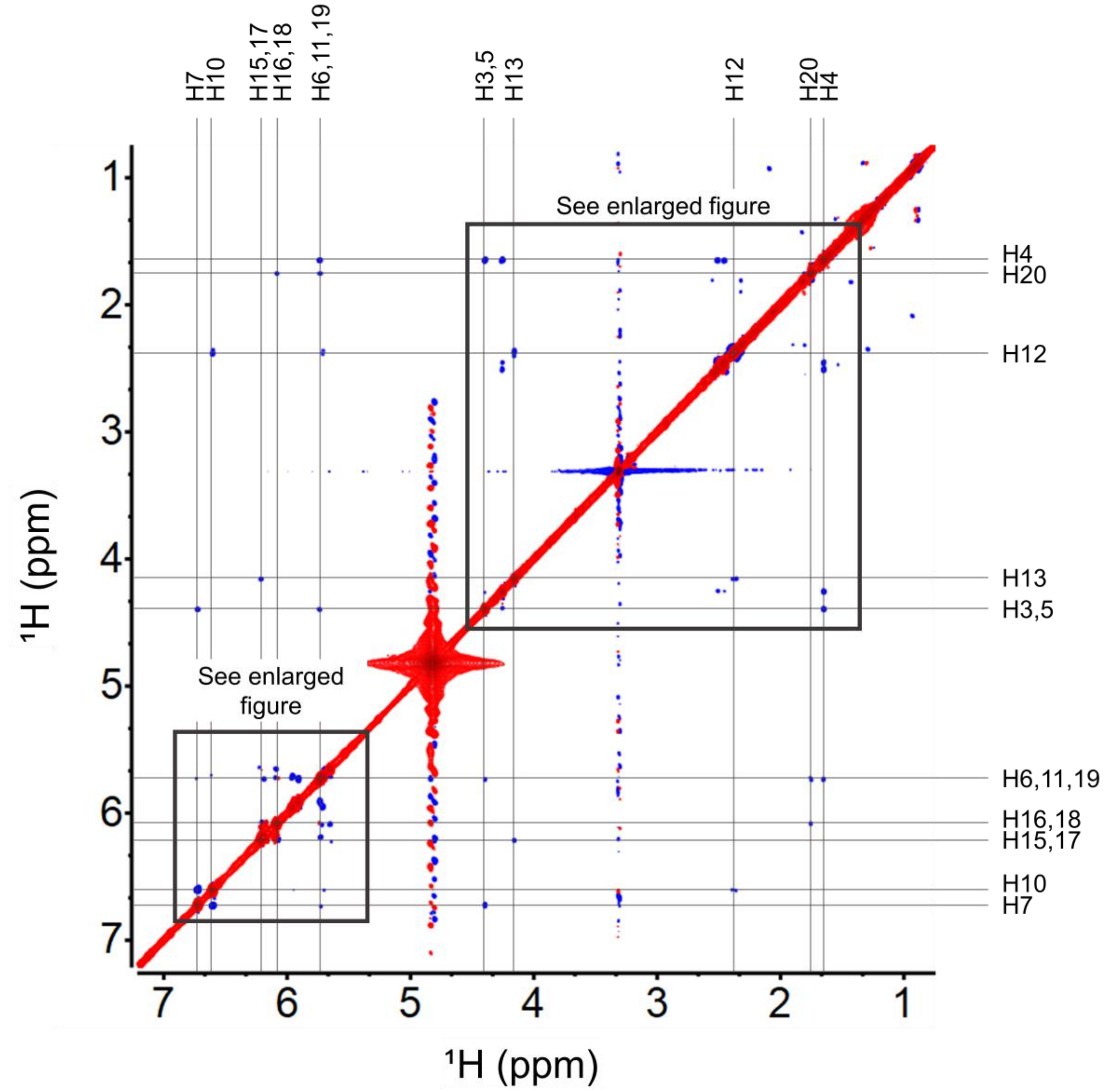

**Figure S26**. ¹H-¹H ROESY spectrum (positive signals in red; negative signals in blue; x-axis: δH, 7.2-0.7 ppm; y-axis: δH, 7.2-0.7 ppm) of **3** in $CD_3OD$. Spectrum acquired on a 900 MHz Bruker AVANCE II spectrometer.

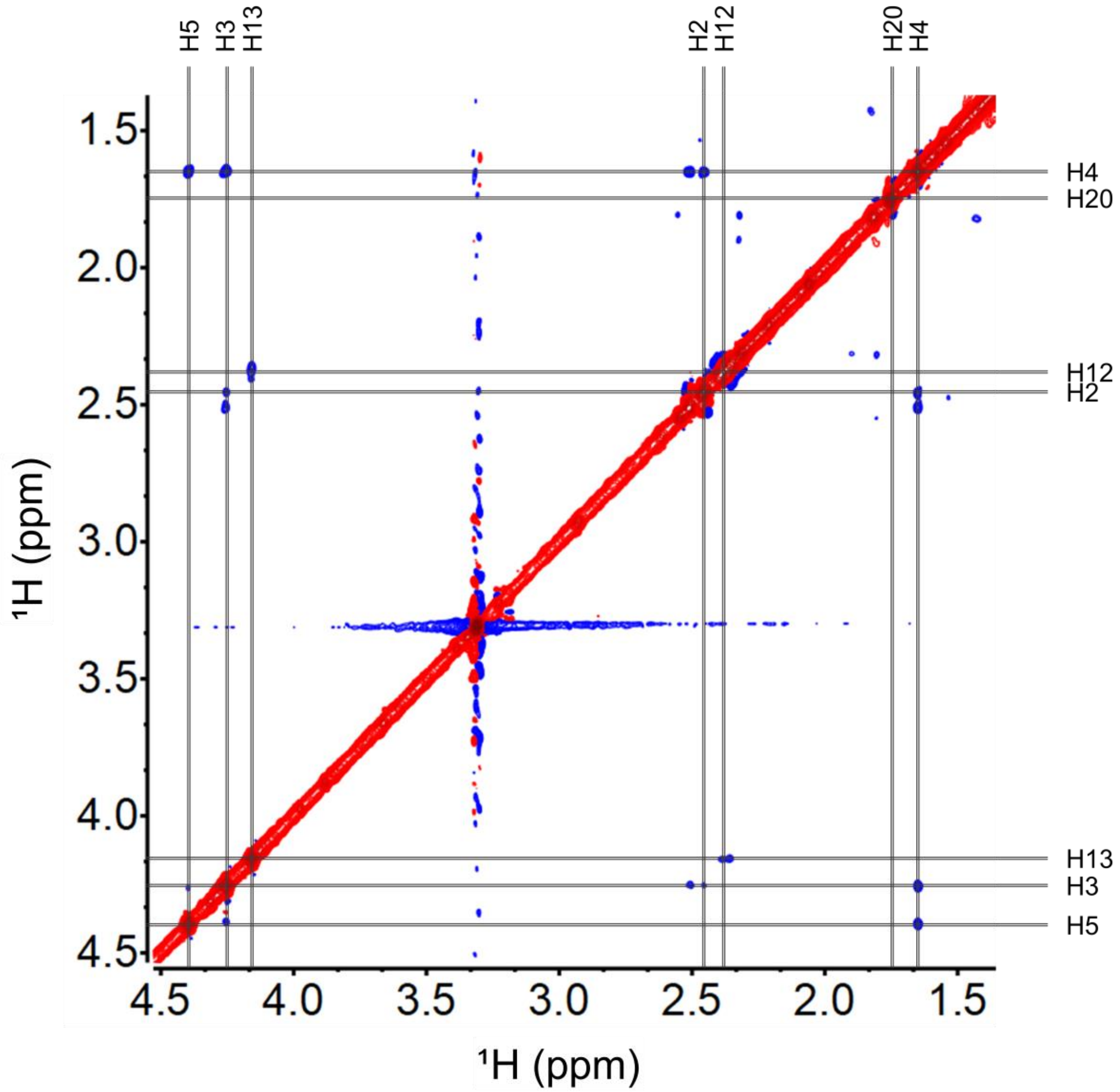

**Figure S27.** Enlarged region of $^1H$-$^1H$ ROESY spectrum (positive signals in red; negative signals in blue; x-axis: δH, 4.5-1.4 ppm; y-axis: δH, 4.5-1.4 ppm) of **3** in $CD_3OD$. Spectrum acquired on a 900 MHz Bruker AVANCE II spectrometer.

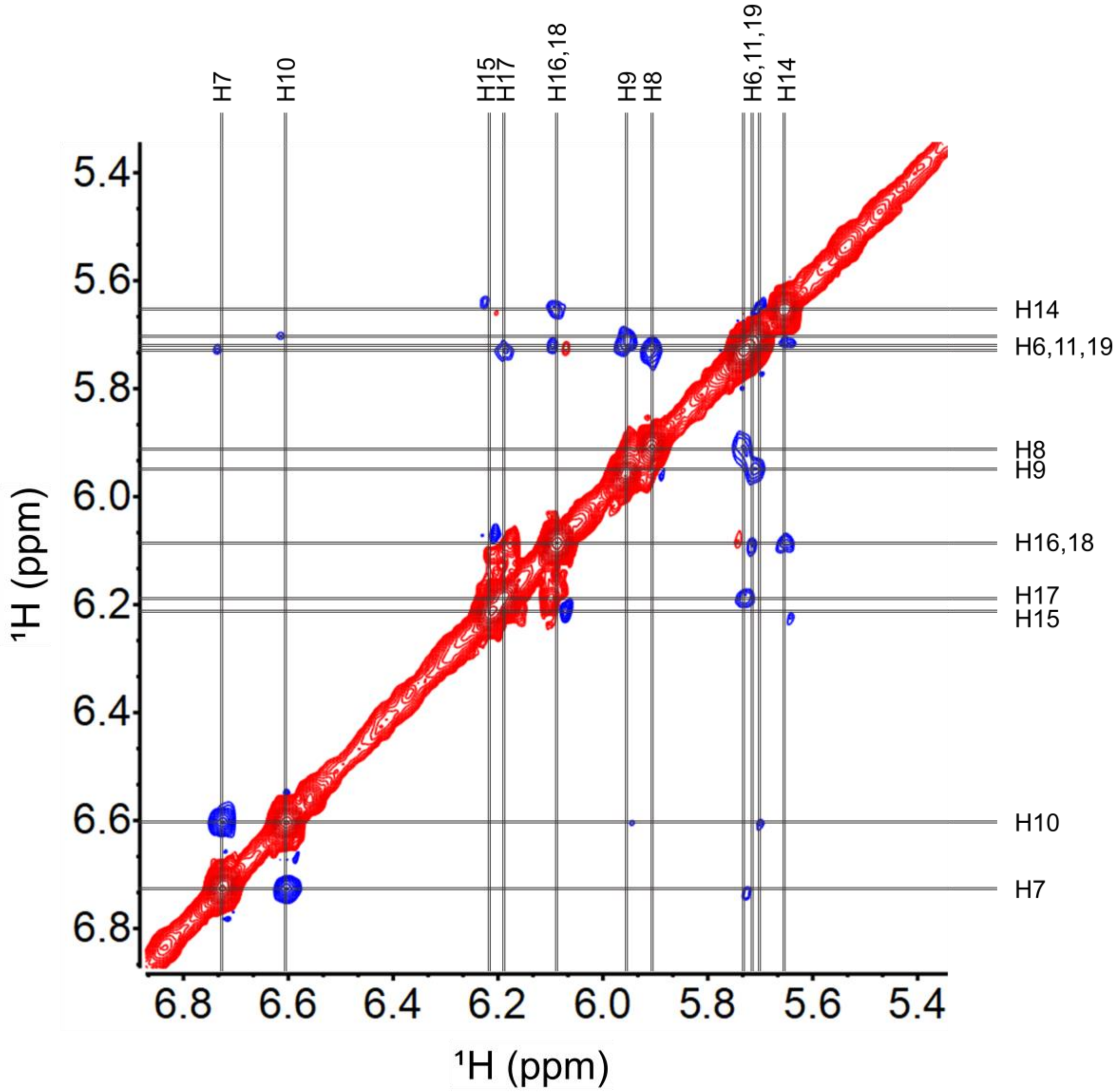

**Figure S28**. Enlarged region of $^{1}$H-$^{1}$H ROESY spectrum (positive signals in red; negative signals in blue; x-axis: δH, 6.9-5.4 ppm; y-axis: δH, 6.9-5.4 ppm) of **3** in $CD_3OD$. Spectrum acquired on a 900 MHz Bruker AVANCE II spectrometer.

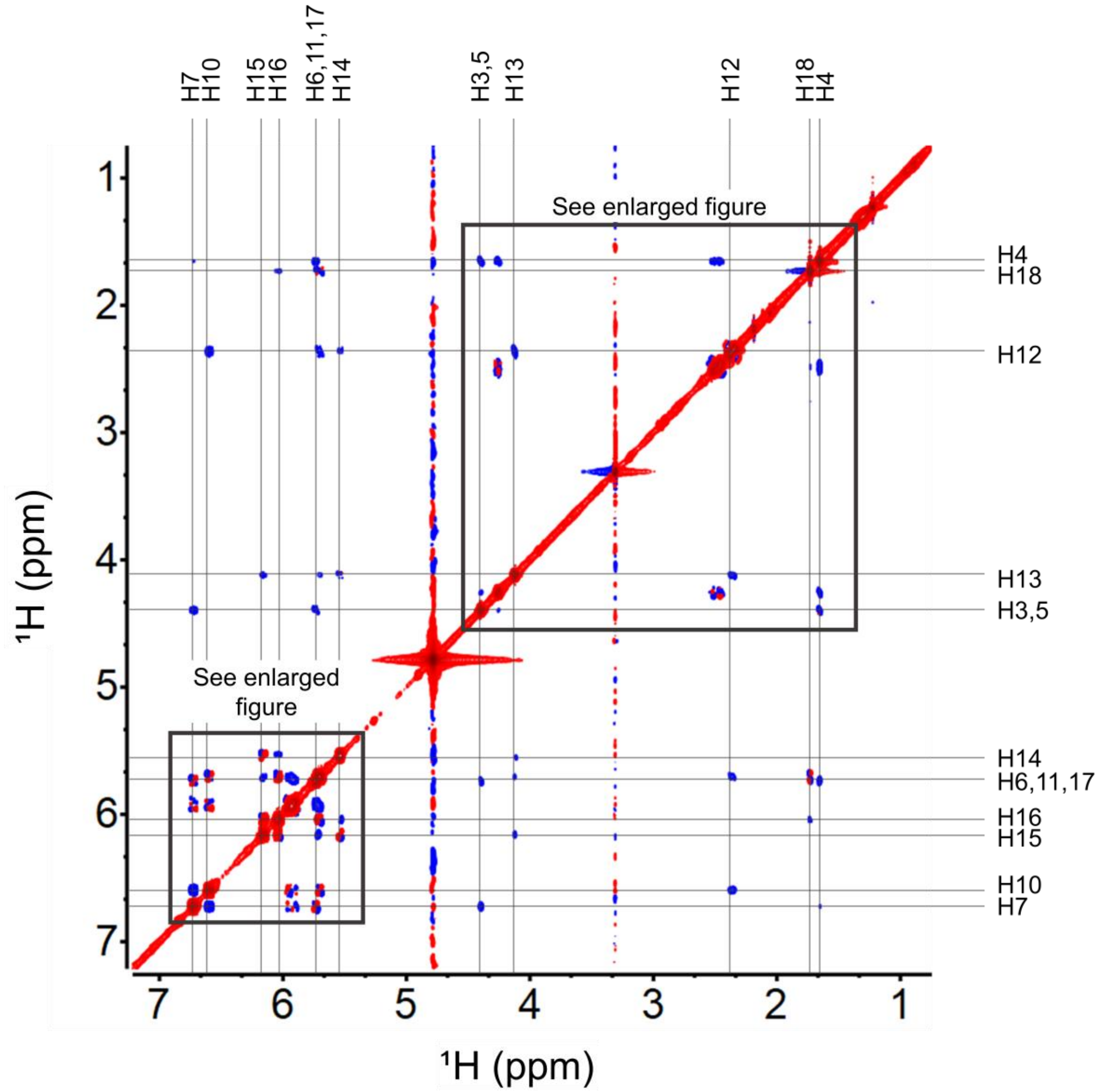

**Figure S29**. $^1H$-$^1H$ ROESY spectrum (positive signals in red; negative signals in blue; x-axis: δH, 7.2-0.7 ppm; y-axis: δH, 7.2-0.7 ppm) of **4** in $CD_3OD$. Spectrum acquired on a 600 MHz Varian Inova spectrometer.

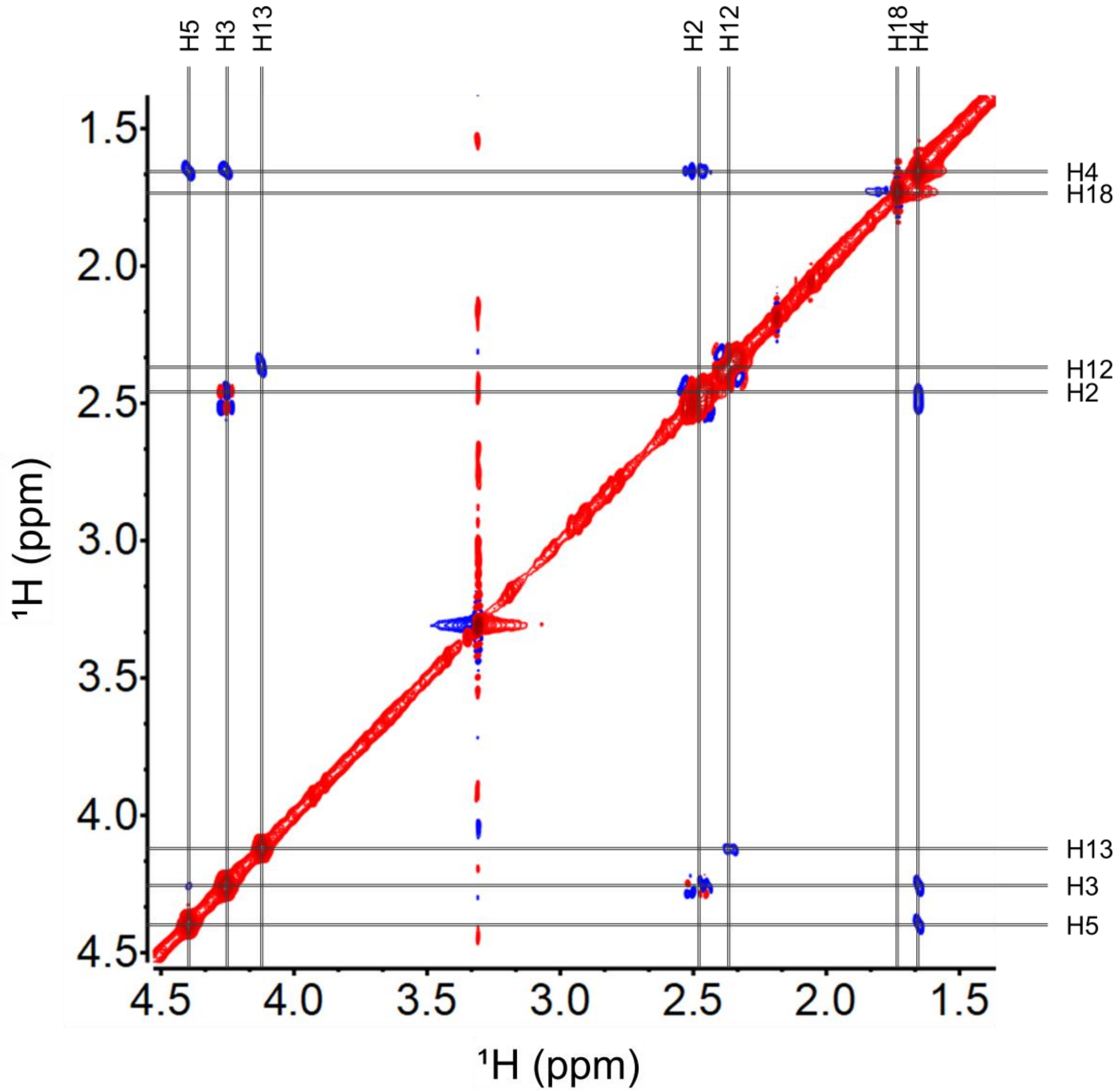

**Figure S30**. Enlarged region of $^1H$-$^1H$ ROESY spectrum (positive signals in red; negative signals in blue; x-axis: δH, 4.5-1.4 ppm; y-axis: δH, 4.5-1.4 ppm) of **4** in $CD_3OD$. Spectrum acquired on a 600 MHz Varian Inova spectrometer.

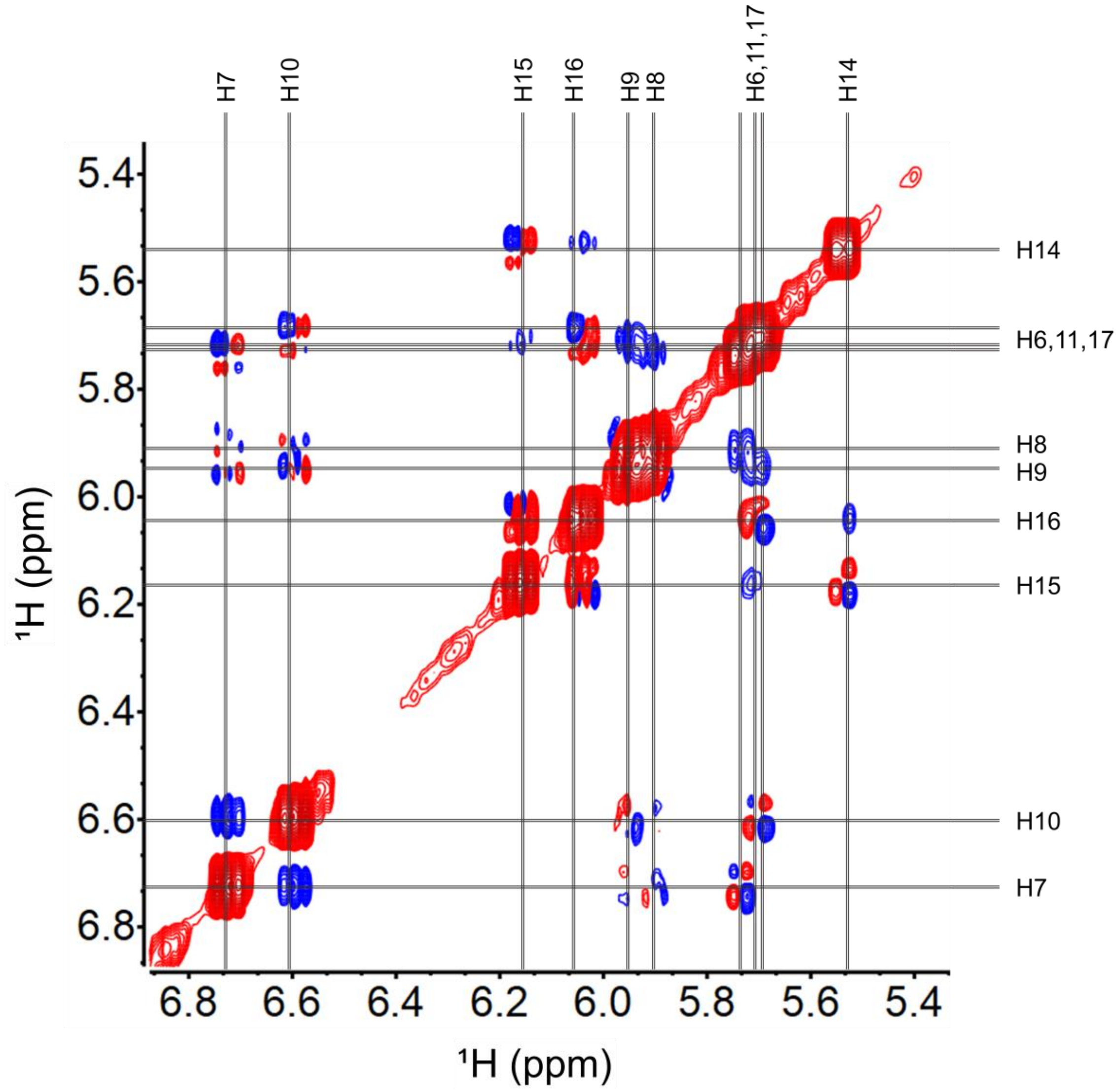

**Figure S31**. Enlarged region of $^{1}H$-$^{1}H$ ROESY spectrum (positive signals in red; negative signals in blue; x-axis: δH, 6.9-5.4 ppm; y-axis: δH, 6.9-5.4 ppm) of **4** in $CD_3OD$. Spectrum acquired on a 600 MHz Varian Inova spectrometer.

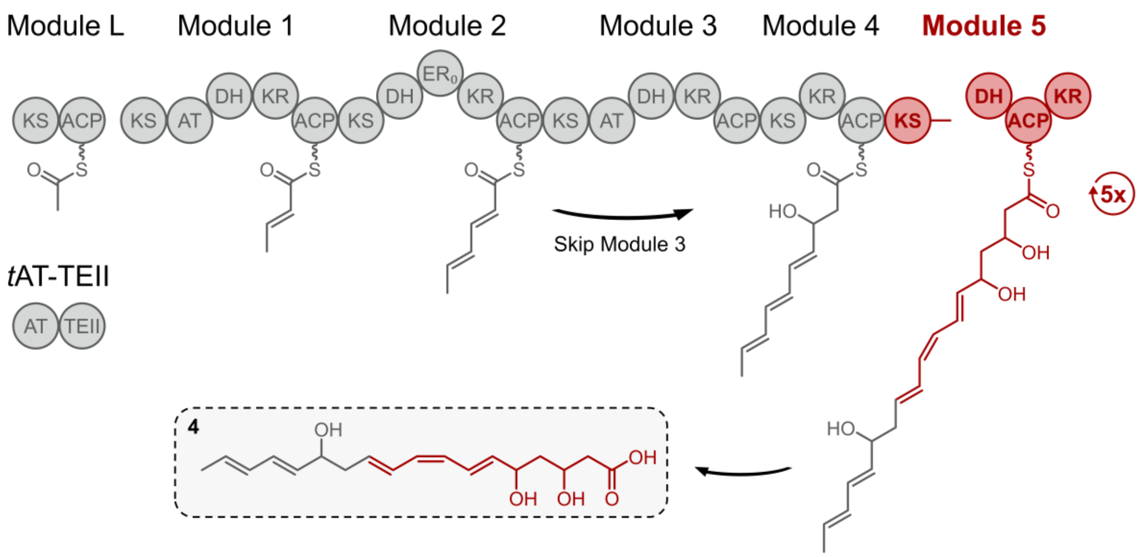

**Figure S32**. Biosynthesis of **4** from NOCAP synthase Modules L-5 + *t*AT-TEII.

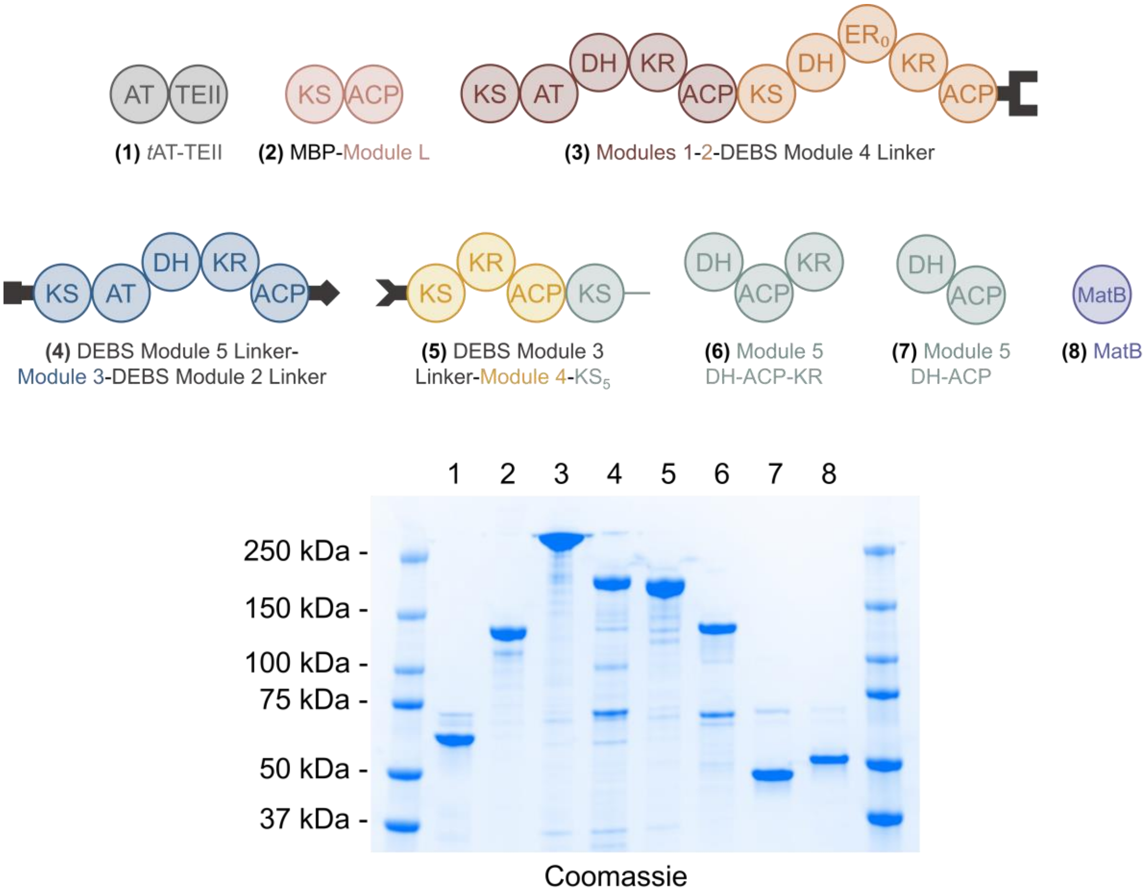

**Figure S33.** SDS-PAGE of purified proteins used for *in vitro* reactions. Left and right Markers: Precision Plus Protein Dual Color Standards (Bio-Rad Laboratories).

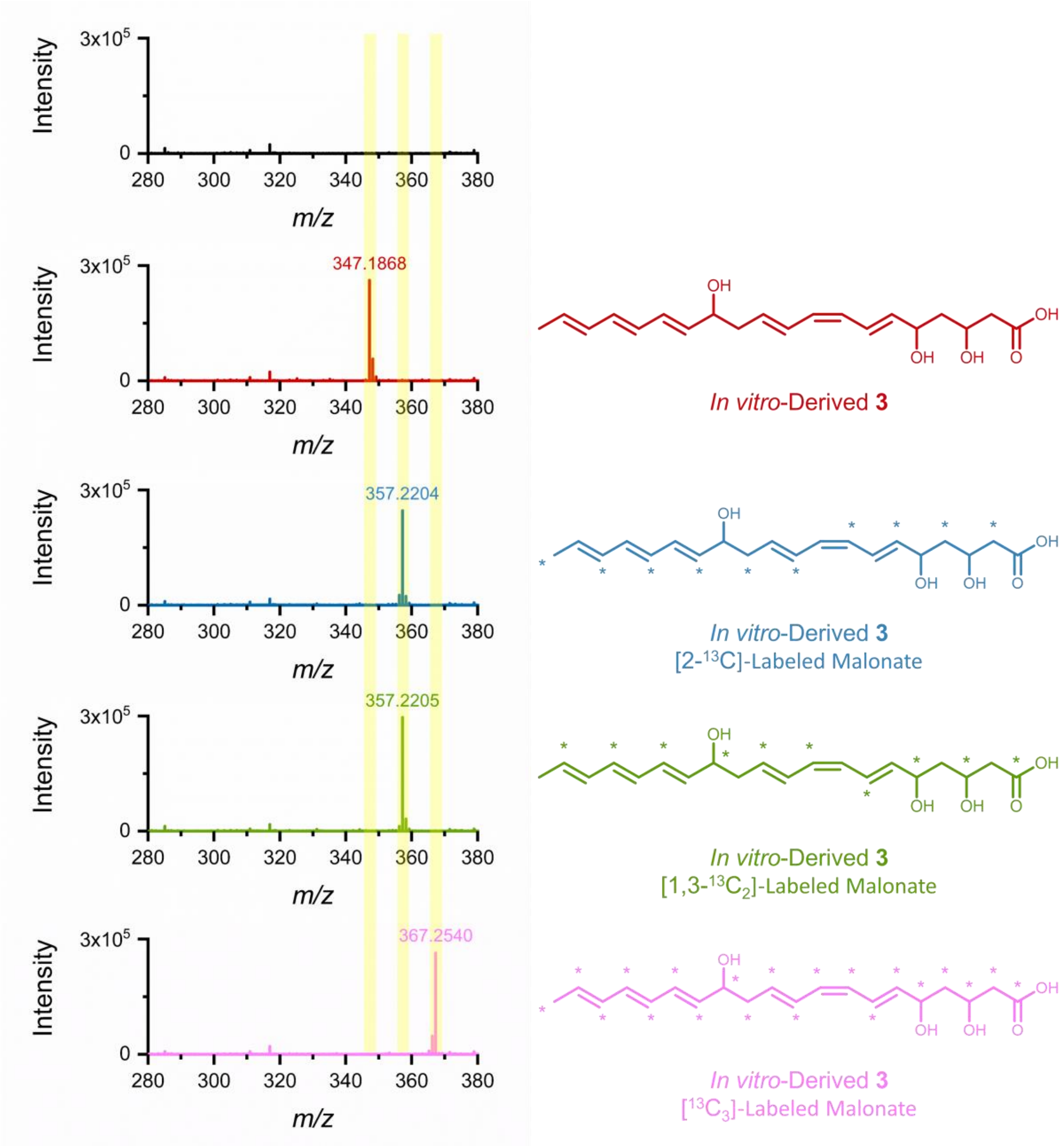

**Figure S34.** Mass spectra (ESI-) of *in vitro*-derived **3** from **no malonate** (black, top); **[$^{12}$C]-malonate** (red, 2nd from top, calcd. [M-H]$^-$ *m/z* 347.1864); **[2-$^{13}$C]-malonate** (blue, 3rd from top, calcd. [M-H]$^-$ *m/z* 357.2204); **[1,3-$^{13}C_2$]-malonate** (green, 4th from top, calcd. [M-H]$^-$ *m/z* 357.2204); and **[$^{13}C_3$]-malonate** (pink, bottom, calcd. [M-H]$^-$ *m/z* 367.2544). *, indicates $^{13}$C incorporation. Data were acquired on an Agilent Q-TOF 6545 LC-MS system and are representative of at least three experimental replicates. HRMS calculated values based on ChemDraw software predictions.
